# Standardized nuclear markers advance metazoan taxonomy

**DOI:** 10.1101/2021.05.07.443120

**Authors:** Lars Dietz, Jonas Eberle, Christoph Mayer, Sandra Kukowka, Claudia Bohacz, Hannes Baur, Marianne Espeland, Bernhard A. Huber, Carl Hutter, Ximo Mengual, Ralph S. Peters, Miguel Vences, Thomas Wesener, Keith Willmott, Bernhard Misof, Oliver Niehuis, Dirk Ahrens

## Abstract

Species are the fundamental units of life and their recognition is essential for science and society. DNA barcoding, the use of a single and often mitochondrial gene, has been increasingly employed as a universal approach for the identification of animal species. However, this approach faces several challenges. Here, we demonstrate with empirical data from a number of metazoan animal lineages that multiple nuclear-encoded markers, so called universal single-copy orthologs (USCOs) performs much better than the single barcode gene to discriminate closely related species. Overcoming the general shortcomings of mitochondrial DNA barcodes, USCOs also accurately assign samples to higher taxonomic levels. These loci thus provide a powerful and unifying framework for species delimitation which considerably improves the DNA-based inference of animal species.

## Introduction

Taxonomy, the science of naming, defining and classifying groups of organisms based on shared characters, faces new opportunities in terms of reproducibility, automation, and robustness due to major innovations in morphological and genomic analytical methods as well as continuously increasing computational power (*1–5*). Rapid and correct identification of species is paramount since knowledge and applications in conservation, medicine, and pest management among others are anchored to species, the fundamental entities of biodiversity. Conventional morphological taxonomic methods for species discrimination, in isolation, have been considered slow and inefficient to meet new challenges in the current era of biodiversity crisis (*6*). Furthermore, descriptions of new taxa based on morphology alone are more likely to require re-examination of types to assess diagnostic characters when new taxa are discovered in the future.

DNA-based approaches revolutionized the possibilities for resolving taxonomic questions that were formerly intractable or unfeasible through morphology-based approaches (*7*). During the past twenty years, DNA barcoding has increased the quality and reproducibility of species delimitation and identification but also enabled rapid assessments and monitoring of biodiversity (*8,9*). Its hallmark is the capability to standardize and automate species recognition by using a single specific and easily amplified gene fragment. In animals, the most widely used marker has been the mitochondrial protein-coding gene cytochrome oxidase subunit 1 (*COI*) (*7*). Beyond that, barcoding paved the way for direct inference of species boundaries from unknown samples (*10*). However, species delimitation and identification based on information from a single mitochondrial gene are prone to errors due to extrachromosomal inheritance, incomplete lineage sorting, sex-biased dispersal, asymmetrical introgression, or *Wolbachia-mediated* genetic sweeps (*11,12*). As a consequence, results from delimiting species by means of barcoding are not always congruent with those obtained from analyzing morphology or other data. Integrative taxonomic approaches have therefore been proposed to overcome these problems by complementing barcode-based species hypotheses with additional evidence (*13–16*).

Recent species delimitation approaches have considerably improved in accuracy by taking advantage of the phylogenetic information contained in multiple nuclear-encoded markers. These approaches mostly implement the multi-species coalescent model (*17–19*), and simulations have demonstrated their increasing accuracy and robustness with the analysis of more genes (*18,19*). In the past, the sampling of a small number of loci has been a compromise between costs and the accuracy of the inferred results (*20,21*). However, progress in DNA sequencing technologies promises a continuous decrease of DNA sequencing costs. Whole genome and transcriptome sequencing (*22–24*) or DNA target enrichment approaches enable the sampling of thousands of single copy target loci from an organism’s genome, even with degraded DNA, to help resolve questions that cannot be answered with data from only a limited number of loci (*5,6,25–27*).

Besides *COI*, different markers have been used for metazoan DNA taxonomy: nuclear ribosomal RNA genes (rDNA) (*28–30*); restriction site associated DNA sequences (RADseq) (*31–33*); and ultra-conserved elements (UCE) (*25,34*), including their more variable flanking DNA regions (*35,36*). However, they can hardly be applied universally across animals, either because of insufficient intraspecific variation or a lack of homologous loci (*21*). Therefore, universal single-copy orthologs (USCOs) have been proposed as a future core set of nuclear-encoded protein-coding genes for species delimitation in Metazoa (*21*). USCOs are proteincoding genes under strong selection to be present in a single copy (*37*). In Metazoa, 978 genes were classified as USCOs based on a representative selection of 65 high-quality genomes (*38*). The prerequisite for a gene to be an USCO is that it is present as single-copy in at least 90% of these genomes (*38*).

So far, USCOs have primarily been utilized for assessing the completeness and quality of sequenced genomes and transcriptomes (*38*). However, they are also promising markers to establish a universally applicable genomic species identification and delimitation procedure (*21,24*). Since the number of single-copy genes increases with increasing relatedness of the species under consideration (*39*), sets of USCOs are larger for taxonomic levels closer to the tips of the tree of life (*21*). Yet, there is evidence that USCOs allow inferring the wider systematic placement (e.g., class, order, or family) of an unknown sample (*21*), despite the fact that the number of single-copy genes decreases with increasing taxonomic rank. Since the amino acid sequence of protein-coding genes is typically more conserved than the underlying coding nucleotides, it permits the identification of even highly diverged orthologous DNA sequences. Protein-coding gene regions can be analyzed on both the transcriptional (nucleotide) and translational (amino acid) level and thus allow a more reliable assessment of homology. These qualities strengthen the premise that USCOs are a highly suitable marker system as a universal tool in DNA taxonomy.

Here we demonstrate the suitability of USCOs for species identification within Metazoa using empirical genomic data. In nine study cases (table S1), we sequenced USCOs for different genera from various arthropod (Chelicerata, Myriapoda, Coleoptera, Diptera, Hymenoptera, and Lepidoptera) and vertebrate (Amphibia) lineages. These taxa include morphologically well-defined species, which we refer to as morphospecies, which are closely related and in some cases cannot be distinguished by means of *COI* barcodes. Validation of USCOs as suitable and superior markers for species delimitation requires (a) sufficient overlap in the recovery of USCOs between different groups of Metazoa to allow comparison and large-scale analyses of even distantly related species and groups (for higher level assignment), (b) robust results by clearly specified wet lab and bioinformatic protocols that recover data from a high proportion of samples, (c) sufficient phylogenetic resolution to separate closely related species, and (d) agreement of resulting groupings with morphospecies (and/or alternative evidence for robust species hypotheses such as hybrid zone analysis).

## Results

### Data recovery success

We obtained sequences by DNA target enrichment comprising up to 950 USCOs with more than 0.5 million base pairs per study case (table S2). USCOs were subsequently assessed for their discriminative power in inferring phylogenetic relationships and species boundaries, using current state-of-the-art species delimitation approaches within selected case study taxa. The success of data recovery by seven different DNA sequence assembly approaches (A1-A7) (figs S1, S2) varied considerably between approaches and taxa (Fig. 1; fig. S2; tables S2, S3). The best performing assembly approaches yielded at least 700 USCOs per case study and more than 200,000 base pairs. With these approaches, most USCOs were recovered in the majority of the nine genera studied (Fig. 1) and frequently in all or almost all individuals. Sometimes USCOs were recovered in only a few (*1-3*) individuals, in some assembly approaches more frequently than in others (fig. S2). These USCOs were excluded to avoid an excess of missing data (see Supplementary Materials).

**Fig. 1.**
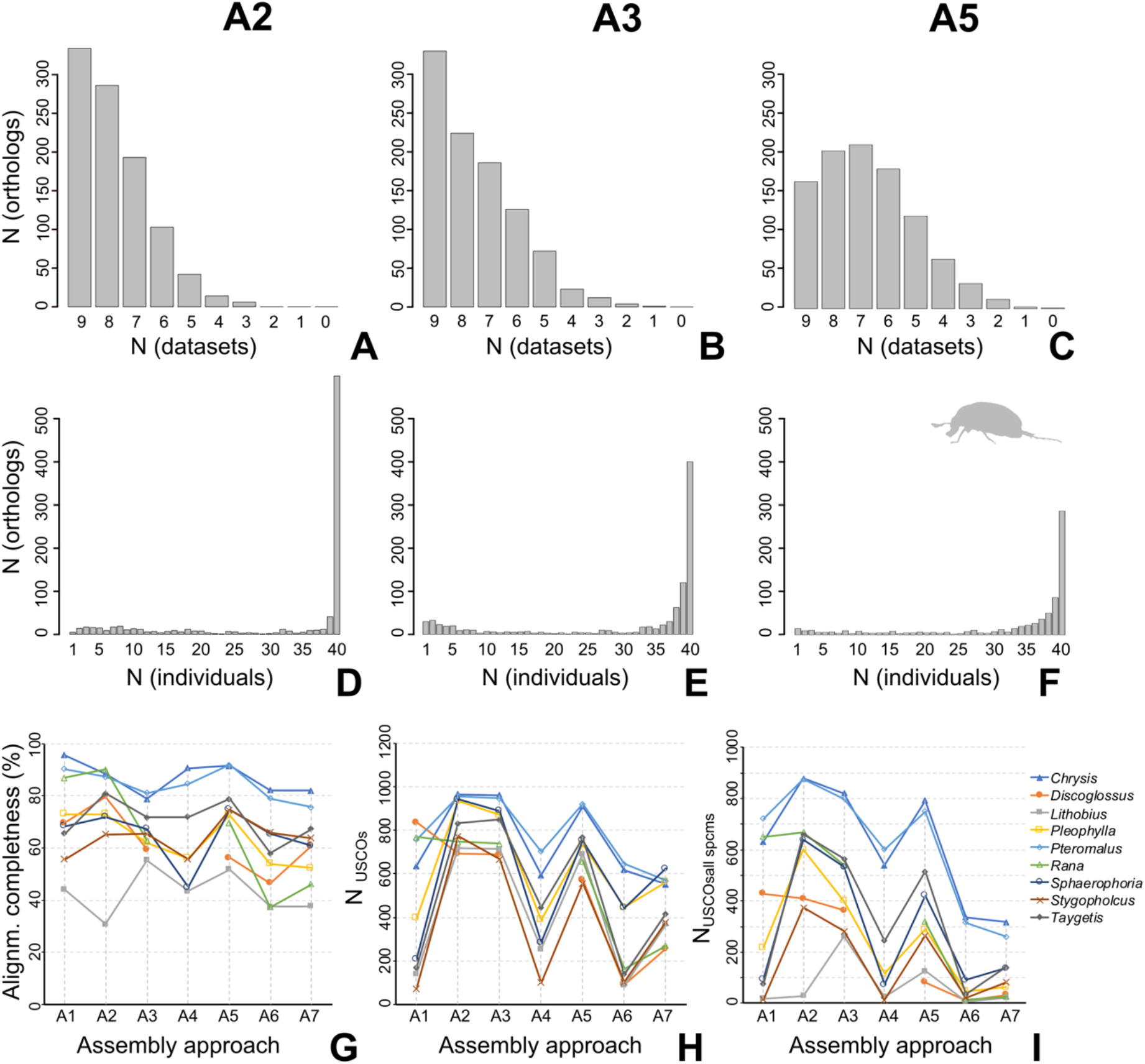
Evidence for the universal applicability of USCOs as taxonomic markers. **(A-C)** Number of orthologs present in different numbers of data sets of the nine study cases in three of the seven assembly approaches (A2, A3, A5). **(B-F)** Distribution of orthologs over the number of specimens within one case study in three of the seven assembly approaches (A2, A3, A5) (*Pleophylla*). **(G)** Completeness of concatenated alignments, **(H)** number of USCOs and **(I)** number of USCOs present in all specimens of each case study in each assembly approach.

The variation in recovery success between assembly approaches (Fig. 1; fig. S2) can be ascribed 1) to the varying degree of relatedness and thus similarity of the reference DNA sequences available for each case study for bait design, assembly (if applicable) and loci selection, but also 2) to methodological differences in the assembly pipelines. Outlier filtering removed only a very small amount of data, never more than 1%. The ratio of data yield between worst and best assembly ranged from 0.6 to 0.09 for the number of USCOs and 0.47 to 0.08 for the number of base pairs. Completeness of the resulting alignments varied more by study group than by assembly method. It was especially high in Hymenoptera (78.7 to 96.5 % in *Chrysis*, 75.6 to 91.2 % in *Pteromalus*), and particularly low in *Lithobius* (31.6 to 55.3 %). As an example of the differences between approaches, A2 usually resulted in greater numbers of recovered genes than A1, because it used reference sequences from within the examined genus, while the sequences used by A1 were often from other genera within the same family (table S3). Therefore, it seems that the reference taxa used were often not sufficiently closely related to the target taxa for A1 to reliably identify homologous genes with *bwa* (*40*). This approach required a seed region length of 30 successive identical base pairs to the reference DNA sequence, which was apparently not present for many genes. In the myriapods (*Lithobius*), this is the case also for A2, as many USCOs were found only in the morphospecies from which the respective reference sequence originated, thus explaining the low recovery success for this assembly method. Each *Lithobius* species was represented by five individuals, thus highest USCO recovery resulted for five specimens, while overall recovery of USCOs for this genus was remarkably low (Fig. S2, table S2). Due to the methodology of A2, reference sequences for different loci came from all included morphospecies. In *Lithobius*, the four tested species were separated by very long branches in the phylogenetic tree, indicating a much higher amount of interspecific divergence than in the other study cases. We refrained from using a shorter seed region length, since this increases the chance of false hits and including paralog genes. Our results partly mirror findings from previous studies comparing different assembly approaches for data extraction from hybrid enrichment data (*41*), which found that *Hybpiper* (*42*) recovers a larger number of loci compared to *Phyluce* (*43*), but the latter produces longer per-locus alignments as the reads are assembled to contigs prior to mapping, resulting in the inclusion of sequence regions without high similarity to the reference.

Visual inspection of the alignments also showed that the approaches using *bwa* mapping (A1 and A2), where only highly similar regions are aligned, produce data of the highest quality and this does not result in a lower amount of data compared to other approaches. This is also reflected, particularly in A2, by the completeness of the alignment and the number of USCOs present in all specimens (Fig.1; table S2). An additional advantage of A1 and A2 is that diploid consensus sequences can be generated (while A3-A7 produced haploid consensus sequences), allowing investigations of heterozygosity and SNP-based analyses. Using A1 (direct mapping with bwa against a reference) seems preferable if a relatively complete reference genome or transcriptome is available from within the studied group, while in other cases USCO sequences can be extracted from the DNA target enrichment data themselves (e.g. with *Orthograph* (*44*) for use as reference for further mapping with *bwa*, as in A2.

### Phylogenetic analyses

Species inference under the phylogenetic and biological species concepts (*18,45*) with USCO data relies on accurate phylogenetic hypotheses. In the phylogenies inferred using coalescent-based species tree approaches and maximum likelihood analysis of concatenated data, 98 % of the morphospecies in the study cases resulted as monophyletic (Fig. 2; figs S3-8). All trees had robust and well-resolved interspecific topologies that widely agreed among tree reconstruction methods and assembly approaches (A1-A7), while intraspecific relationships often differed in topology among different assemblies. Morphospecies were sometimes not separated in our *COI* benchmarking analyses (see below) but were almost always recovered as monophyletic groups by USCOs (Fig. 2, fig. S6). Just one case showed a disagreement between USCO tree topology and morphology-based taxonomy: the beetle *Pleophylla fasciatipennis* was split into two separate independent clades, possibly reflecting the presence of true cryptic species. The universality of the approach is demonstrated by the ability to present a meaningful tree (Fig. 2A; fig. S9) resulting from a concatenated super-alignment of all target groups as well as the reference taxa (table S3). All genera and higher-level systematic groups (e.g., orders), as well as most of the morphospecies, were monophyletic, with well-resolved internal relationships.

**Fig. 2.**
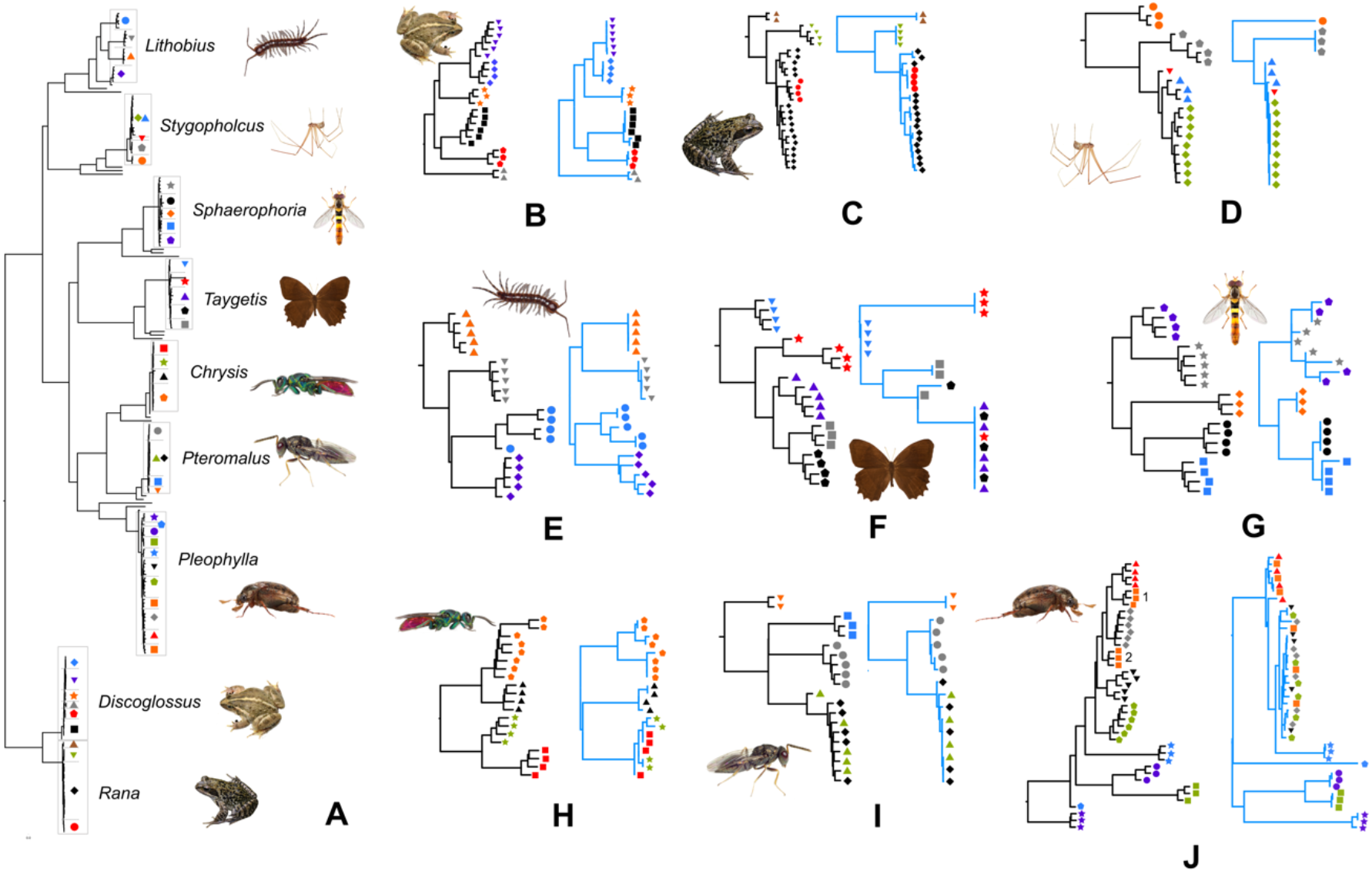
Phylogenetic resolution of USCOs. **(A)** Tree computed with Maximum Likelihood from concatenated USCOs of all study cases (A3; all nucleotides); branches of reference taxa (Supplementary Information) are unlabeled. (**B–J**) USCO trees (black) obtained from ASTRAL analysis of USCOs (A2) of individual study cases compared with the *COI* benchmarking tree (blue). Study cases: *Discoglossus* (**B**), *Rana* (**C**), *Stygopholcus* (**D**), *Lithobius* (**E**), *Taygetis* (**F**), *Sphaerophoria* (**G**), *Chrysis* (**H**), *Pteromalus* (**I**), *Pleophylla* (**J**). Morphospecies indicated by colored symbols.

### Phenetic analyses with SNPs

Single-nucleotide polymorphisms (SNPs) extracted from the USCOs obtained with assembly approach 2 (A2) varied considerably in number among the different study cases due to different numbers of recovered USCOs, ranging from 1,950 (*Lithobius*) to 17,364 (*Pleophylla*). Multidimensional scaling analyses on SNPs (NMDS; Fig. 3) showed nearly all morphospecies as discrete clusters. In the diphyletic morphospecies *Pleophylla fasciatipennis*, the two separate clades were also recovered by NMDS as obviously distinct from each other and from other species. However, the visibility of this outcome is dependent on scaling, since the different species included in some cases differ strikingly in the amount of genetic divergence (Fig. 3). Therefore, closely related species often group very closely together in the NMDS plots, meaning that it is not immediately apparent whether they still form distinct clusters. However, after removing more distantly related taxa from the NMDS analysis, even closely related morphospecies can easily be seen to form distinct clusters. This shows that all results need to be interpreted carefully and counterchecked, ideally also with other lines of evidence. In one case, individuals of a clearly monophyletic morphospecies (*Lithobius crassipes*) did not form a distinct cluster, probably due to the low data recovery in the group and strong infraspecific divergence in this species (see supplement file).

**Fig. 3.**
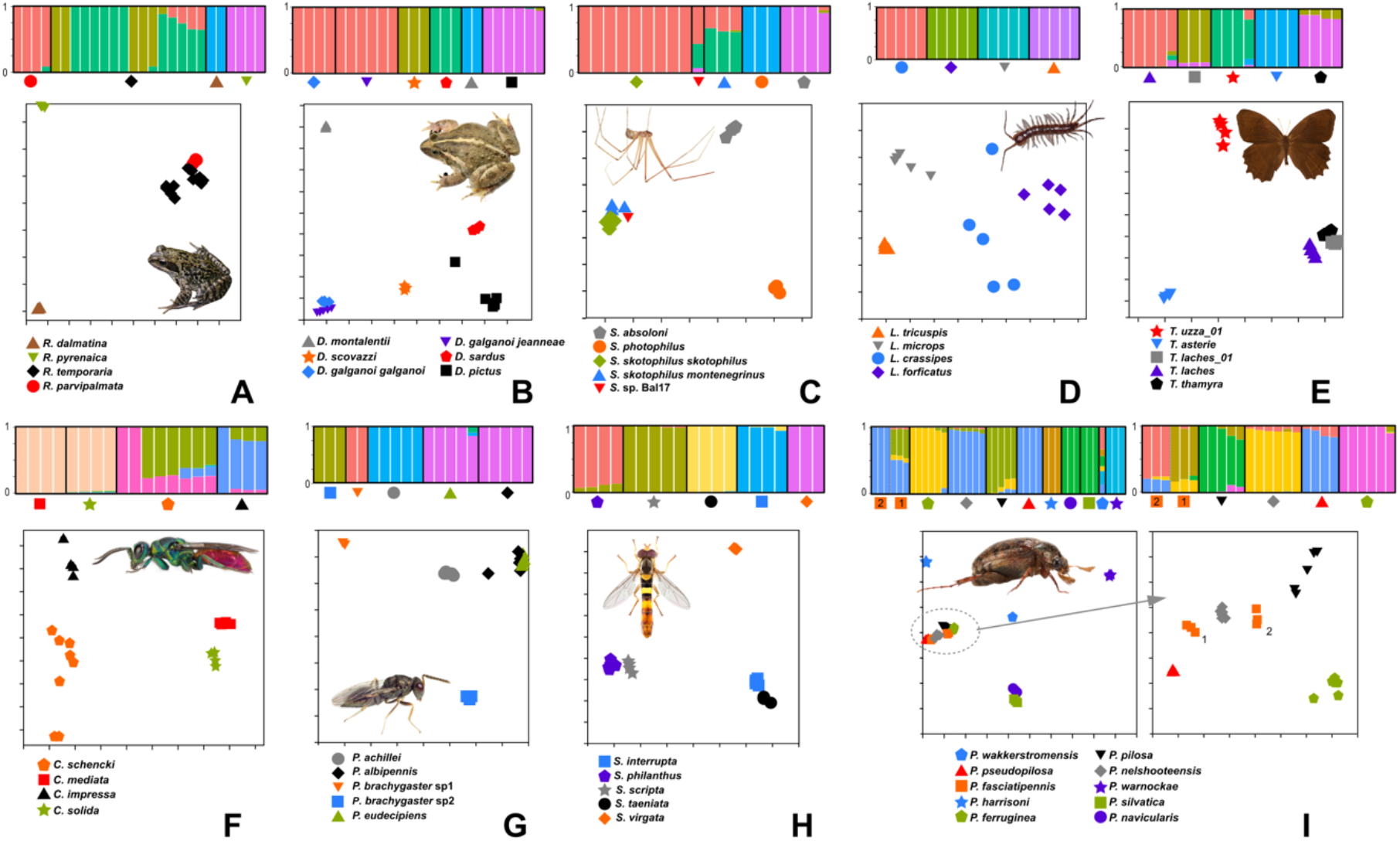
Discriminative power of USCOs. **(A–I)** Probabilities of cluster assignment from STRUCTURE analysis (above) and plots from NMDS (below) on SNPs derived from USCO data. Study cases: *Rana* (**A**), *Discoglossus* (**B**), *Stygopholcus* (**C**), *Lithobius* (**D**), *Taygetis* (**E**), *Chrysis* (**F**), *Pteromalus* (**G**), *Sphaerophoria* (**H**), *Pleophylla* (**I**). (**I**) shows the results of analyses of all species, and a subset (*P. pilosa* group (encircled in left NMDS plot). Morphospecies indicated by colored symbols.

Population admixture analyses with *STRUCTURE* (*46*) confirmed the monophyletic morphospecies in most cases while further subsplitting was not evident (Fig. 3). Taking only the results with the highest likelihood into account, individuals were assigned to the clusters corresponding to species with at least 90% probability. In all our study cases, increasing K beyond a certain number, usually corresponding to the number of morphospecies, had no effect on the results, as no individuals were assigned to the additional populations. However, the MCMC frequently stabilized in a local maximum where certain distinct species were not distinguished. Such phenetic analyses may therefore currently be an alternative method of species delimitation with a large amount of data, although the analyses have to be repeated several times to avoid local maxima. This is especially the case when certain species are very closely related to each other, as in the case of *Taygetis thamyra* and *T. laches_01*, or of *Pleophylla pseudopilosa* and *P. fasciatipennis* (syntopical population). *Chrysis* showed another source of misleading results, where the females of *C. impressa* and *C. schencki*, which are diploid and to some degree heterozygous, were inferred to have a large degree of admixture with each other, while the males, which are haploid as in all Hymenoptera, did not. Lack of heterozygosity in males also caused longer terminal branches for males in the concatenation-based phylogenetic trees for analyses based on diploid data, i.e. those using *bwa* mapping (A1 and A2) (fig. S13).

Morphologically cryptic species with narrow contact zones and almost without admixture, such as the frogs *Discoglossus pictus* and *D. scovazzi*, were resolved as distinct by the clustering approaches, whereas lineages known to admix over wide hybrid zones, such as the two subspecies of *D. galganoi* (*47*), were correctly grouped into a single species-level unit. The European common frog (*Rana temporaria* complex) is partitioned into three distinct entities by population admixture analyses, which are also apparent in the phylogenetic trees of the more complete data assemblies. One of these corresponds to the taxon *R. parvipalmata* which was recently recognized as a distinct species due to the very narrow hybrid zone separating it from *R. temporaria* (s. str.) (*48*). This congruence with previous in-depth studies validates the USCO-based analyses. It is remarkable that the USCO data suggest the possibility of at least one further, previously unrecognized species-level split between Eastern and Western European populations. This calls for further in-depth study and highlights that even in the European common frog – one of the most prominent and well-studied amphibians thought to occur from Spain to Britain and northern Norway, and into much of Siberia, taxonomic surprises can be expected.

Admixture between species was generally only found if the species were very closely related to each other. In the case of *Pleophylla fasciatipennis*, where the morphospecies was not recovered as monophyletic, no admixture between the two different clades was detected, at least at sufficiently scaled analysis, suggesting that the morphological similarity between those lineages cannot be explained by hybridization.

### Species delimitation analyses

The overall outcome of multi-species coalescent analyses with parametric (*BPP*) (*49*) and non-parametric methods (*tr2*) (*50*) suggested that morphospecies were split into additional entities. Only in one case two morphospecies were lumped (*Pteromalus eudecipiens* and *P. albipennis*), probably due to the taxonomic misinterpretation of one of the species (a final decision, however, requires a thorough morphology-based taxonomic revision of types and reexamination of the sequenced specimens). As expected, the species delimitation analyses using Bayesian Phylogenetics and Phylogeography (*BPP*) showed great influence of the choice of priors for theta and tau on the results (figs S16-S30, see Supplementary Materials). For *BPP* analyses the criterion for species delimitation was the median posterior probability, computed from five independent runs of each analysis. The median was computed for each of the nine predefined prior combinations of population size (theta) and divergence time (tau) (Fig. 4; figs S10-S15). Over-splitting was sometimes observed with the full USCO data set at all levels of infraspecific nodes (even in syntopic specimens) (figs S13-S15). However, after excluding sites containing missing or ambiguous (heterozygote) data, over-splitting was reduced and mostly geographically separated populations were split (Fig. 4; figs S10-12, S16-S24).

**Fig. 4.**
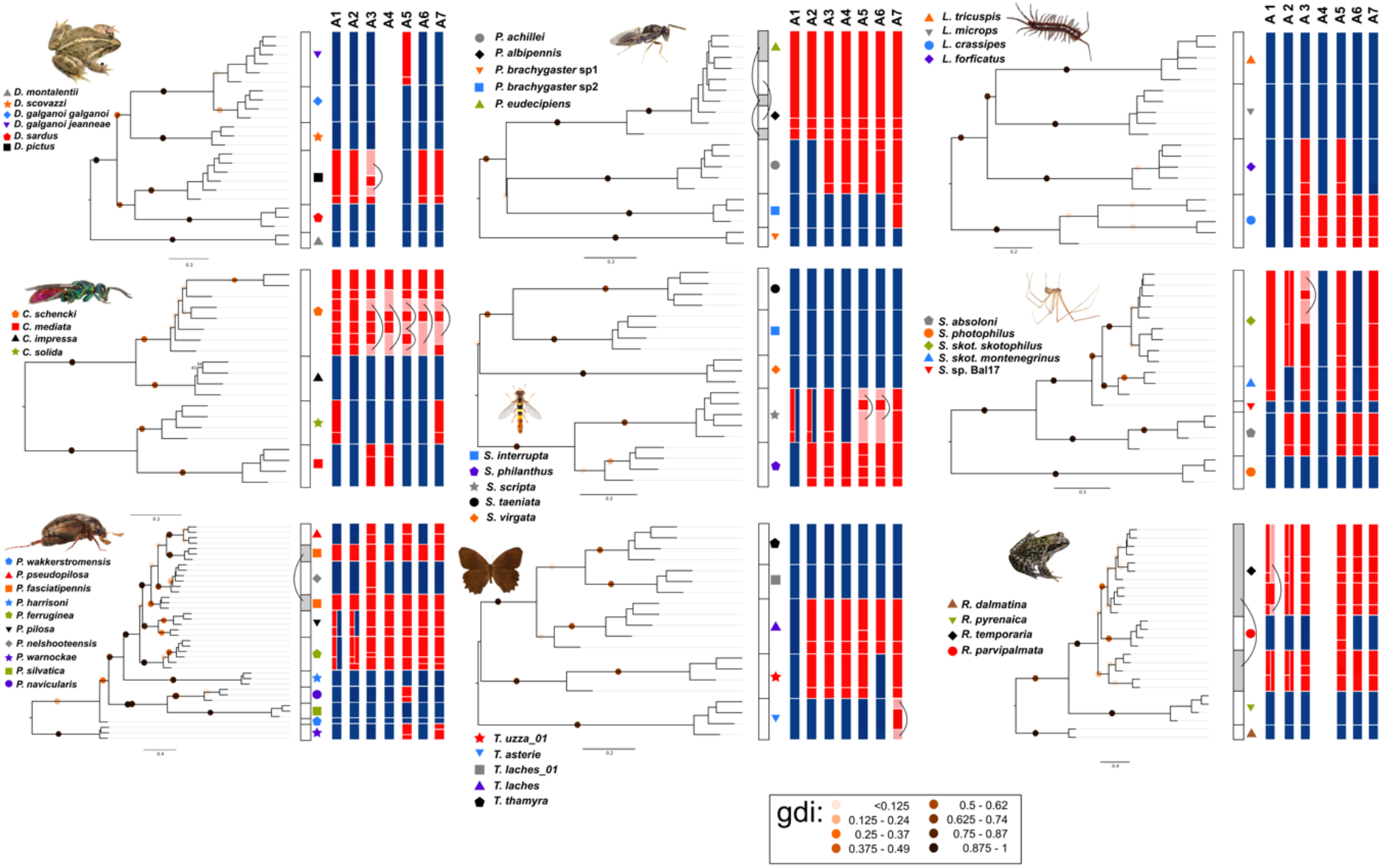
Results of *BPP* species delimitation based on reduced USCO data* of each case study and assembly approach (A1-A7) are mapped onto ASTRAL trees (inferred from all data of A2) and compared to morphospecies assignments (bars and colored symbols next to terminals). *BPP* results are based on the median of posterior probabilities of all nine prior combinations. Gdi values are mapped onto branches. Squares in columns indicate inferred species entities (white: morphospecies; grey: non-monophyletic morphospecies; blue: concordant with morphospecies; red: incongruent with morphospecies; pink: entity not reflected by shown tree topology and linked by a bracket, incongruent with morphospecies. Species entities from different assembly approaches may not be monophyletic in this tree because alternative assembly approaches may result in differing guide tree topologies). * Sites with gaps were removed; for the right column in A1 and A2, also sites with ambiguous data were removed.

Using all USCO data of *BPP* analyses, the genealogical divergence index (gdi) (*51,52*) proved not to be a suitable general proxy to evaluate species status and to refute over-splitting (figs S13-S15). For many of the splits, the index value fell between the established inter- and intraspecific gdi thresholds, assuming gradual values according to the degree of divergence of the examined lineages. Many of these splits with intermediate gdi values were not only well differentiated morphologically and genetically but also occurred syntopically (tab S5). While some species, especially those with long branches in the phylogenetic trees, such as *Pleophylla harrisoni* and *Stygopholcus photophilus*, were clearly supported as distinct (gdi > 0.7) and in other cases splits within morphospecies could clearly be rejected (gdi < 0.2, e.g. within *Pleophylla nelshoogteensis, P. pseudopilosa* and *Sphaerophoria scripta*), for most groupings gdi was between 0.2 and 0.7, making their species status ambiguous.

With reduced USCO data, obtained after excluding missing or ambiguous nucleotides the gdi values at nodes matching morphospecies boundaries were all well above the established inter- and intraspecific gdi thresholds (i.e., 0.7) (*52*), with very few exceptions in *Sphaerophoria philanthus* and *Taygetis laches* (Fig. 4). Gdi values were then distinctly below the threshold in the few available infraspecific nodes.

### COI benchmarking

In only two of the nine study cases (*Discoglossus, Lithobius*) the ML tree based on *COI* data recovered all morphospecies as monophyletic groups (Fig. 2; fig. S31). In one additional case (*Pteromalus*) two potential morphospecies could not be distinguished which also were not separated with the USCO data. The worst performance of *COI* occurred in *Pleophylla*, where five morphospecies easily distinguished by morphology (*53,54*) could not be resolved as monophyletic. While three *Taygetis* morphospecies had identical haplotypes, in *Lithobius* two morphospecies showed very deep coalescence, leading to problems with species delimitation. Furthermore, most species delimitation approaches with *COI* also yielded over-splitting in most of the study cases (fig. S31). Results of the different species delimitation methods were partly quite divergent within each study case, particularly in *Taygetis* and *Sphaerophoria*, in which some methods produced just one MOTU (parsimony network analysis), others up to 17 and 16, respectively (*bPTP*). The most consistent results again were obtained with *Discoglossus*, were in *mPTP* there was a 100% correspondence between morphospecies and MOTUs.

## Discussion

In our study we used empirical evidence to evaluate the feasibility of a unified standardized marker system using nuclear-encoded genomic data using USCOs. The fact that alignments based on all assembly approaches as well as different tree reconstructions methods (concatenation-based vs. coalescent analyses) resulted in very similar trees can be seen as evidence for the robustness of USCO data in relation to variation of alignment completeness (*ASTRAL* (*55*): figs S3-S5; *IQ-TREE* (*56*): figs S6-S8). More thorough concatenation-based *IQ-TREE* analyses (parameter: –m MFP+MERGE, 50 times repeated) resulted in only slight changes of topology for poorly-resolved intraspecific nodes compared to the “explorative” runs, showing that the tree topology of the infraspecific nodes and nodes around species-level do not depend greatly on parameter choice of tree searches. Even in cases with a large amount of missing data, or general poor data recovery, USCOs yielded enough information to generate mostly well-resolved (at species level) and reliable phylogenies. Phylogenetic analyses with reduced data (ambiguous and/or gaps omitted) showed very similar tree topologies (figs S22-S27), except for three assembly approaches in *Chrysis* and one in *Discoglossus* (fig. S27) where monophyly of one morphospecies failed to be recovered. Altogether, this shows that USCOs have the potential to reliably differentiate between species and obtain well-resolved phylogenetic trees even in cases where taxa are not distinguished with *COI* (seven out of nine study cases). Beside the striking congruence between USCO phylogenies and species delimitations based on morphology, our results also show the potential to resolve open taxonomic questions and detect unrecognized, truly cryptic species. In *Pleophylla*, USCO data suggest that *P. fasciatipennis* consists of two geographically separated lineages that are not each other’s closest relatives and that might represent separate species, even though they are extremely similar morphologically (*53,54*). *STRUCTURE* clustering allowed two alternative interpretations, depending on the scaling of the analysis: results of the sampling including all *Pleophylla* species suggest that in one clade of *P. fasciatipennis* there was a large amount of hybridization with the syntopic *P. pseudopilosa*. However, with a systematically narrower sampling of only closely related lineages (*P. pilosa* clade), the existence of two separate and well-distinct genetic lineages (or species) within *P. fasciatipennis* is supported (Fig. 3I). In the spider genus *Stygopholcus* we show that some specimens originally published as a separate species (*S. montenegrinus*) and later assigned as a subspecies of *S. skotophilus* are genetically well separated (gdi=0.8) from *skotophilus* (fig. S11). In the wasp genus *Pteromalus*, the results suggest that the species *P. albipennis* and *P. eudecipiens* should probably be synonymized, while the individuals currently classified as *P. brachygaster* sp1 and sp2 belong to two distinct lineages, which await formal taxonomic treatment. Among the amphibians, *Discoglossus pictus* and *D. scovazzi* are cryptic species that cannot be distinguished based on morphology or bioacoustics, but that share a narrow contact zone without relevant inter-species gene flow, while *Rana pyrenaica* and *R. temporaria* are closely related but highly distinct in morphology and ecology (*48,57*). These well-established but closely related species were recovered in all USCO-based species delimitation analyses, suggesting that under-splitting will rarely be an issue, except for extremely young adaptive radiations. On the contrary, there is a wide hybrid zone of about 200 km between *Discoglossus galganoi jeanneae* and *D. g. galganoi*, which are thus to be seen as intraspecific lineages and therefore can serve as benchmark to detect over-splitting. Hence, the fact that most multispecies coalescent analyses based on USCOs or *COI* separate *galganoi* and *jeanneae* as different species suggests over-splitting is commonplace. It also highlights that some of the SNP-based approaches not relying on coalescence models (NMDS, *STRUCTURE*) are less prone to objective over-splitting. This was also found for the *tr2* analysis based on assembly strategies A5 and A6, but in those cases it may be caused by a lower quantity of available data.

Results of the phylogenetic analysis including all taxa with 260,233 amino acids or 780,699 nucleotides in 978 USCOs indicate that USCOs can also differentiate between higher taxa at the level of orders or classes. Thus, USCOs are suitable to assign unknown samples (with lacking species-specific reference data) to higher systematic categories, such as genera, tribes, families, orders etc. (*21*) for which single markers often fail. This was one of the main reasons to suggest USCOs as universal standard marker system among metazoans (*21*). For the tree analysis including all taxa, we found that the phylogenetic tree obtained from the amino acid data set is compatible with the current view of within-arthropod relationships (*23*) (fig. S9). Only the placement of the tetrapod species with respect to the arthropods is in conflict with the generally accepted phylogeny. This could be attributed to a lack of taxa between the distantly related groups. The nucleotide tree (Fig. 2A, fig. S9) based on all three codon positions shows conflicts with the accepted insect phylogeny (*23*), but since this data also includes the hypervariable third codon positions, this does not come as a surprise. The nucleotide tree based on all codon positions is shown, since it contains more phylogenetic signal for very recent divergences. Altogether these results show that USCOs are very well suited to elucidate relationships within and among the taxonomic groups studied here.

Despite the observed over-splitting with multispecies coalescent species delimitation approaches, which is likely to be resolved in future species delimitation methods, our results show that USCOs provide a powerful marker system to distinguish closely-related species. Similar outcomes of over-splitting have also been reported in previous studies using the multispecies coalescent, even if only a few markers were used (*58–62*). The main difficulty to date, particularly in young species, is thus to distinguish population structure from speciation (*58,59,62*), a problem that seems to be more serious rather than alleviated in datasets with large numbers of loci, because population structure is more clearly resolved with more data (*63,64*). This has been demonstrated with *tr2* (*60*) and is also expected for *BPP* (*51*). In our analyses with all USCO data, even syntopic specimens, which in the phylogenetic trees are shown to be closely related, are often split into different species.

The genealogical divergence of populations/species as expressed by the genealogical divergence index (gdi) (*52*) turned out to be of limited use as general proxy to evaluate species status: (i) many of the infraspecific splits did not allow a calculation of the gdi as it cannot be calculated for lineages represented by only one sample and (ii) many gdi values were found to be in the unspecific value range, where species status could not clearly be determined. Nevertheless, the calculation of the gdi enabled us to evaluate at least some of the observed over-splitting.

Finally, USCOs conform to all of our four initial criteria (a-d) as a suitable and superior marker system for species delimitation (see Introduction). The striking success of these protein-coding and rather conserved markers to resolve shallow lineages alleviates the need to exploit introns for species delimitation purposes. This is advantageous, because the nucleotides of orthologous introns are more difficult to align, especially if the simultaneously examined taxa are phylogenetically relatively distant from each other. To our knowledge, USCOs represent the only universal multi-marker system that is applicable to DNA taxonomy of all metazoans (*21*). Our results show USCOs to perform successfully for both arthropods and vertebrates and, at the same time, to overcome all problems of *COI* barcoding.

Some of our study cases found an extraordinary congruence of USCO-based analyses with species hypotheses derived from previous integrative approaches with independent evidence (Supplementary Information). However, given the tendency of over-splitting, our results also underline the steady need for critical evaluation and additional refinement (when possible) of the results of species delimitation (*16*) with additional evidence and methodology (*57,65,66*), also when inferring species boundaries with data-rich multi-gene datasets under the multispecies coalescent model (*18,19,60*). Limitations of current species delimitation approaches (*58*), but also the nature of species and speciation (*67*), continue to urge for an integrative evaluation of results of species delimitation (*14*) and emphasize the need to further improve protocols, algorithms and implemented models for species delimitation based on DNA data. The accurate application of these will require knowledge of the speciation circumstances (e.g., geography, hybrid zones, host information) and sufficient sampling depth, taxonomically and geographically. We show that USCOs are very well suited to significantly improve cross-taxon hypothesis testing and to substantially increase the accuracy, sustainability and comparability of DNA markers in species delimitation: they are primary keys to advancing taxonomy in the era of genomics.

## Materials and Methods

### Organism groups and samples of the case studies

We sampled the DNA of USCOs of selected species of seven genera representing six major groups of Arthropoda (Coleoptera: *Pleophylla*; Diptera: *Sphaerophoria*; Hymenoptera: *Chrysis* and *Pteromalus*; Lepidoptera: *Taygetis*; Araneae: *Stygopholcus*; Myriapoda: *Lithobius*) and of selected species of two genera of Amphibia (Anura: *Discoglossus, Rana*) (table S1; Supplementary Material). Sets of taxa were chosen to include (i) well-studied examples of closely related species, as well as genetically divergent but clearly conspecific lineages, to provide controls for the accuracy of species delimitation procedures; and (ii) particularly challenging cases of morphologically cryptic lineages or species complexes affected by mitochondrial introgression, where species delimitation with classical data sets is likely to fail.

### Data generation and assembly

USCO data were produced using target DNA enrichment (*5, 21*). Bait design (*5*) and wet lab procedures are extensively described in the Supplementary Materials.

We particularly considered different data assembly approaches which may impact species delimitation if very closely related taxa are compared. A major objective was to maximize the number of USCOs found per studied taxon, but also to infer the robustness of species delimitation given different theoretical backgrounds of the data assembly methods. We used seven different assembly techniques (A1-A7) partly using new and partly published pipelines and verified orthology (for details see Supplementary Materials, fig. S1).

In addition, we also generated reduced datasets for all approaches in which alignment positions having a gap in at least one individual were removed. For A1 and A2, we generated datasets in which all positions were removed that were recovered as either having a gap or ambiguity (i.e. heterozygous) in at least one individual.

### Phylogenetic analyses

For phylogenetic analysis, we followed a two-fold strategy:

1. For all resulting assemblies (A1-A7), the alignments of all recovered genes (see table S2) were concatenated into a single dataset for each of the nine study cases using in-house scripts (see Dryad data). Maximum-likelihood phylogenetic analyses based on these datasets using all nucleotides were conducted with *IQ-TREE* v. 1.6.3 (*56*). We conducted tree searches, (i) using a non-partitioned analysis with the model GTR+I+G (“explorative” run; for all data combinations), and (ii) a more thorough approach where the datasets were initially partitioned into individual genes and the best model and partitioning scheme was determined using the option –m MFP+MERGE in *IQ-TREE*. This analysis was repeated 50 times for all taxa and full datasets (just A2) selecting the tree and model with the highest likelihood. Branch support was assessed by ultrafast bootstrapping (*68*) within *IQ-TREE* using 1,000 replicates. We also combined the results of A3 for all taxa into a single super-alignment, including also the reference sequences (table S3) that were used for bait design, and excluding all regions not covered by the Hidden Markov Models (HMMs)(*69*). Approach 3 is the method best suited for this purpose, as the sequences were assembled and aligned directly making use of the HMMs as references, which were the same for all taxa. A phylogenetic analysis of the concatenated alignment was performed as described above for nucleotides and amino acid sequences. For both datasets, the model of evolution was determined using *ModelFinder* (*70*) as implemented in *IQ-TREE*.
2. For all assemblies (A1-A7), phylogenetic analyses of individual gene alignments were carried out using *IQ-TREE* and all nucleotides with the model GTR+I+G, and the gene trees were then used as input for coalescent-based analyses with *ASTRAL III* v. 5.6.1 (*55*). As an additional measure of support, we counted the number of gene trees for which each clade of the concatenation-based trees was recovered using *nw_clade* (part of the *Newick Utilities* 1.6 package) (*71*). *ASTRAL* analyses were performed for all USCO datasets, both full and reduced.

### Analysis of single nucleotide polymorphisms (SNPs)

SNPs were extracted from the USCO datasets obtained with A2 with the software *SNP-sites* (*72*), excluding low-quality sequences marked by lowercase nucleotides. Additionally, they were filtered in the following steps, using custom Perl scripts: (i) all non-informative SNPs (with all individuals except one having the same allele) were removed from the dataset. (ii) SNPs for which no sequences were available for at least 50% of individuals were removed. (iii) for each gene, only the SNPs with information for the highest number of individuals were taken into account. In the case of *Pleophylla*, we used two datasets, one comprising all species, the other one being limited to the *P. pilosa* species complex for which species cannot be distinguished with *COI* (Fig. 2). The SNPs were then used for a population structure analysis by clustering with the software *Structure* v. 2.3.4 (*42*) using an MCMC chain length of 50,000 (burnin of 20,000) and a number of ancestral populations (K) from 1 to 10. For each K value, the analysis was repeated ten times. This procedure places individuals into a predetermined number of clusters, which can be varied across independent runs of the algorithm. Additionally, non-metric Multidimensional Scaling (NMDS) analyses were conducted with the software *PAST* v. 4.03 (*73*). We repeated the analyses at least ten times to avoid local optima, preferring the results with the lowest stress value. For NMDS, only SNPs with exactly two alleles were used. With a custom Perl script, SNPs with more than two alleles were excluded and the more common allele was coded as 0 and the rare allele as 1, if heterozygous, or 2, if homozygous.

### Species delimitation analysis

USCO gene data of species, representing independently evolving meta-populations (*45,74*), should fit a species tree with gene tree distributions described using the multi-species coalescent model (*75*). Consequently, we applied various implementations of the multispecies coalescent model to each study case employing parametric (*18,19,76,77*) and nonparametric (*50*) methods to delimit species based on genomic data (see Supplementary Materials for further details). Species delimitation analyses were run on full and reduced datasets from all seven assembly approaches. To avoid potential biases from a predefined clustering, we assigned each specimen to a separate cluster.

### COI-benchmarking

USCO-based tree inferences and results of species delimitation were compared with those from *COI* sequence data. Details on *COI*-related wet lab procedures, assembly techniques, tree searches, and species delimitations are provided in the Supplementary Materials.

## Acknowledgments

Amphibian samples and data were contributed by Pierre-Andre Crochet, David Donaire, and Pedro Galán. We also thank Tomas Flouri, Michael Hofreiter, Loïs Rancilhac, and Ziheng Yang for fruitful discussions, as well as Karen Meusemann, Alexander Donath, and Lars Podsiadlowski for their help to access 1Kite and ZFMK graduate school data.

## Author contributions

Design of the study and acquired funding: D.A., O.N. and B.M.

Conceptualized and supervised data collection and sequencing: D.A., C.H., M.V., C.M., O.N., B.M. and J.E.

Assembly and analysis of USCO data: L.D.

Provided and identified specimens: D.A., M.E., H.B., B.A.H., C.H., X.M., O.N., R.S.P., T.W., K.W.

*COI* data assembly: C.M., J.E., C.B.

Analysis of the *COI* data: C.B.

Writing—original draft: L.D., J.E., C.M., O.N., D.A.

Writing—review & editing: L.D., J.E., C.M., S.K., C.B., H.B., M.E., B.A.H., C.H., X.M., R.S.P., M.V., T.M., K.W., O.N., and D.A.

## Data and materials availability

Specimen data including reference to collection permits are given in table S1. Raw sequence data are deposited on NCBI (see table S1). Protocols, code, trees, and alignments are available on Dryad (available for this paper at https://doi.org/10.5061/dryad.hhmgqnkg5).

