## Supplementary Material for "Standardized nuclear markers advance metazoan taxonomy"

### Material and methods

*Organism groups and samples*

Specific information on the species identities and sampling locations is given in table S1. All analyzed samples are deposited either in the BioBank of the AG Niehuis at the University Freiburg (most samples of the genus *Chrysis*), in the BioBank of the ZFMK (all remaining samples of arthropods except *Taygetis* which are currently housed at Florida Museum of Natural History, Gainesville, Florida), or in the BioBank of the Vences lab at Braunschweig University of Technology (all amphibian samples).

*Research and collection permits*

The information on sampling permits for the respective study case is given below, including along each specimen in table S1.

**Coleoptera (*Pleophylla*). South Africa:** Eastern Cape (Permit No. WRO 122/07WR and WRO123/07WR), Gauteng (Permit No. CPF6 1281), Limpopo (Permit No. CPM-006-00001), Mpumalangma (Permit No. MPN-2009-11-20-1232), Cape Province (Permit No. AAA0007-00097-0056), and Kwazulu-Natal (Permit Nos. OP3752/ 2009, 1272/2007, 3620/2006).

**Lepidoptera (*Taygetis*). Ecuador:** Lepidoptera samples were obtained with the support of the Instituto Nacional de Biodiversidad and Ministerio del Ambiente, most recently under permit number No. 006-19 IC-FLO-FAUDNB/MA.

**Diptera (*Sphaerophoria*). Cyprus:** Ministry of Agriculture, Department of Environment (Permit Nos. 02.15.001.003 and 04.05.002.005.006) and the Department of Forests (Permit No.

2.15.05.3). **Georgia:** Agency of Protected Areas (Permit No. 1452-0-2-201905011744) and the Ministry of Environment Protection and Agriculture of Georgia (Permit No. 4654-01-2-201905081440). **Germany:** 55-8841.06/0, LAU-MLU_43.14-Schn., NP-Eifel_WV-Nr. 061/2013, SGD-Nord_425-104.1204.

**Hymenoptera (*Chrysis*). Germany:** Hessen-Forst LBL (Permit Aktenzeichen IV.2 R 28), Nationalpark Schwarzwald (Baden-Württemberg; Permit, Schreiben vom 6. April 2020), Struktur- und Genehmigungsdirektion Süd (Rheinland-Pfalz; Permit Aktenzeichen 42/553-254).

**Hymenoptera (*Pteromalus*).** N/A

**Arachnida (*Stygopholcus*). Crete:** “SPIDOnetGR”, ARISTEIA II Programme, NSRF 2007-2013, co-funded by the European Union (European Social Fund) and National Resources under the Operational Programme “Education and Lifelong Learning”- Action 81324 (M. Chatzaki). **Croatia:** “Inventory and mapping of the subterranean, spring, and bat fauna, with making of the biospeleological cadaster of Nature Park Biokovo”, Ur. Br. 584/2001 – financed by Nature Park Biokovo (M. Pavlek). **Montenegro, Bosnia and Herzegovina:** “A catalogue of spiders (Arachnida, Araneae) of Montenegro”, supported by the Montenegrin Academy of Sciences and Arts (G.S. Karaman).

**Myriapoda (*Lithobius*). Germany:** Permits 5-N-A1007/480-2012; City Bonn 2014 and 2016; NRW 2015; Rhein-Sieg 2013; RU5-BE-66/008-2012.

**Anura (*Rana/ Discoglossus*). Germany:** 16-605105-269/14 (Landkreis Helmstedt); **Spain:** 201799900546597-06/10/2017, Junta de Andalucia; Xunta de Galicia, Ref. EB-041/2017 (years of validity 2017, 2018, 2019). **Morocco:** May 2013 HCEFLCD/DLCDPN/DPRN/CFF, High Commissariat for Water and Forest, Morocco. Other samples provided by collaborators.

*Bait design*

Baits to capture gDNA of the 978 metazoan USCOs in arthropods were designed with the *BaitFisher* software package (*BaitFisher* v. 1.2.8 and *BaitFilter* v. 1.0.6) (*5*), using a tiling design with three consecutive 120 bp long baits, spanning a 160 bp bait region with a new bait every 20 bp. Generated bait DNA sequences were based on transcriptome and genome sequences of species that were phylogenetically related as closely as possible to the target species (table S3). The reference sequences were extracted from transcriptome assemblies or coding sequences of genomes using *Orthograph* v. 0.6.3 (*44*). *Orthograph* was provided with the original Hidden

Markov Models from the *BUSCO* software package v. 3.0.2 (*38*). Amino acid sequences of the 65 species that were used to define USCOs were used as reference sequences when performing the reverse BLAST searches. Baits were designed so that they did not cross exon-intron boundaries by using the alignment cutting option in *BaitFisher*. With this option, coding (CDS) regions were excised with the aid of an annotated reference genome of a related species (fasta file and gff file, see table S3 for the references used for each case study) before baits were designed. DNA sequences from reference genomes that solely served for the identification of exon-intron boundaries were not used for inferring bait DNA sequences (see option "remove-reference-sequence" in the *BaitFisher* configuration file), except in the case of *Nasonia* whose gene sequences (*78*) were considered when designing baits for enriching USCOs of species in the genus *Pteromalus*. In a second run, baits were designed without alignment-cutting for those USCOs for which bait design did not succeed in the first run due to continuous CDS that were shorter than the length of the bait region. In both runs a clustering threshold of 0.15 was used. Bait design was separately conducted for each of the targeted taxon-groups and the resulting baits were subsequently combined; i.e., only one bait solution was ordered containing baits for all taxon-groups. The combined bait files were checked for duplicate baits before ordering at Agilent Technologies Germany.

For the amphibian samples, the enrichment approach used had the goal to create a set of universal amphibian markers that could be targeted using a single set of probes. To select and design markers, we used the FrogCap marker set (GitHub: https://github.com/chutter/FrogCap-Sequence-Capture) (*79*). We selected all FrogCap markers that were successfully captured across Anura, selecting markers captured in the ranoid, hyloid, and archaeobatrachian clades. Next, these markers were matched against the *Ambystoma* genome (*80*), and markers were retained if they matched with at least 65% similarity. The total number of markers was 7,720 (UCEs: 2,122; exons: 5,598). To these, we added a set of USCO markers present across amphibians, based on the *Ambystoma mexicanum*, *Nanorana parkeri, Oophaga pumilio*, *Rana catesbiana*, *Rhinella marina*, and *Xenopus tropicalis* genomes (*80-84*). We used the program *BUSCO* (v. 3.0.2) (*85*) to locate and identify the Metazoa and Tetrapoda USCO markers in these genomes. We included all metazoan genes found in all genomes, which totaled 677 out of 978 genes. We also added 29 genes from the Tetrapoda set to use the full size of the probe set. We then extracted each gene from each genome and separated all genes into exons based on the boundaries determined by each genome's annotations. We aligned each exon and created a consensus sequence for that exon from across all the genomes. The consensus sequences were used to design a MYbaits-2 (40,040 baits) custom bait library (Arbor Biosciences), using 120mer baits to best capture sequences with greater than 5% divergence from the probes. The finalized set of markers were separated into probe sequences following a 2x tiling scheme, starting 20 bp behind the start codon of the exon and tiling 120 bp probes every 60 bp until 20 bp past stop codons. Individual probes were filtered using these criteria: 1) probes that matched 70% length or greater to multiple locations in the genomes above with *BLAST* with a 70% similarity were excluded; 2) probes with a GC content between 30–50% were kept; 3) probes without any repetitive sequences based on *RepeatMasker* (*81*) annotations using the online server were kept; and 4) probes which had no matches with *BLAST* to other probes (using a 70% similarity criterion) were kept. After filtration 40,211 baits remained; whole genes from the Tetrapoda set and their baits were randomly selected to total 189 baits from 7 genes to drop from the dataset to fit into the 40,040 bait limit. This resulted in a final set of 8,720 markers covering a total of 2,702,951 bp. The USCO markers totaled 14,153 baits (Metazoa: 13,196; Tetrapoda: 957) while the FrogCap markers totaled 25,884 baits (UCEs: 3,844; exons: 22,040).

*Wet lab procedures*

*Arthropods*: Genomic DNA was extracted using Qiagen Blood & Tissue Kits (Qiagen, Hilden, Germany) following the manufacturers' protocol except that a RNAse-digestion step was added after lysis and isolated DNA was resolved in water. 100 ng genomic DNA (if available) was sheared with a Bioruptor PICO sonicator (Diagenode s.a., Seraing, Belgium) to gain approximately 350 bp long DNA fragments. End-repair and A-tailing were done with Agilent SureSelectXT2 Reagent (Agilent Technologies Inc., Santa Clara, USA) Kit, while NEBNext Quick Ligation Module, NEBNext Multiplex Oligos for Illumina (Dual Index Set1) and NEBNext Q5 HotStart HiFi PCR Master Mix were used for pre-capture adaptor ligation and dual-indexing amplification. During PCR the adaptor-ligated DNA was dual-indexed and amplified with the following PCR-program: ten, twelve or sixteen cycles of (98°C/10s–65°C/75s) – the number of cycles was dependent on the concentrations after the A-tailing concentration measurement. Between these steps, the samples were purified with AMPure XP beads (Beckman Coulter GmbH, Krefeld, Germany) in different ratios and the DNA quantity was checked with Quantus Fluorometer (Promega, Fitchburg, Wisconsin, USA). After library PCR the quality was checked additionally with a Fragment Analyzer (Advanced Analytical, now Agilent Technologies Inc.).

After amplification of the libraries, equal DNA amounts of eight samples a time were pooled. SureSelect XT2 Pre-Capture ILM Module Box 2 Kit (Agilent Technologies Inc., Santa Clara, USA) was used for the following hybridization steps. After dsDNA was denatured for 5 min at 95°C in a thermal cycler, hybridization took place at 65°C for approximately 48 h. The library of hybridized baits and DNA fragments was captured with Dynabeads MyOne Streptavidin T1 beads (Thermo Fisher, Waltham, Massachusetts, USA). The captured libraries were amplified in twelve or fourteen cycles, dependent on the final concentration needed for sequencing, using the SureSelect XT2 primer mix and Herculase II PCR Master Mix and purified with AMPure XP beads. Libraries were sequenced on an Illumina Nextseq 500 platform (StarSEQ GmbH, Mainz, Germany) with 260 million reads in total for twelve pools, so that we had 21.7 million reads per pool of eight samples.

*Amphibian samples*: Genomic DNA was extracted from the tissue samples using a Promega$^{TM}$ (Madison, USA) Maxwell bead extraction robot. The obtained DNA was quantified using a Promega Quantus$^{TM}$ fluorometer. Approximately 500 ng total DNA was acquired per sample and set to a volume of 50 µl through dilution (with $H_2O$) or concentration (using a vacuum centrifuge) of the extraction when necessary. The genomic libraries for the samples were prepared by Arbor BioSciences (Ann Arbor, USA) library preparation service. Prior to library preparation, the genomic DNA samples were quantified using a Qubit and up to 4 µg was then taken to sonication with a QSonica (Newtown, USA) Q800R instrument. After sonication and SPRI bead-based size-selection to modal lengths of roughly 300 bp, up to 500 ng of each sheared DNA sample were taken to Illumina TruSeq-style sticky-end library preparation. Following adapter ligation and fill-in, each library was amplified for 6 cycles using unique combinations of i7 and i5 indexing primers, and then quantified with a Qubit. For each capture reaction, 125 ng of 8 libraries were pooled, and subsequently enriched for targets using the *MYbaits* v 3.1 protocol (*MYbaits* User Manual version 3.1). Enrichment incubation times ranged from 18–21 hours. Following enrichment, library pools were amplified for 10 cycles using universal primers and subsequently pooled in equimolar amounts for sequencing. Samples were sequenced on an Illumina HiSeq X with 150-bp paired-end reads in a shared lane with 96 total samples.

*Data assembly*

Before assembly, adapters and low-quality regions were trimmed with *fastq-mcf* (*86*). We expect different assembly approaches to have different yields of USCO DNA sequence data. Since obtaining taxonomically informative DNA sequence information is crucial for reliable phylogenetic and species delimitation results, it is necessary to assess which assembly approach is most suitable for our purposes. Therefore, the filtered paired-end reads were assembled using seven different approaches (see table S1; fig. S1) in order to examine their eventual impact on phylogenetic analyses and species delimitation results:

*Approach 1 (A1):* Reads were mapped with the BWA-MEM algorithm of *bwa* v. 0.7.17 (available from https://bio-bwa.sourceforge.net) (*40*) against reference DNA sequences of the targeted exons from taxa closely related to the studied ones (see table S3). For *bwa*, default parameters were used, except that the length of the seed sequence was increased to 30 bp to reduce the likelihood of generating false positives. For every individual, a diploid consensus sequence of the mapped reads was generated with *samtools* 1.6 and *bcftools* 1.6 (available from https://github.com/samtools/bcftools) (*87*). This resulted in sequences that were already aligned, a further alignment was therefore not necessary.

*Approach 2 (A2):* Reads were assembled with *Trinity* v. 2.8.3 (*88*), and *Orthograph* v. 0.6.3 (*44*) was used with default settings to search the resulting contigs in each individual for USCO orthologs using the pre-existing Metazoa dataset. For every gene, the DNA sequences output by *Orthograph* for all individuals of a genus were extracted and the longest of these sequences was chosen. This sequence was further used as reference for mapping the reads of all individuals with *bwa*. All additional steps are identical to those used in A1.

*Approach 3 (A3):* The first steps (*Trinity, Orthograph*) were identical to those used in A2. However, instead of using them as a mapping reference, the protein sequences output by *Orthograph* were aligned against the Hidden Markov Models (HMM) generated by *Orthograph* for the USCO Metazoa dataset with *hmmalign* (part of the *HMMER* package available from www.hmmer.org). Positions outside the length of the HMM were removed with a custom Perl script. Nucleotide alignments based on these protein alignments were created with *pal2nal* (*89*).

We also assembled the data with the published pipelines *IBA* (A4) (*90*), *Hybpiper* 1.2 (A5) (*42*), and *Phyluce* 1.6 (*43*) using the default parameters of the respective software. However, for *Phyluce* we used DNA sequences that were pre-assembled with *Trinity* (from A2) instead of assembling them within the pipeline. For *Phyluce*, we conducted two assemblies, one (A6) using the genomic/transcriptomic reference data, the other (A7) using the *Orthograph* output as a reference as in A2. The output of *Hybpiper* was aligned with *hmmalign* as described for A3. For *IBA*, which does not differentiate between coding and non-coding sequences, the output was aligned with *MAFFT* 7.305b (*91*) using the same reference alignments that were used for bait design. The *IBA* approach failed to recover a sufficient amount of sequences in both *Rana* and *Discoglossus* where for some specimens no sequences were found at all. Results from the *IBA* approach were therefore not further used for those study cases.

For those approaches for which amino acid sequences were obtained (A1, A2, A3 and A5), all gene alignments were quality filtered with the program *OIilnSeq* 0.9.3 (available upon request from C.M.). This program uses a sliding window approach to detect outlier sequences on the amino acid level. In each window, the mean BLOSUM62 similarity score of each sequence to all other sequences is calculated. For the set of mean scores, the interquartile distance (IQD) is calculated and sequences for which the mean score is lower than the lower quartile by an amount of at least IQD * f, where f is a specified factor, are considered as outliers. Here, we used the default parameters, with a sliding window of length 30 and f=1.5. DNA sequence segments identified as outliers were masked with X in the protein alignments and with N in the corresponding nucleotide alignments.

After filtering, alignment positions or whole exons recovered in less than three individuals were removed from the final dataset.

All specimen samples were benchmarked additionally with *COI* barcodes (see below).

*Orthology verification*

As paralogous sequences can mislead phylogenetic and other analyses (*92,93*), it is important that multiple sequence alignments contain only orthologous sequences. However, this is non-trivial if target genes are extracted from enriched whole genomic libraries. The different assembly approaches use different techniques to ensure that only orthologous sequences are combined in multiple sequence alignments.

In A1 and A2, sequences are mapped iteratively against a start sequence. Here the long minimum mapping seed of 30 bp ensures that reads are only mapped to the starting sequence if the identity is very high. An orthology assessment in the sense of a reciprocal similarity searches is not done.
In A3 a thorough orthology assignment is accomplished with *Orthograph* (*44*), which uses reciprocal searches of putative sequences of genes against the whole official gene sets of the reference species. From a theoretical point of view this is probably the best methodology for identifying exclusively orthologous sequences. Unfortunately, the assembly of the enriched sequences remains a problematic step in A3, as theoretically different variants could be assembled to a single sequence.
In A4, reads similar to the query sequence are assembled with *Bridger* (*94*). In a few cases, this produced more than one contig, indicating the presence of paralogous or non-target sequences. Such genes were then manually checked and removed from the dataset.
*Hybpiper* (A5) assembles reads which are found to be similar to a given reference sequence with Spades, and uses the longest of the resulting contigs. In cases where two or more contigs have a length of at least 75% of the reference, a paralogy warning is given.
*Phyluce* (A6 and A7) uses pre-assembled contigs to search for sequences similar to a given specified reference sequence. In cases where two or more contigs are found for a given reference, that locus is automatically excluded from the dataset.

*Species delimitation analysis*

We applied various implementations of the multi-species coalescent model to each study case using parametric (*18,19,76,77*) and non-parametric (*50*) methods to delimit species based on genomic data.
For the first, we analyzed both full and reduced USCO data with the program *BPP* v. 4.1.4 which tests the validity of species-level groupings using multi-gene data with a Bayesian approach (*49*). To avoid potential biases from a predefined clustering, we assigned each specimen to a separate cluster. We performed *BPP* analyses of type A10 (*49*), in which the species tree is fixed but species delimitations are inferred by the analysis (*49*). As guide trees, we used the species tree generated with *ASTRAL* for the full dataset of the respective approach. The Monte Carlo Markov Chain (MCMC) was run for 200,000 generations, with a sample frequency of 2 and burn-in of 8,000.

In our *BPP* analyses, all sequences were coded as haploid. We also tried to run analyses with diploid data in the diploid mode of *BPP* (for data from A1 and A2), but we came to the conclusion that for the larger data sets with a high degree of heterozygosity, such as e.g. the *Pleophylla* data set, these analyses required too much computational resources und would take months to complete. This was likely caused by the fact that in the diploid mode, all possible combinations of heterozygous sites within a locus are taken into consideration, which drastically increases the number of site patterns. Diploid analyses of some small data sets did complete, but had the tendency to support unrealistic models in which all or almost all individuals were classified as separate species (results not shown). The cause of this effect is unknown and should be explored with more empirical data. Since *BPP* with heterozygous data coded as haploid and being analyzed under the haploid option in *BPP* theoretically would incorrectly handle the data (Z. Yang, pers. comm.; Feb 12th, 2021), but handling the heterozygous data as diploid was not an option either, we feel at the moment that the reduced data analysis would generate be the most reliable results.
To assess the robustness of delimitations we applied a series of analyses with various theta (population size) and tau (divergence time) prior combinations, which has been shown to be crucial for obtaining unbiased results (*53,61,95*). Specifically, we used all combinations of theta priors with beta = 0.4, 0.04, and 0.004, as well as tau priors with beta = 0.2, 0.02, and 0.002 (with alpha = 3 in all cases). Analyses were repeated five times, and the median values of the posterior probabilities of the five runs were used for final species inference. In order to distinguish the influence of priors and data, we also performed an additional run without data for all parameter setups, thus sampling the marginal prior density. Finally, we also reran all *BPP* analyses with reduced data and the *BPP* guide trees generated with *ASTRAL* from the reduced data set (removing only gaps, not ambiguities).
We also calculated the Genealogical Diversity Index (gdi) (*52*) for the datasets obtained with A2 and intermediate prior values (beta = 0.04 for theta, beta = 0.02 for tau) with *BPP* using the approach of Leaché (*51*). Specifically, we conducted a *BPP* analysis where the species delimitation and species tree are fixed, in order to infer theta and tau values for each branch or node, respectively, in the tree (*BPP* analysis type A00) (*49*). The option “e” was used for the theta priors, specifying that the theta values are explicitly estimated during in the Bayesian analysis, rather than being integrated out. As this may lead to a slower convergence of the

MCMC, the analysis was conducted for 1,000,000 generations with a burn-in of 100,000. The other parameters were as described above. We used the most likely species delimitation inferred for this prior combination by the full *BPP* analysis and the tree inferred by *ASTRAL*. For each grouping containing more than one specimen, gdi was then calculated as

1-e^(-2*tau_AB/theta_A),

where tau_AB is the inferred divergence time of the clade from its sister group, and theta_A is the inferred population size for the clade. We determined the final theta and tau values as the medians of the (median theta and tau) values from five independent runs. The gdi then represents the probability that sequences of the same gene from two different individuals within the clade share a common ancestor after its divergence from the sister clade. According to Jackson (*52*), a gdi < 0.2 supports the conspecificity of the two sister clades, while gdi > 0.7 supports them being distinct species, and intermediate values are ambiguous.

To assess whether differences in the results between the full and reduced datasets are simply due to the lesser amount of data, we performed jackknife analyses on the A2 dataset of *Pleophylla*. Using a custom Perl script, we randomly removed gene loci from the full dataset to match the number of loci of the dataset without gaps. Additionally, we randomly removed nucleotide positions until the total number of positions also matched that of the reduced dataset. We constructed five such jackknife datasets and analyzed each of them for three of the nine prior combinations used for our main analyses (low theta/high tau, medium theta/medium tau, low theta/high tau). For each of the five jackknife datasets, five replicate analyses were conducted, using the same conditions as for the main analysis.

To infer the effect of the theta and tau priors on the posteriors, we also conducted a *BPP* analysis of type A00 (*49*) for all nine previously described prior combinations and the *Pleophylla* A2 dataset. To increase the likelihood of convergence, we used a burnin of 100,000 and an MCMC length of 1,000,000, sampling every 10$^{th}$ generation. The analysis was repeated five times, and the median posterior probability from these was used as proxy for the prior effect on split behavior.

We also used the program *tr2* (*50*) to infer species boundaries based on variations in gene trees. As input, we used the gene trees obtained with *IQ-TREE* that include all specimens within the respective taxon (as trees with missing specimens cannot be used by the program). Input trees were rerooted using *nw_reroot* (part of the Newick Utilities 1.6 package) (*71*). For this we chose the root found in the phylogenetic analyses of the whole datasets including outgroup taxa. *tr2* was used with all USCO data as well as with reduced data. Finally, we tested the newly developed quartet-based species delimitation method *SODA* (*96*), which is designed to be scalable to large datasets, on the A2 datasets for all nine genera.

Congruence of tree topology and species delimitations was assessed with respect to morphology-based *a priori* species identifications and *COI*-based species delimitations (*COI* benchmarking) (*16*).

*COI benchmarking*

USCO-based species delimitation results were compared with results obtained from species delimitation methods using *COI* sequence data. For the latter we applied the Poisson tree process (PTP) (*97*), statistical parsimony analysis (*98,99*), and Automatic Barcode Gap Discovery (ABGD) (*100*).

For *COI* benchmarking, the *COI* gene (5′-end) of all arthropod specimens was Sanger-sequenced, except for *Pleophylla*, where we used the *COI* gene (3′-end) dataset (*53*). Lab work followed the standard protocols of the German Barcode of Life project (*101*).

DNA was extracted from various body parts using the Qiagen DNeasy Blood and Tissue Kit (Hilden, Germany). The PCR reaction was carried out in total reaction mixes of 20 µl, including 2 µl of undiluted DNA template, 0.8 µl of each primer (10 pmol/µl), and standard amounts of the reagents provided with the "Multiplex PCR" kit from Qiagen (Hilden, Germany) using various sets of primers (table S7). Thermal cycling was performed on Applied Biosystems 2720 thermal cyclers (Life Technologies, Carlsbad, CA, USA), using a PCR protocol with two cycle sets, combining a "touchdown" and a "step-up" routine as follows: hot start Taq activation: 15 min at 95°C; first cycle set (15 repeats): 35 s denaturation at 94°C, 90 s annealing at 55°C (−1°C per cycle) and 90 s elongation at 72°C; second cycle set (25 repeats): 35 s denaturation at 94°C, 90 s annealing at 40°C, and 90 s elongation at 72°C; final elongation 10 min at 72°C. Unpurified PCR products were subsequently sent for bidirectional Sanger sequencing to BGI Tech Solutions (Hongkong, China).

Raw DNA sequences were assembled (forward and reverse sequence) and edited in *Geneious* R7 (version 7.1.3, Biomatters Ltd.) to correct base-calling errors and to assign ambiguities (when forward and reverse sequence were not congruent for certain nucleotides). Sequences were aligned with *Muscle* (*102*) as implemented into *Geneious* using the default settings. Primer regions were trimmed subsequently. For the frog specimens (*Discoglossus*, *Rana*), we searched the enriched libraries for reads aligning well to complete *COI* reference sequences as follows: we created a reference amino acid consensus sequence separately for Ranidae and *Discoglossus* from *COI* data available in NCBI Genbank (https://www.ncbi.nlm.nih.gov/genbank/). The list of NCBI accession numbers of the sequences from which the two consensus sequences were determined as well as the consensus sequences are given in table S6. The amino acid sequences from NCBI were aligned separately for the two taxonomic groups with *mafft-linsi* v. 7.475 (*91*) and consensus sequences were determined from these alignments with in-house scripts and a consensus threshold of 50%. Finally we aligned the reads from each species library and the corresponding reference amino acid sequence with *exonerate* v. 2.4.0 (*103*), using the options "–frameshift -9 –Q protein –T dna –model protein2dna –showvulgar 1" and specifying the vertebrate mitochondrial genetic code. In this way we obtained alignments of reads for each sequence library, i.e. for each specimen. From the resulting read alignments, we computed a 50% majority consensus sequence. If no nucleotide at a given site had a majority of 50%, the unknown nucleotide N was inserted. The NCBI accession numbers of all specimens are given in table S1.

Phylogenetic trees of aligned *COI* sequences were estimated using the maximum likelihood (ML) criterion as implemented in *PhyML* v. 3.0 (*104*) with automatic model selection (*105*) on the web interface (http://www.atgc-montpellier.fr/phyml/; accessed August, 18th, 2020). Branch support was measured using the approximate likelihood ratio test (aLRT) as implemented in *PhyML* (*106*). The resulting tree was midpoint-rooted in *FigTree* v1.4.3 (*107*), serving as basis for the tree-based species delimitation analysis.
We used the two variants of the Poisson tree process model (*PTP*) on the *PTP* web server (https://species.h-its.org/; accessed August, 20th, 2020): *bPTP*, which adds Bayesian support (pp) values to branches that delimited species in the input tree, and the refined multi-rate *mPTP*. *PTP* uses the shift in the number of substitutions at internal nodes to identify branching rate transition points (*97*) which indicate speciation events. We used default settings for the *bPTP* analysis (100,000 MCMC generations, a thinning of 100, a burn-in of 0.1, and a random seed of 123).

Statistical network analysis as performed with *TCS* v. 1.21 (*108*) separates the sequence data into clusters of closely related haplotypes connected by changes that are non-homoplastic with a 95% probability (*98*); if applied to mtDNA the extent of the networks has been found to be largely congruent with morphospecies (*109,110*).

Automatic Barcode Gap Discovery (ABGD) was conducted using the ABGD webserver (https://bioinfo.mnhn.fr/abi/public/abgd/abgdweb.html; accessed August, 25th, 2020) with default parameters (i.e., using a relative gap width of 1 and 50 steps, Pmin=0.001, Pmax=0.1) but using Kimura 2-parameter (K2P) distances. ABGD partitions individuals for a range of prior intraspecific distances, instead of using one predefined distance threshold (*100,111*).

### Supplementary Results

#### *Species delimitation analyses*

As expected, the *BPP* analyses showed great influence of the choice of priors for theta and tau on the results (figs S16–S30). In general, theta (population size) has a higher influence than tau (divergence time), with lower theta values leading to the recognition of more species. The splits are often inconsistent between the results of different assembly approaches. The influence of tau, where higher values lead to greater splitting, is lower but for some taxa still noticeable. In *Pleophylla*, for example, lower or intermediate theta priors lead to the splitting of several morphospecies in all assembly approaches (e.g., *P. navicularis, P. nelshoogteensis, P. pseudopilosa*), while a high theta prior generates results mostly agreeing with the previously recognized species limits, except that for some approaches *P. ferruginea* and *P. pilosa* are split into two species each.

The analysis of the effect of the prior on the posterior values of theta and tau shows that, in general, theta values depend on the priors while tau values seemingly do not. In fact, the theta priors seem to have a stronger effect on the tau posteriors than the tau priors do. This may be explained by tau (divergence time) being more or less strictly determined by the data, while theta (population size) is far less so. However, it should be noted that for almost all our datasets the posterior theta values are relatively low (between $10^{-3}$ and $10^{-2}$), meaning that the lowest of our specified theta priors are the most realistic. This contradicts expectations based on comparison with morphospecies, as these priors lead to the highest number of species-level splits.

The exclusion of gaps led to lower support for splits within morphospecies in *BPP* across all datasets. Exclusion of ambiguous sites further increases this effect and reduced over-splitting (figs S10–S12, S16–S24). This may suggest that such sites provide misleading signals that lead to intraspecific splits, also because we had to use the haploid option in *BPP to* analyze potentially heterozygotic data, as the diploid version was too computationally intensive for most of our datasets. However, the effect may be simply caused by loss of resolution due to fewer data. The jackknife analyses often resulted in intermediate posterior probabilities for nodes where the full and reduced datasets led to different results, which suggests that the lesser amount of data only partly explains the discrepancies.

All *tr2* analyses similarly result in over-splitting several morphospecies. However, the details often differ between the two approaches, and no consistent tendency towards more or less splitting relative to *BPP* was observed (figs S10–S15). In contrast to *BPP*, removing gaps leads to more intraspecific splits. This may be explained by *tr2* being based on congruence in gene trees, and removal of gaps leads to removal of highly incomplete gene alignments that give incongruent gene tree topologies.

The *SODA* analyses similarly resulted in over-splitting comparable to *BPP*, suggesting that this algorithm, like the others, does not always reliably distinguish between species- and population-level differentiation (results not shown in detail).

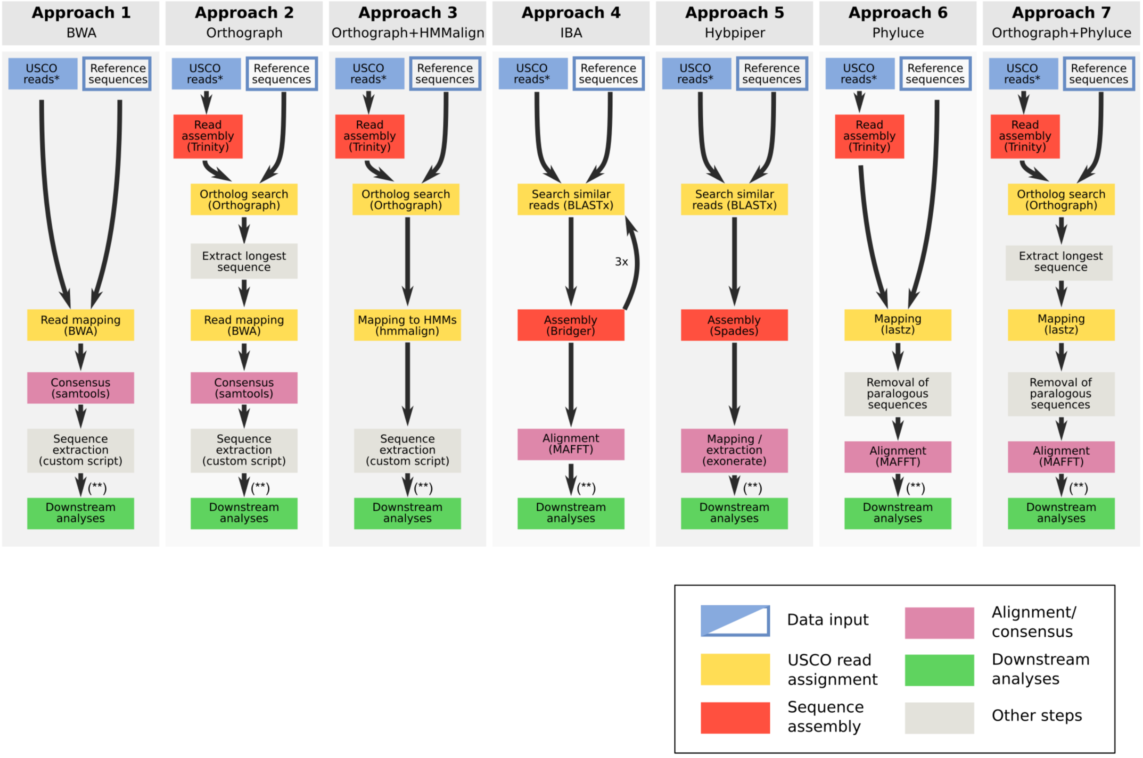


**Fig. S1.**
Pipeline of data assembly for the seven different approaches used.

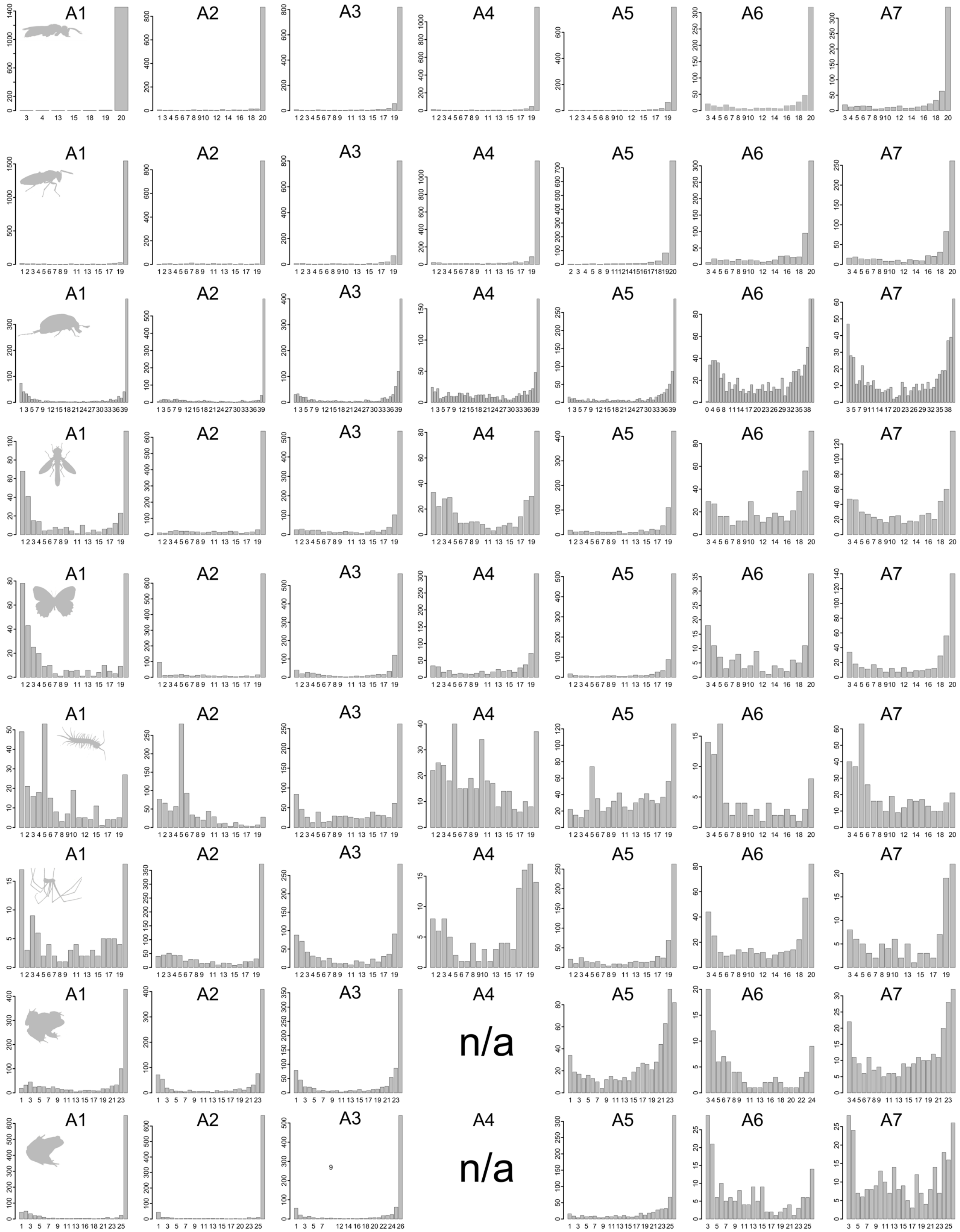

**Fig. S2.**
Distribution of USCOs over the number of specimens in each study case and assembly method.

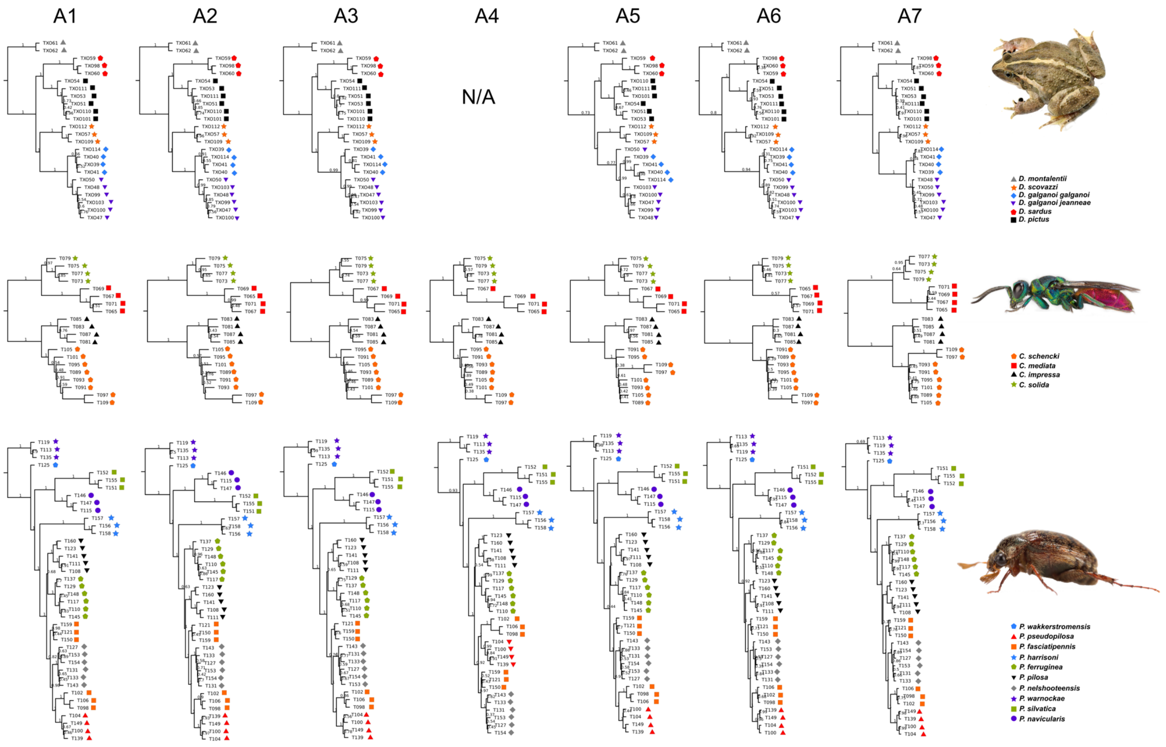


**Fig. S3.**
Tree topology (*ASTRAL*) for study cases (*Discoglossus*, *Chrysis*, *Pleophylla*) and all assembly methods (A1–7) (morphospecies assignment mapped on each terminal).

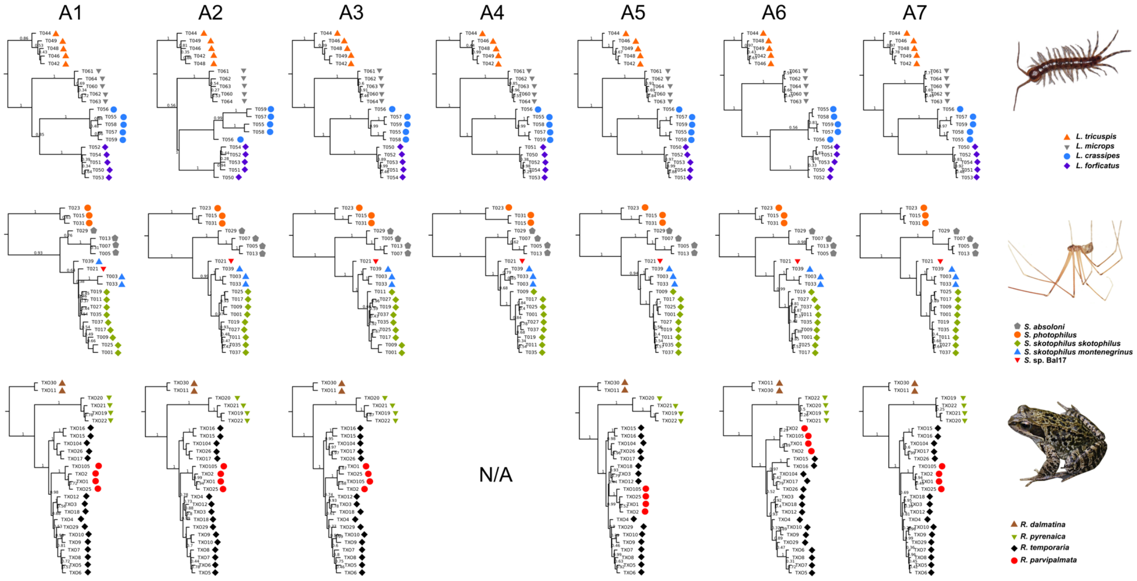


**Fig. S4.**
Tree topology (*ASTRAL*) for study cases (*Lithobius*, *Stygopholcus*, *Rana*) and all assembly methods (A1–7) (morphospecies assignment mapped on each terminal).

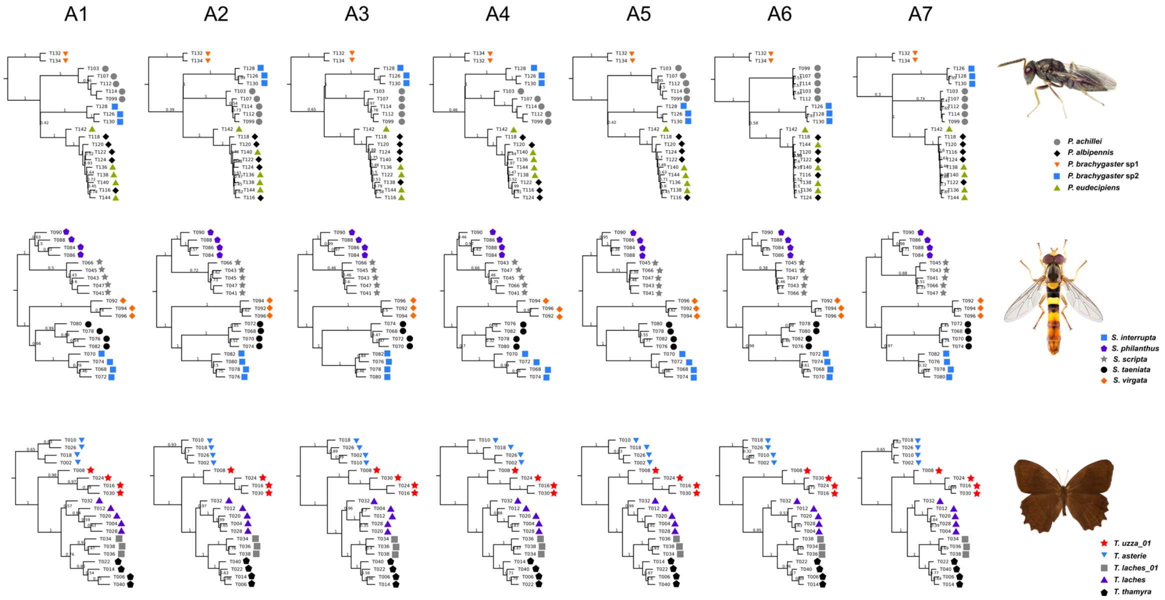


**Fig. S5.**
Tree topology (*ASTRAL*) for study cases (*Pteromalus*, *Sphaerophoria*, *Taygetis*) and all assembly methods (A1–7) (morphospecies assignment mapped on each terminal).

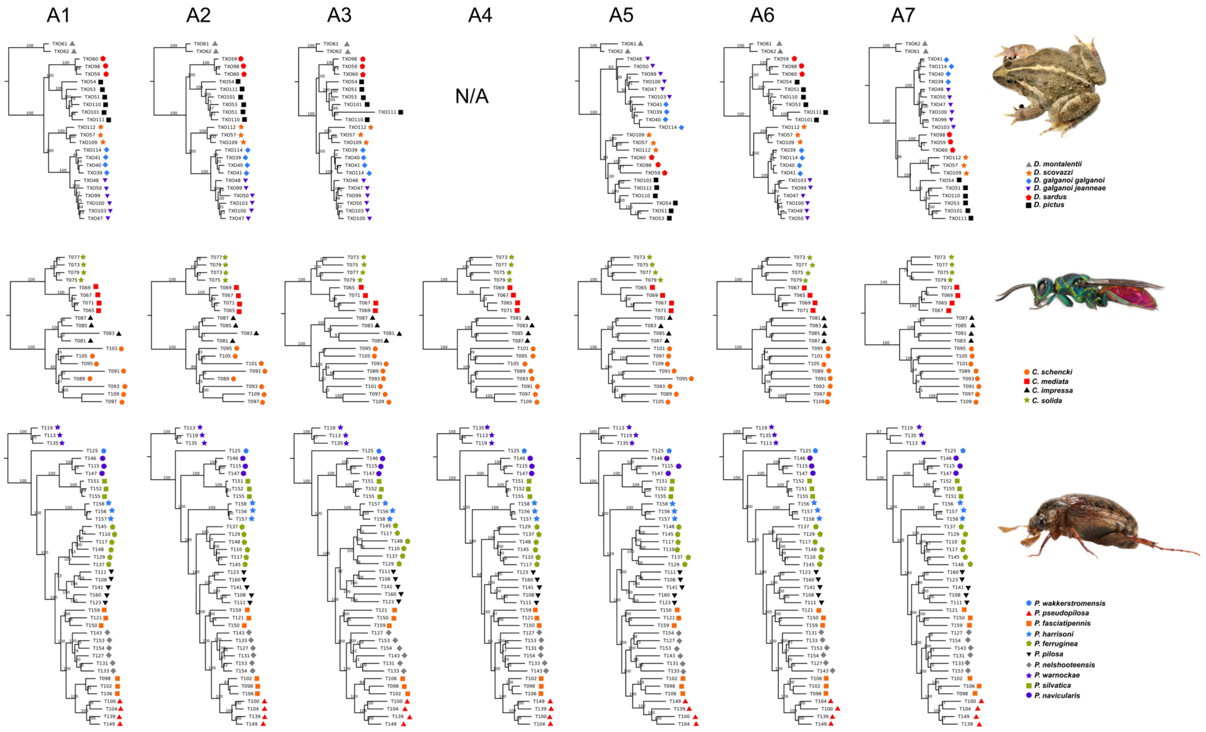


**Fig. S6.**
Tree topology (*IQ-TREE* and concatenated data) for study cases (*Discoglossus*, *Chrysis*, *Pleophylla*) and all assembly methods (A1–7) (morphospecies assignment mapped on each terminal).

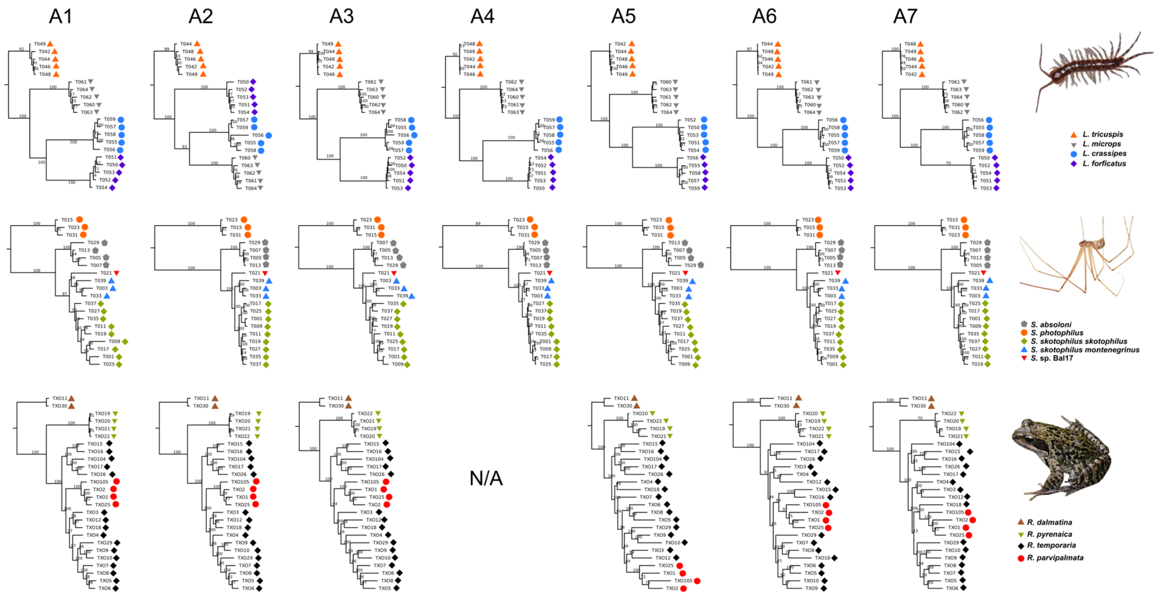


**Fig. S7.**
Tree topology (*IQ-TREE* and concatenated data) for study cases (*Lithobius*, *Stygopholcus*, *Rana*) and all assembly methods (A1–7) (morphospecies assignment mapped on each terminal).

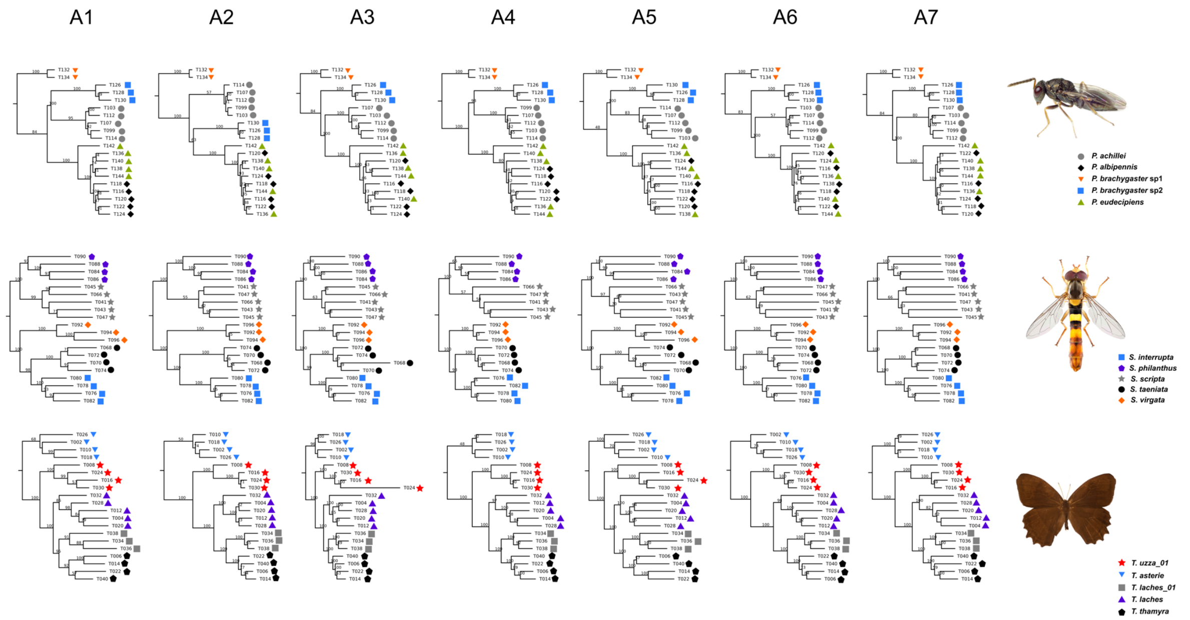


**Fig. S8.**
Tree topology (*IQ-TREE* and concatenated data) for study cases (*Pteromalus*, *Sphaerophoria*, *Taygetis*) and all assembly methods (A1–7) (morphospecies assignment mapped on each terminal).

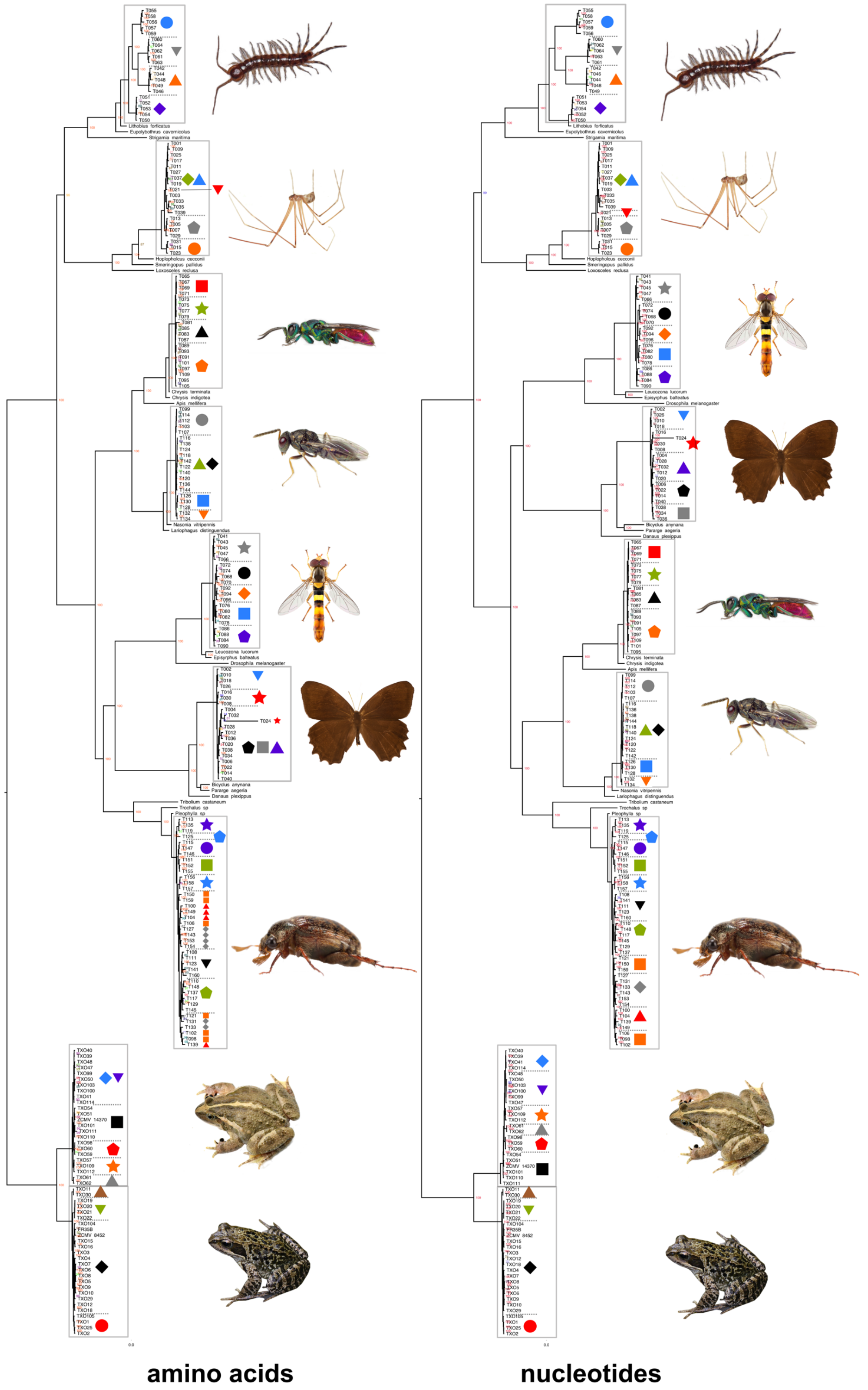


**Fig. S9.**
Maximum likelihood tree based on concatenated data of all taxa (left side, amino acids; right side: nucleotides); study cases in grey boxes, morphospecies indicated by symbols beside terminal labels. Fast bootstrap values are shown close to each branch.

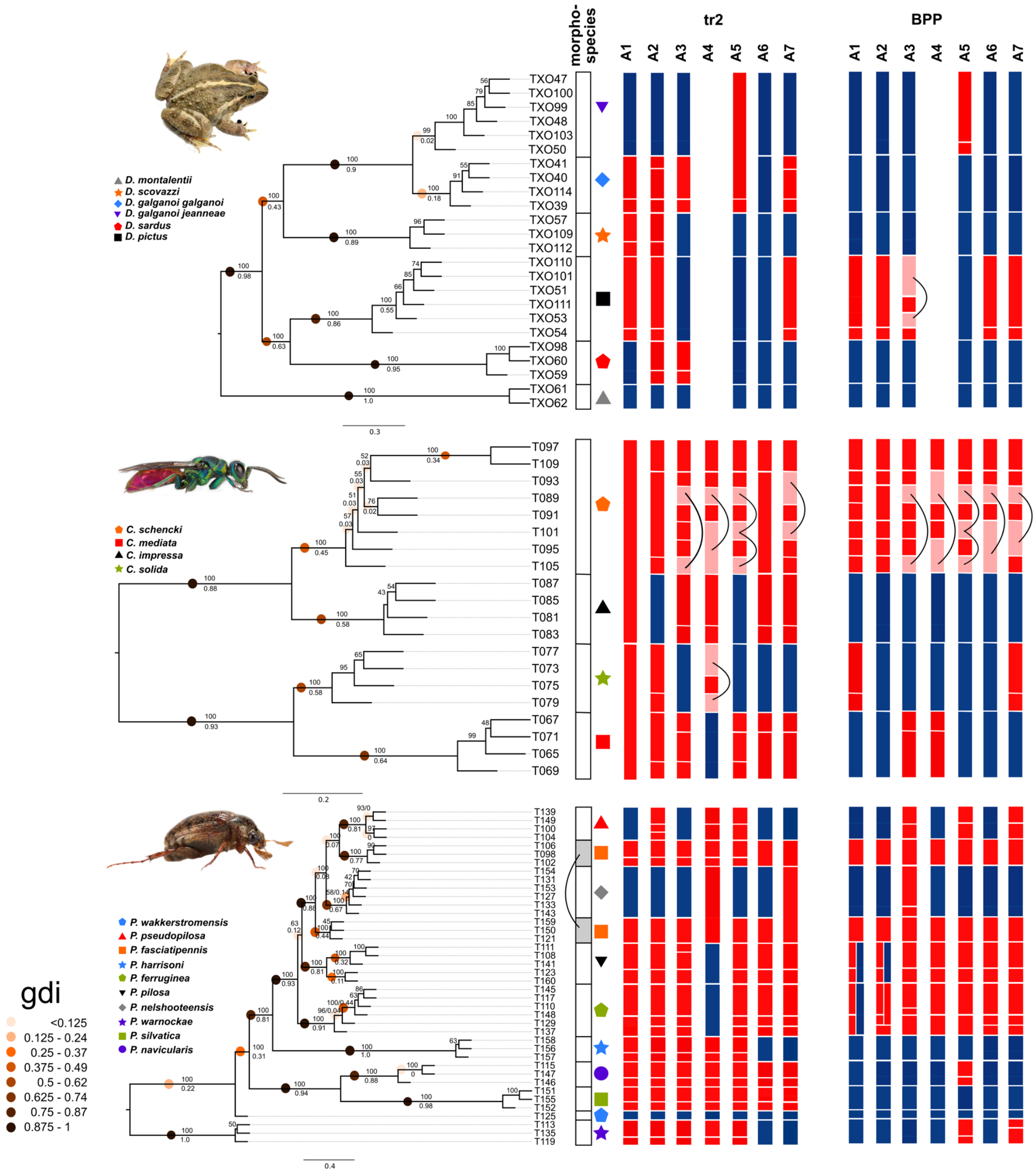


**Fig. S10.**

Results of *tr2* and *BPP** species delimitation for *Discoglossus*, *Chrysis* and *Pleophylla* with reduced USCO data (gaps removed; for the right column in A1 and A2, also ambiguous data were removed), mapped onto all data *ASTRAL* tree (A2) and compared to morphospecies assignments (colored symbols). For *BPP*, results based on median of posterior probabilities for all nine prior combinations are shown. Squares in columns indicate inferred species entities (white: morphospecies; grey: non-monophyletic morphospecies; blue: concordant with morphospecies; red: incongruent with morphospecies; pink: entity not reflected by shown tree topology linked by a bracket, incongruent with morphospecies. Species entities from other assembly approaches may not be monophyletic in this tree because alternative assembly approaches may result in differing starting tree topologies). (* using an *ASTRAL* guide tree obtained from analysis of the respective assembly including all data).

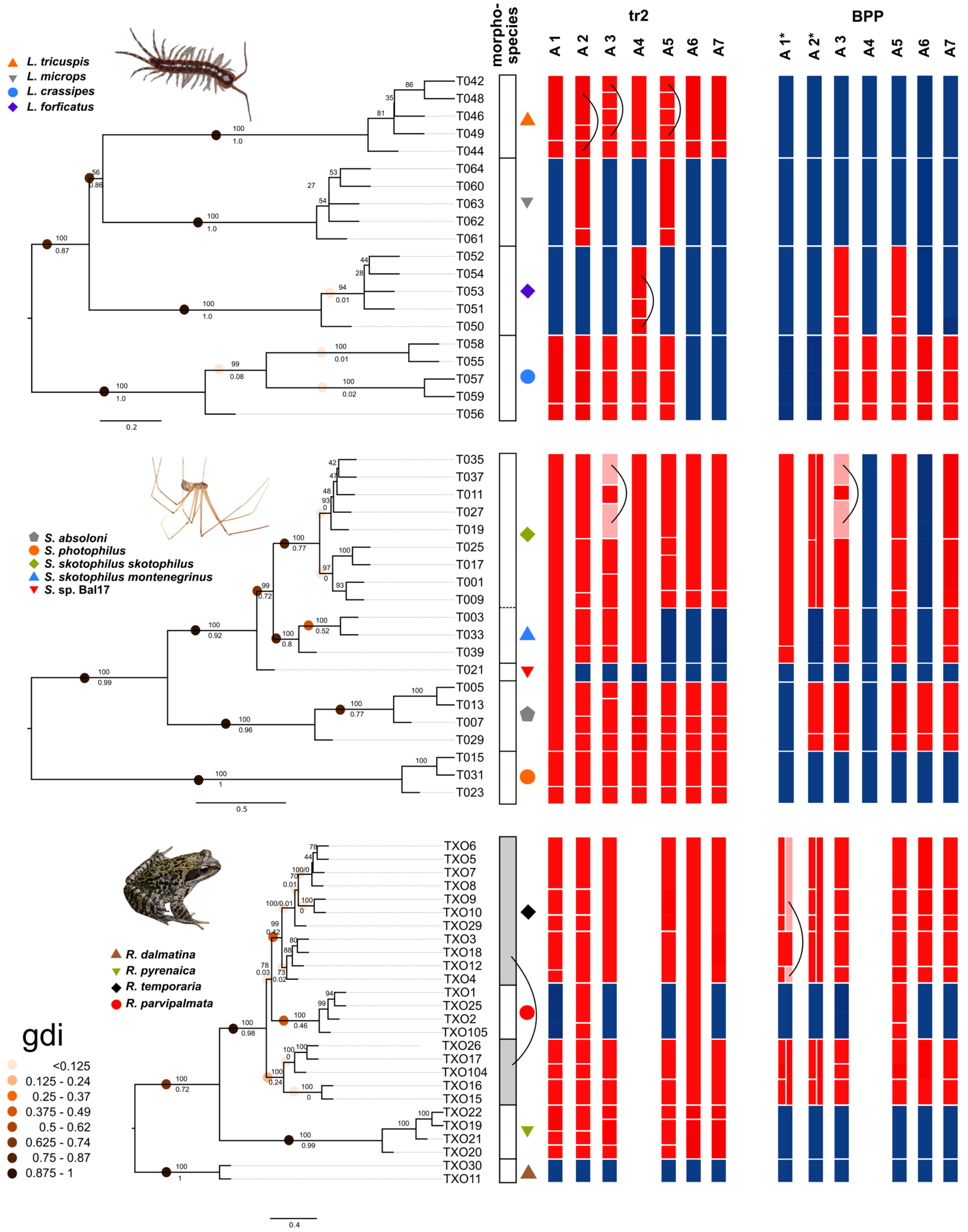


**Fig. S11.** Results of *tr2* and *BPP** species delimitation for *Lithobius*, *Stygopholcus*, and *Rana* with reduced USCO data (gaps removed; for the right column in A1 and A2, also ambiguous data were removed), mapped onto all data *ASTRAL* tree (A2) and compared to morphospecies assignments (colored symbols). For *BPP*, results based on median of posterior probabilities for all nine prior combinations are shown. Columns showing species entities colored as in Fig. S3. (* using *ASTRAL* tree obtained from analysis of the respective assembly including all data).

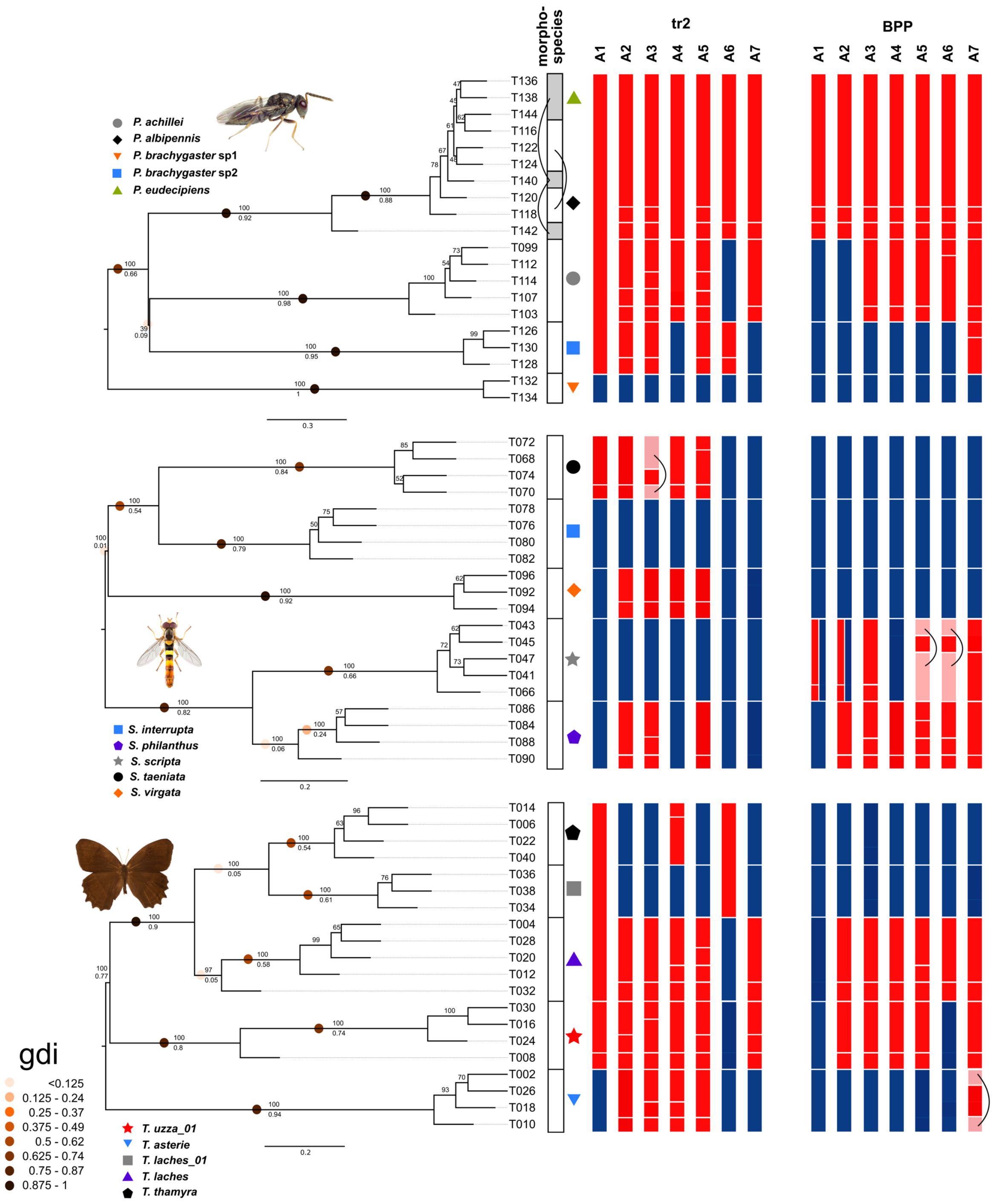


**Fig. S12.** Results of *tr2* and *BPP** species delimitation for *Pteromalus, Sphaerophoria,* and *Taygetis* with reduced USCO data (gaps removed; for the right column in A1 and A2, also ambiguous data were removed), mapped onto all data *ASTRAL* tree (A2) and compared to morphospecies assignments (colored symbols). For *BPP*, results based on median of posterior probabilities for all nine prior combinations are shown. Columns showing species entities colored as in Fig. S3. (* using *ASTRAL* tree obtained from analysis of the respective assembly including all data).

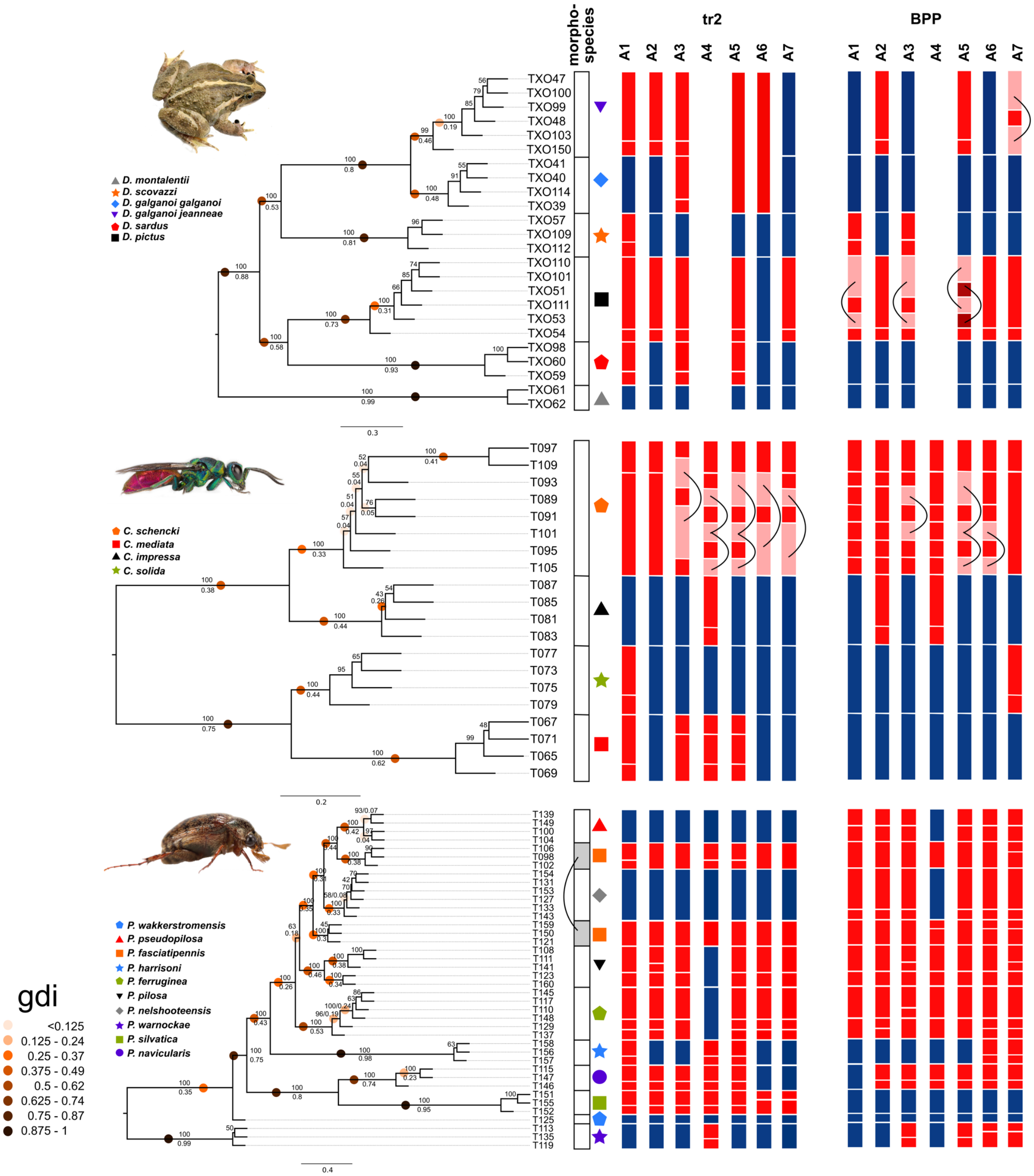


**Fig. S13.**
Results of *tr2* and *BPP* species delimitation using all USCO data for *Discoglossus*, *Chrysis* and *Pleophylla*, mapped onto *ASTRAL* tree from A2, compared to morphospecies assignments (colored symbols). For *BPP*, results based on median of posterior probabilities for all nine prior combinations are shown. Columns showing species entities colored as in Fig. S3. On branches *ASTRAL* support value as well as genealogical divergence index (gdi) of A2 tree is shown. Colored dots on selected nodes indicate gdi values according to legend for better visual illustration.

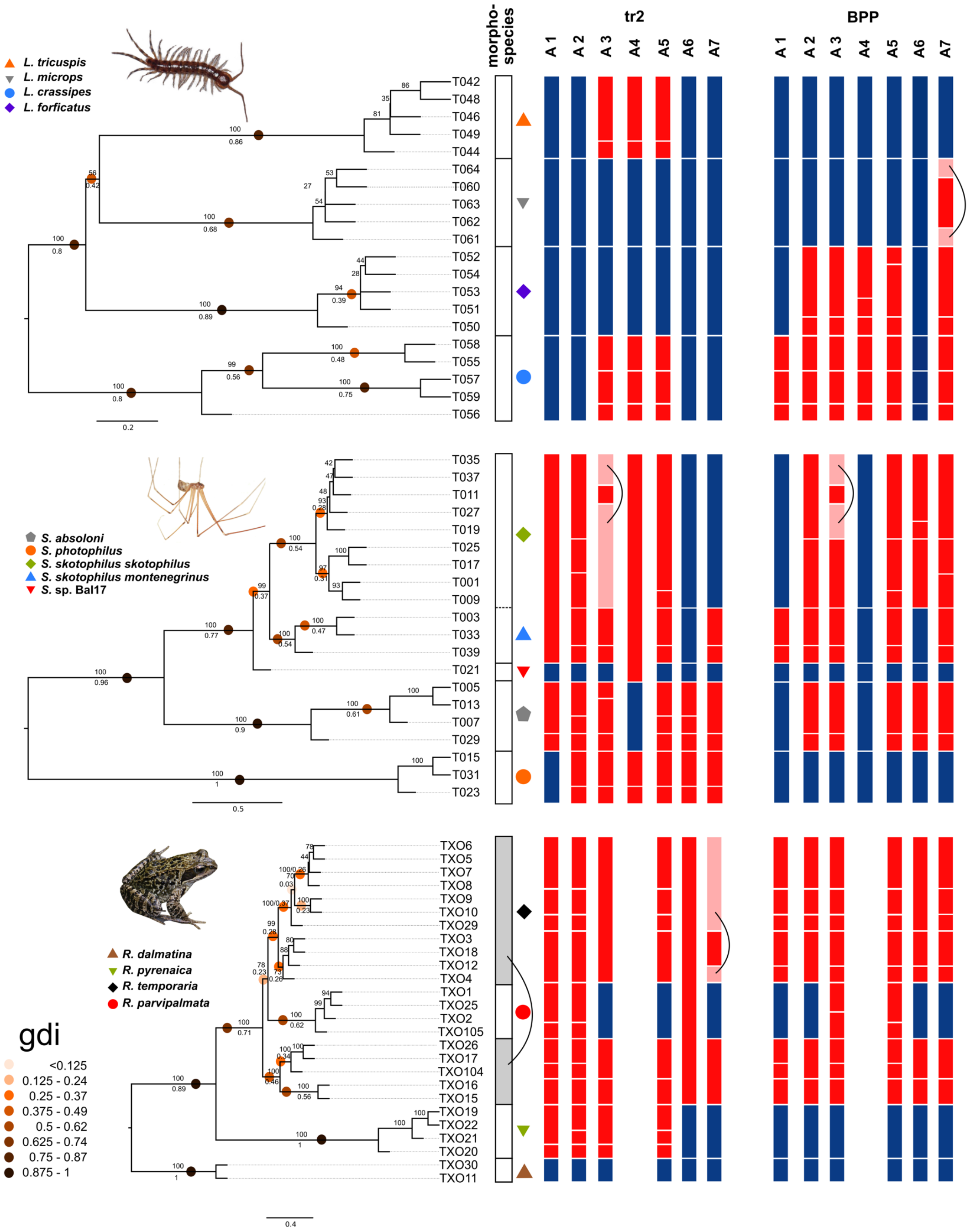


**Fig. S14.**

Results of *tr2* and *BPP* species delimitation using all USCO data for *Lithobius*, *Stygopholcus*, and *Rana*, mapped onto *ASTRAL* tree from A2, compared to morphospecies assignments (colored symbols). For *BPP*, results based on median of posterior probabilities for all nine prior combinations are shown. Columns showing species entities colored as in Fig. S3. On branches *ASTRAL* support value as well as genealogical divergence index (gdi) of A2 tree is shown. Colored dots on selected nodes indicate gdi values according to legend for better visual illustration.

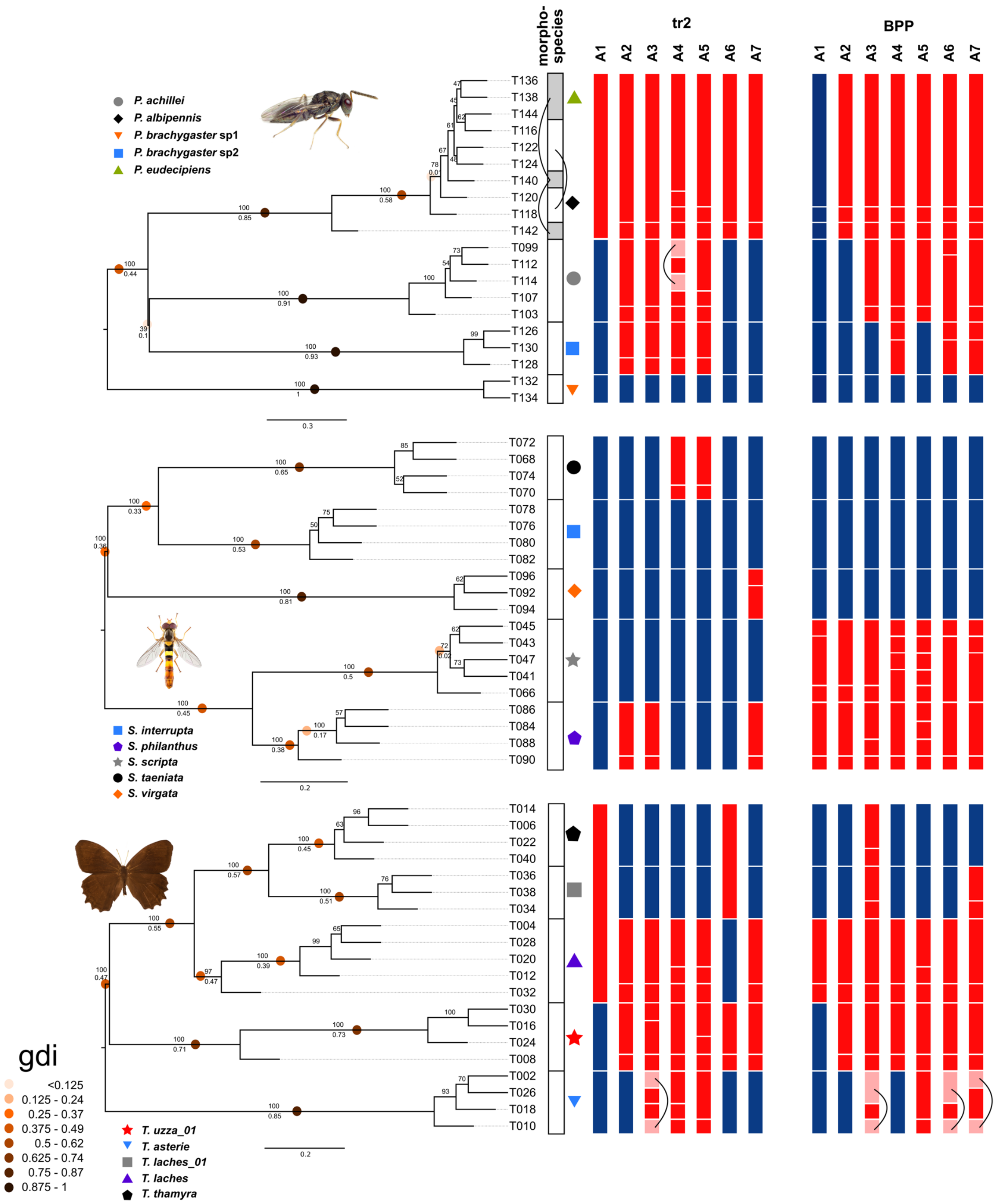


**Fig. S15.**
Results of *tr2* and *BPP* species delimitation using all USCO data for *Pteromalus, Sphaerophoria,* and *Taygetis*, mapped onto *ASTRAL* tree from A2, compared to morphospecies assignments (colored symbols). For *BPP*, results based on median of posterior probabilities for all nine prior combinations are shown. Columns showing species entities colored as in Fig. S3. On branches *ASTRAL* support value as well as genealogical divergence index (gdi) of A2 tree is shown. Colored dots on selected nodes indicate gdi values according to legend for better visual illustration.

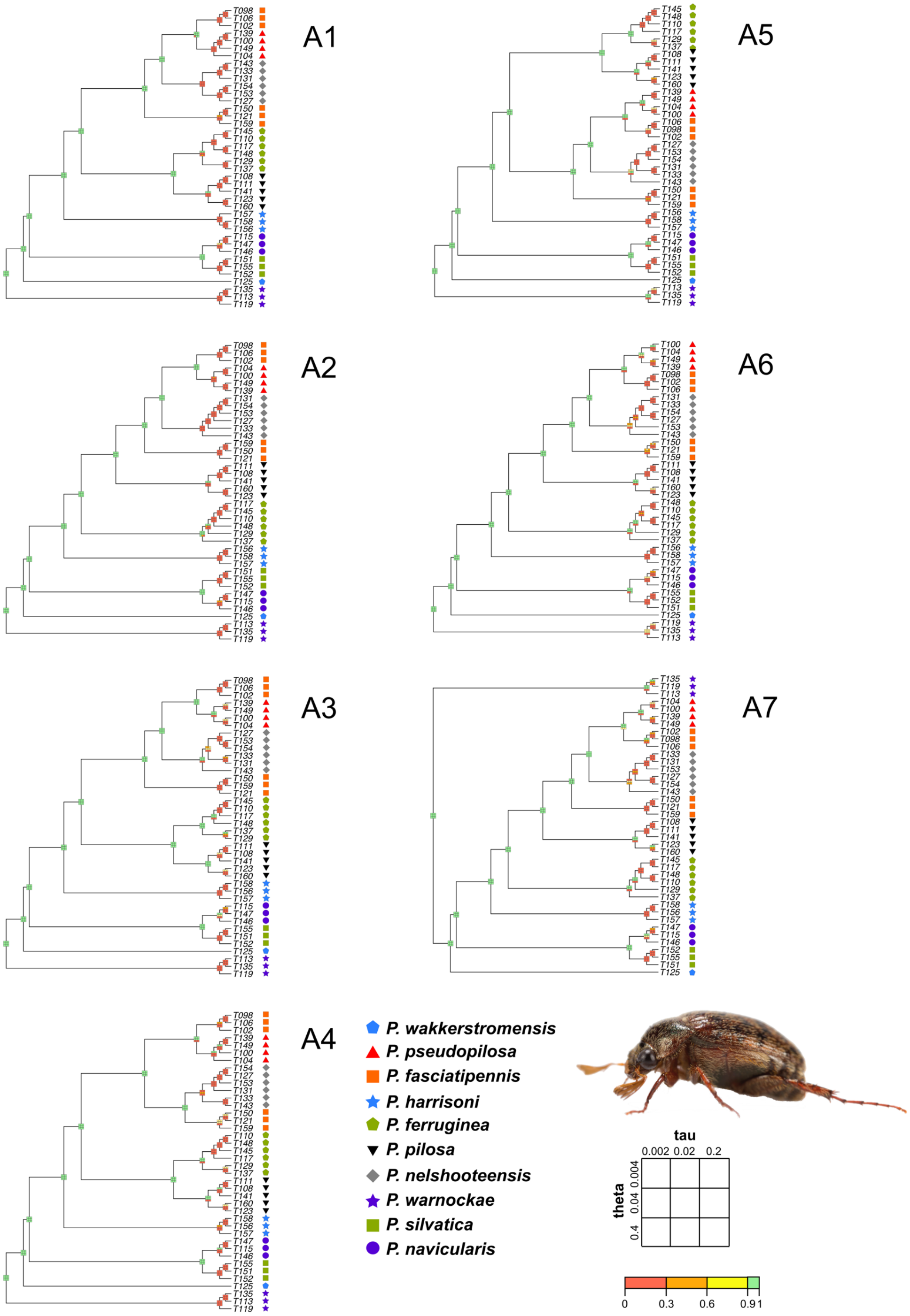


**Fig. S16.**
Results of *BPP* species delimitation with gaps (and ambiguous data, A1 and A2) removed using the all data *ASTRAL* tree as guide tree and different prior combinations for tau and theta for *Pleophylla*, mapped onto respective *ASTRAL* tree from A1–7, compared to morphospecies assignments (shown by symbols associated to each individual terminal).

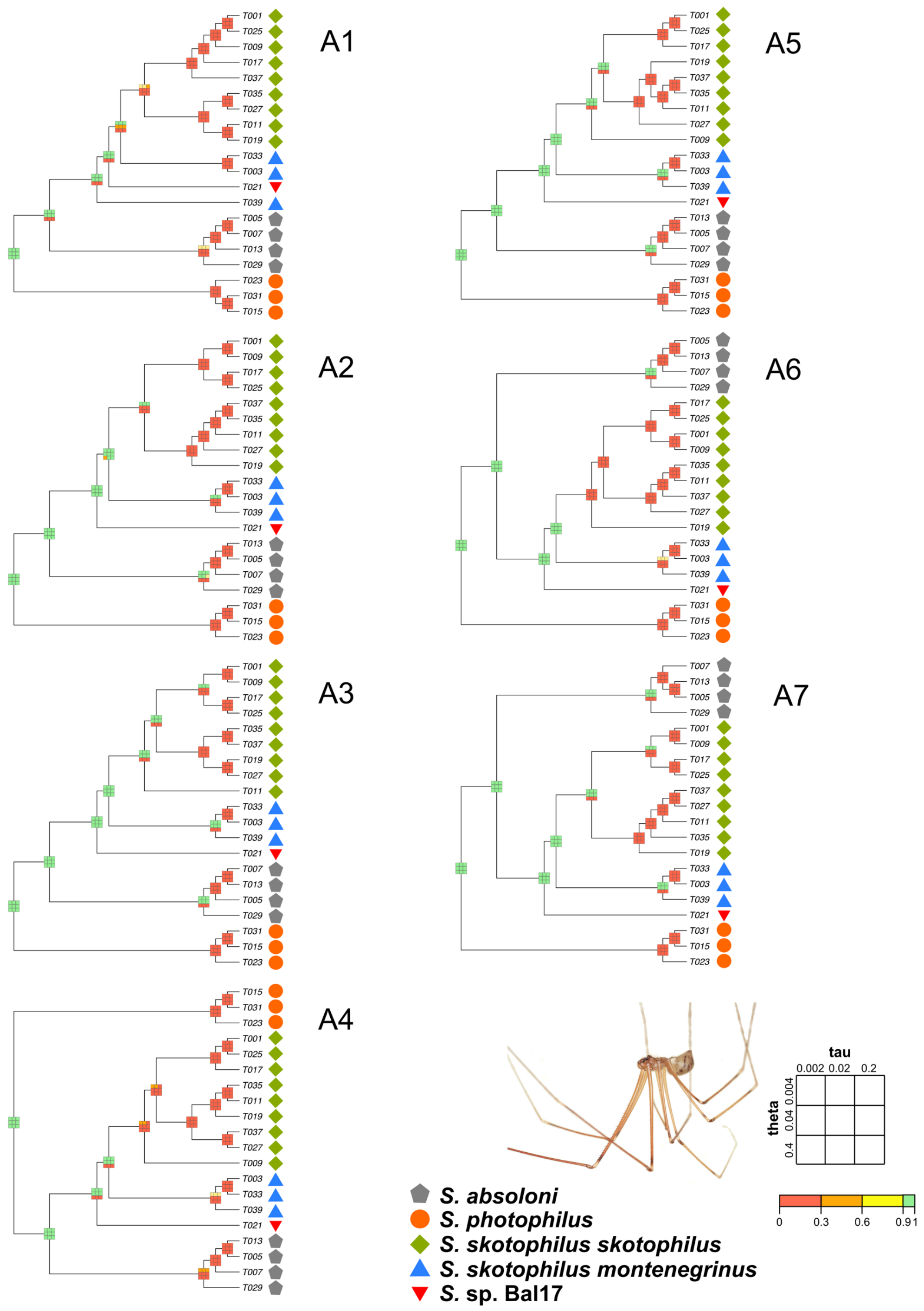


**Fig. S17.**
Results of *BPP* species delimitation with gaps (and ambiguous data, A1 and A2) removed using the all data *ASTRAL* tree as guide tree and different prior combinations for tau and theta for *Stygopholcus*, mapped onto respective *ASTRAL* tree from A1–7, compared to morphospecies assignments (shown by symbols associated to each individual terminal).

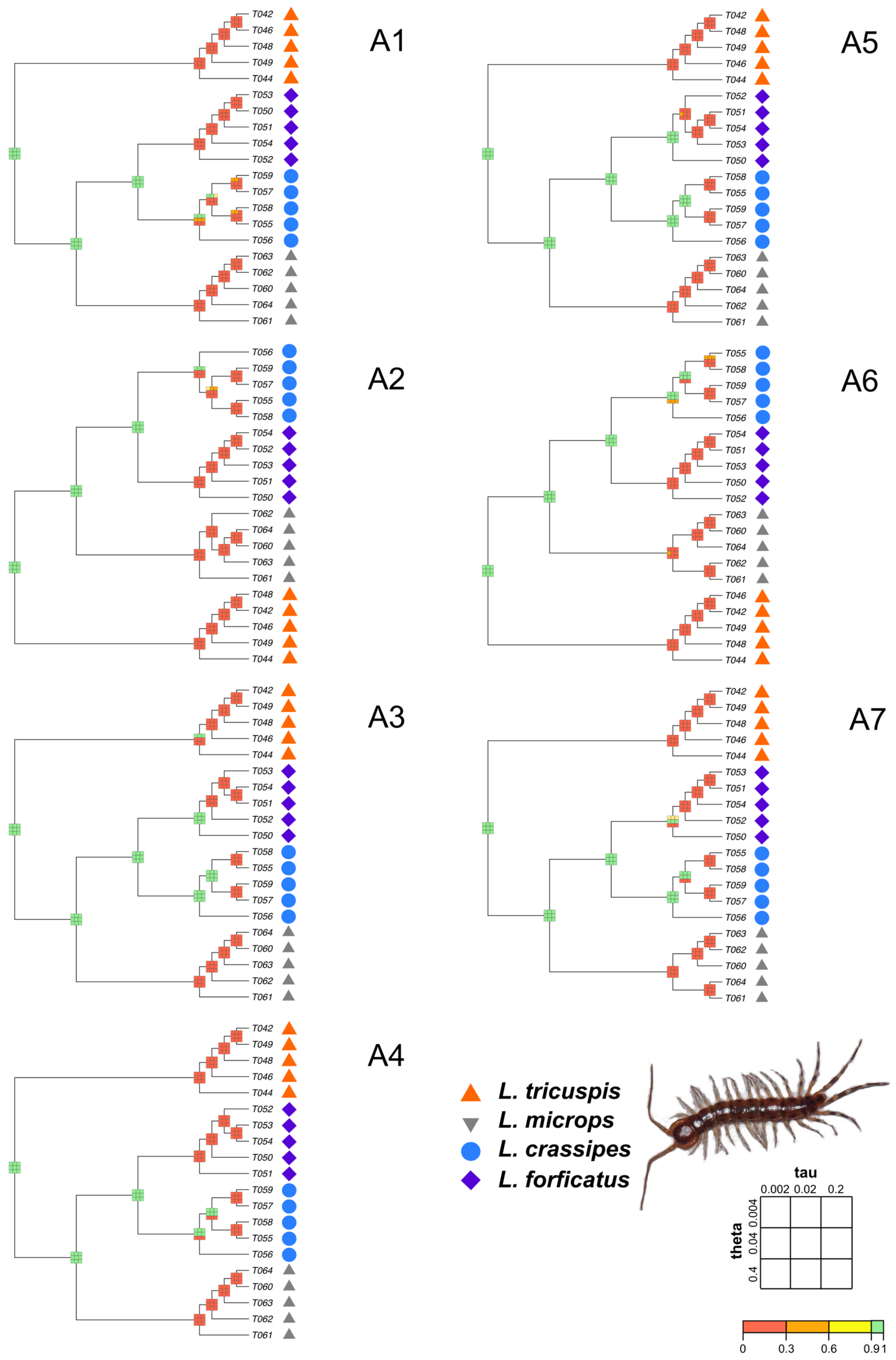


**Fig. S18.**
Results of *BPP* species delimitation with gaps (and ambiguous data, A1 and A2) removed using the all data *ASTRAL* tree as guide tree and different prior combinations for tau and theta for *Lithobius*, mapped onto respective *ASTRAL* tree from A1–7, compared to morphospecies assignments (shown by symbols associated to each individual terminal).

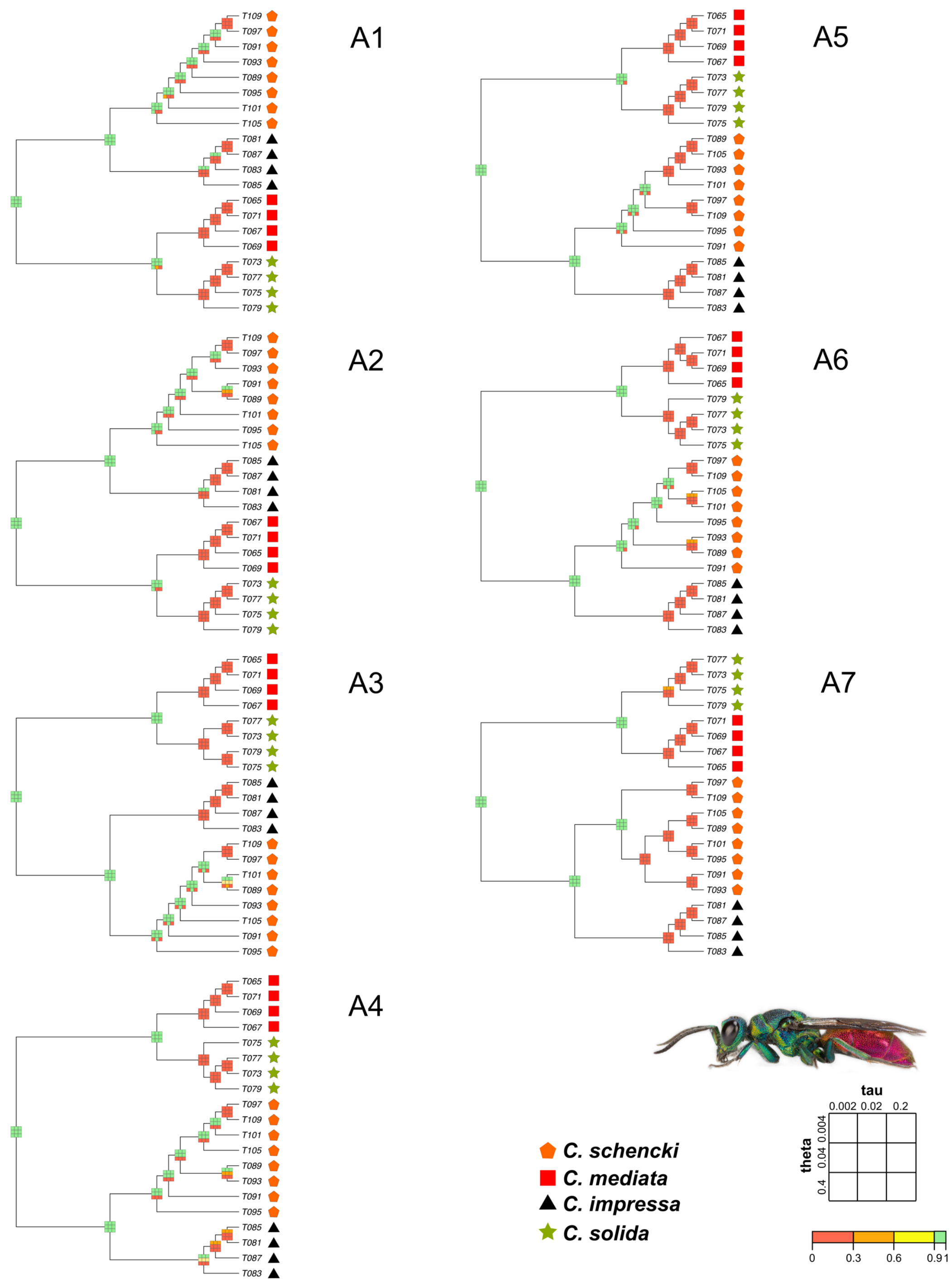


**Fig. S19.**
Results of *BPP* species delimitation with gaps (and ambiguous data, A1 and A2) removed using the all data *ASTRAL* tree as guide tree and different prior combinations for tau and theta for *Chrysis*, mapped onto respective *ASTRAL* tree from A1–7, compared to morphospecies assignments (shown by symbols associated to each individual terminal).

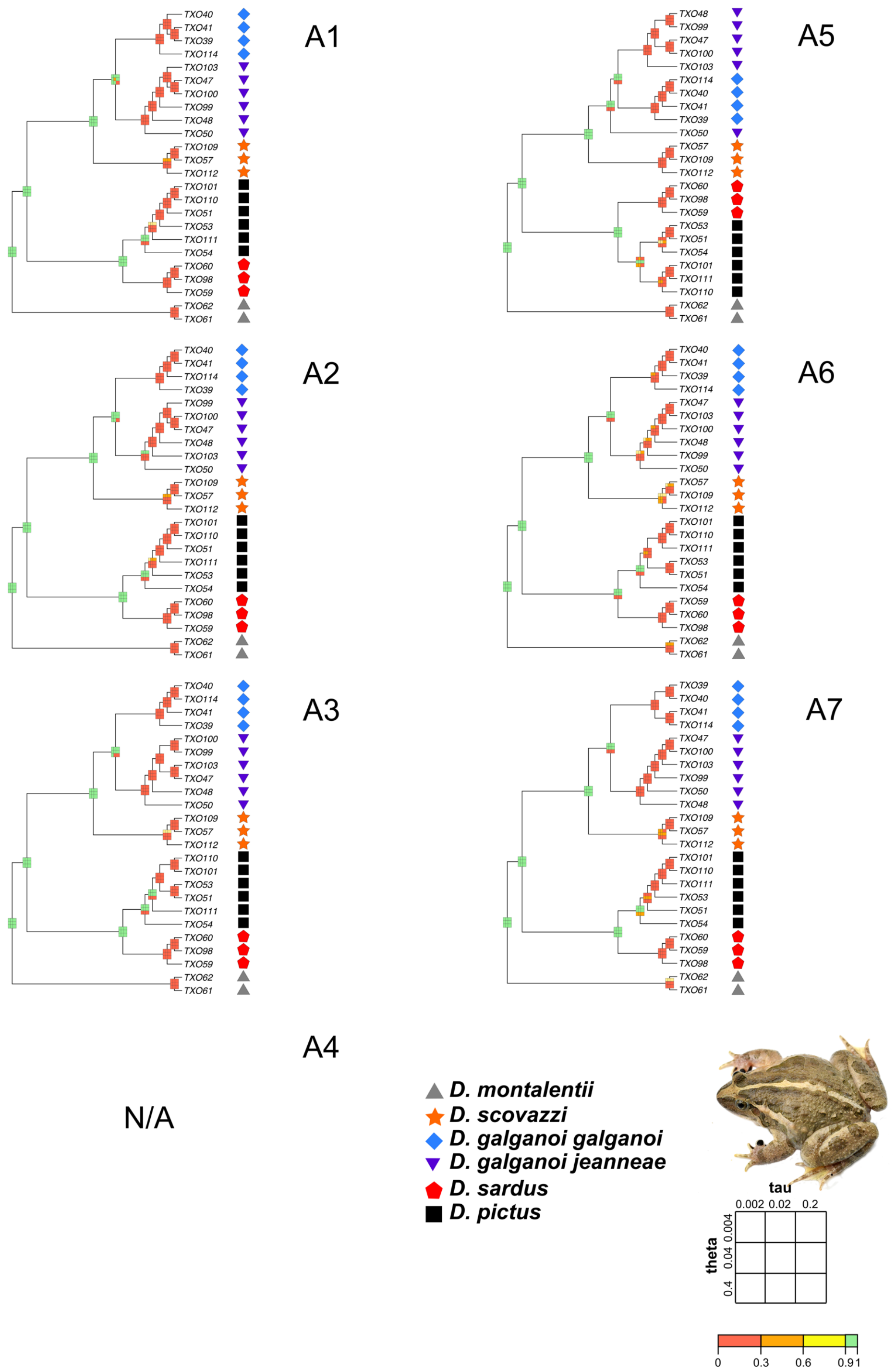


**Fig. S20.**
Results of *BPP* species delimitation with gaps (and ambiguous data, A1 and A2) removed using the all data *ASTRAL* tree as guide tree and different prior combinations for tau and theta for *Discoglossus*, mapped onto respective *ASTRAL* tree from A1–7, compared to morphospecies assignments (shown by symbols associated to each individual terminal).

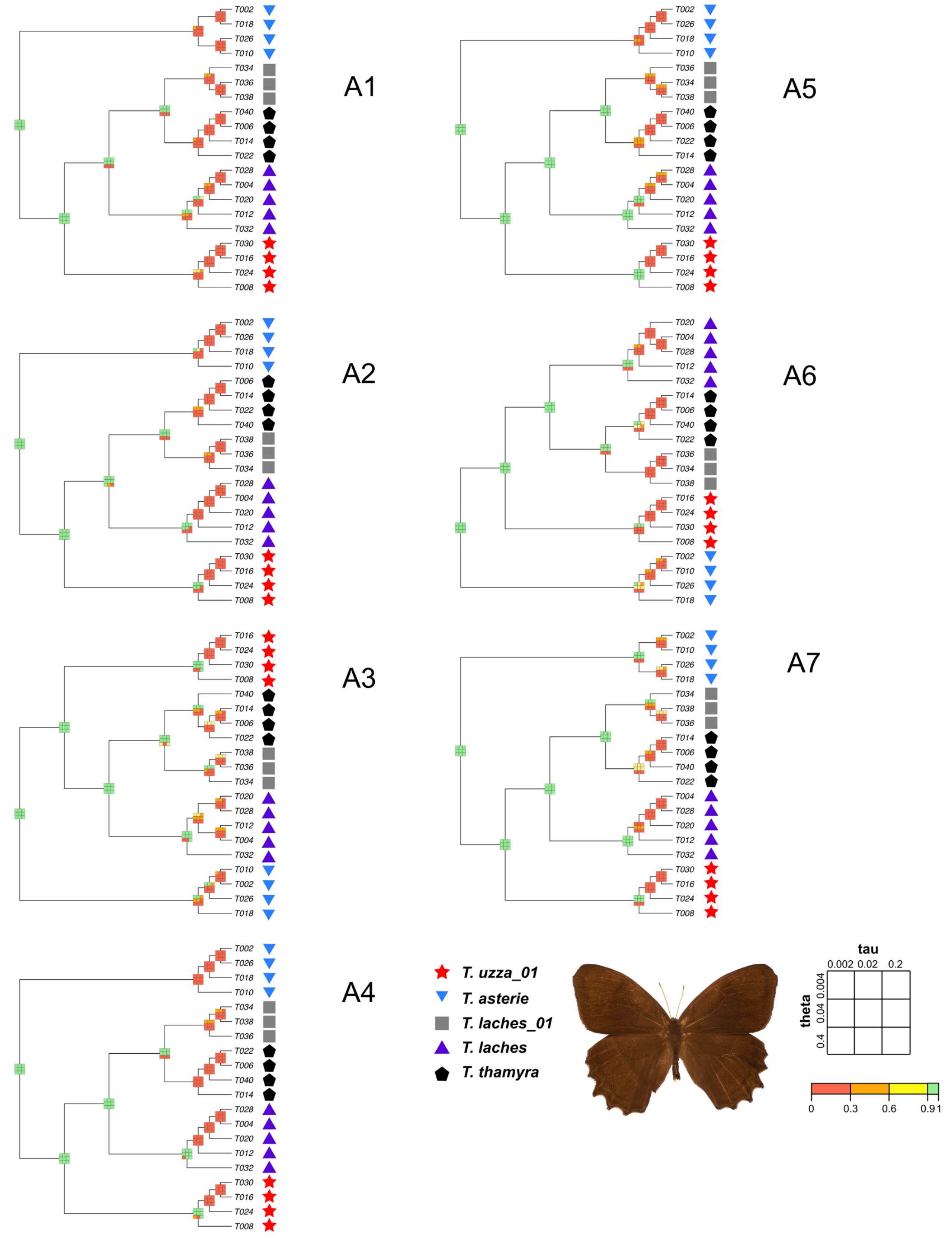


**Fig. S21.**
Results of *BPP* species delimitation with gaps (and ambiguous data, A1 and A2) removed using the all data *ASTRAL* tree as guide tree and different prior combinations for tau and theta for *Taygetis*, mapped onto respective *ASTRAL* tree from A1–7, compared to morphospecies assignments (shown by symbols associated to each individual terminal).

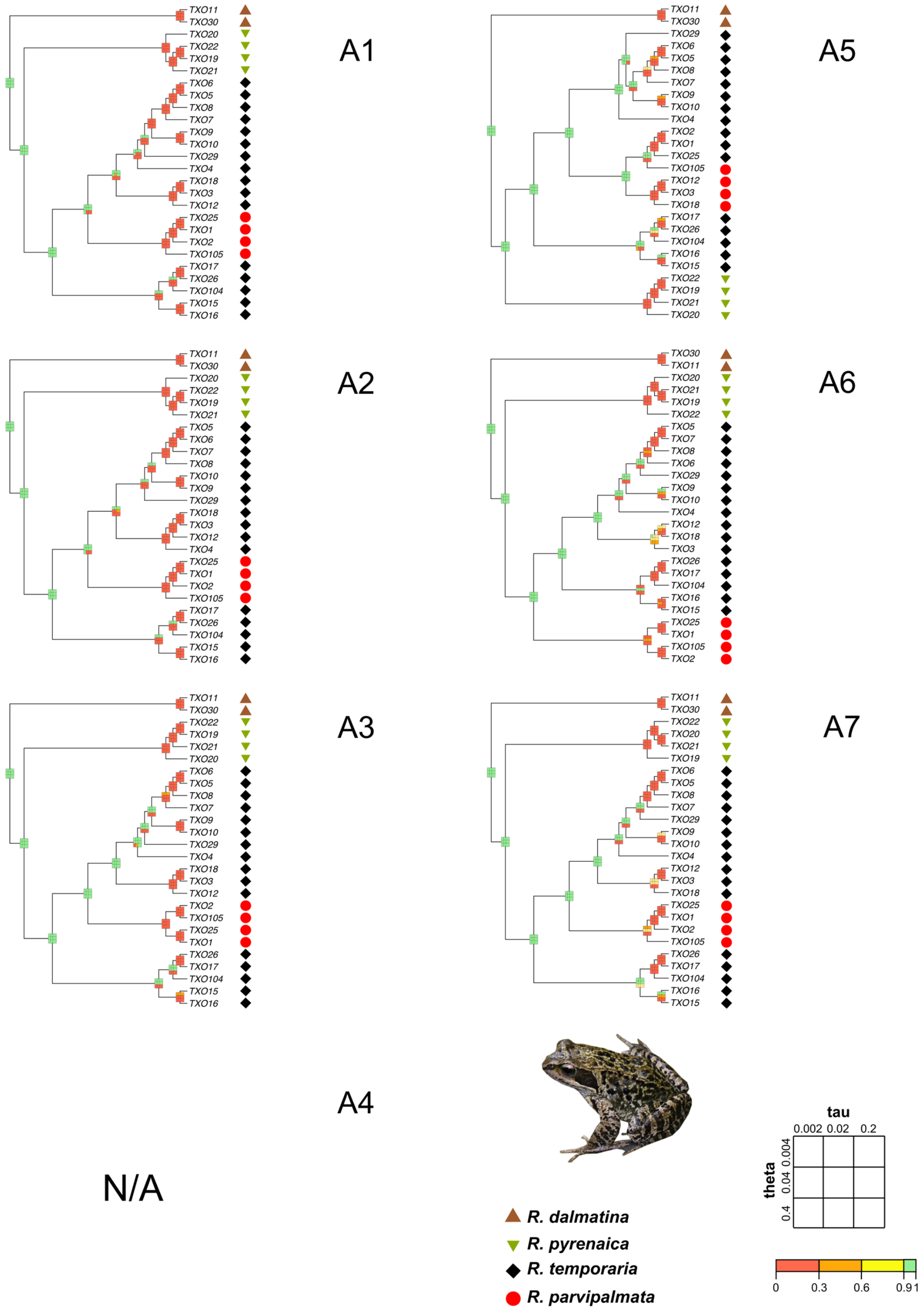


**Fig. S22.**
Results of *BPP* species delimitation with gaps (and ambiguous data, A1 and A2) removed using the all data *ASTRAL* tree as guide tree and different prior combinations for tau and theta for *Rana*, mapped onto respective *ASTRAL* tree from A1–7, compared to morphospecies assignments (shown by symbols associated to each individual terminal).

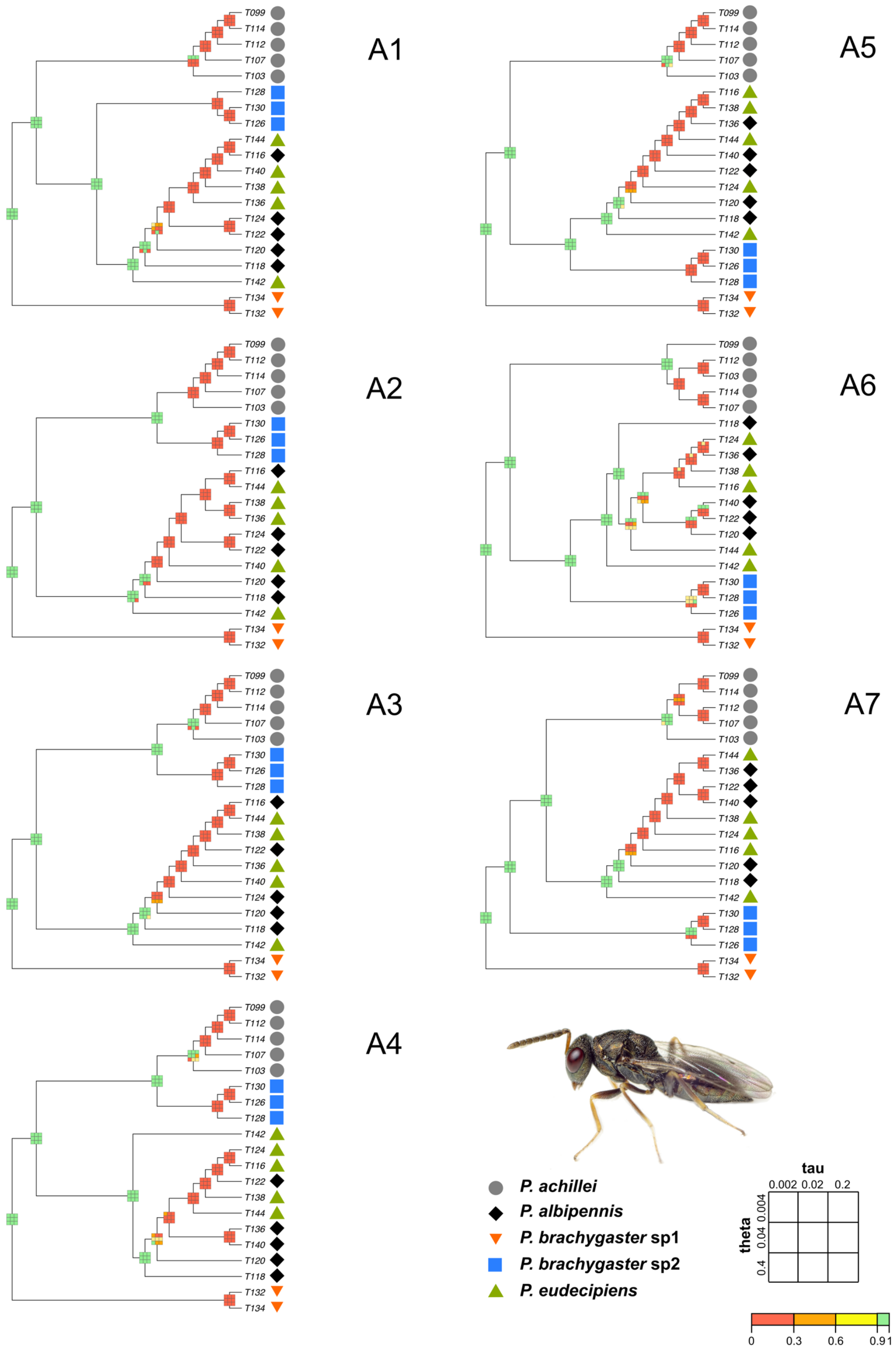


**Fig. S23.**
Results of *BPP* species delimitation with gaps (and ambiguous data, A1 and A2) removed using the all data *ASTRAL* tree as guide tree and different prior combinations for tau and theta for *Pteromalus*, mapped onto respective *ASTRAL* tree from A1–7, compared to morphospecies assignments (shown by symbols associated to each individual terminal).

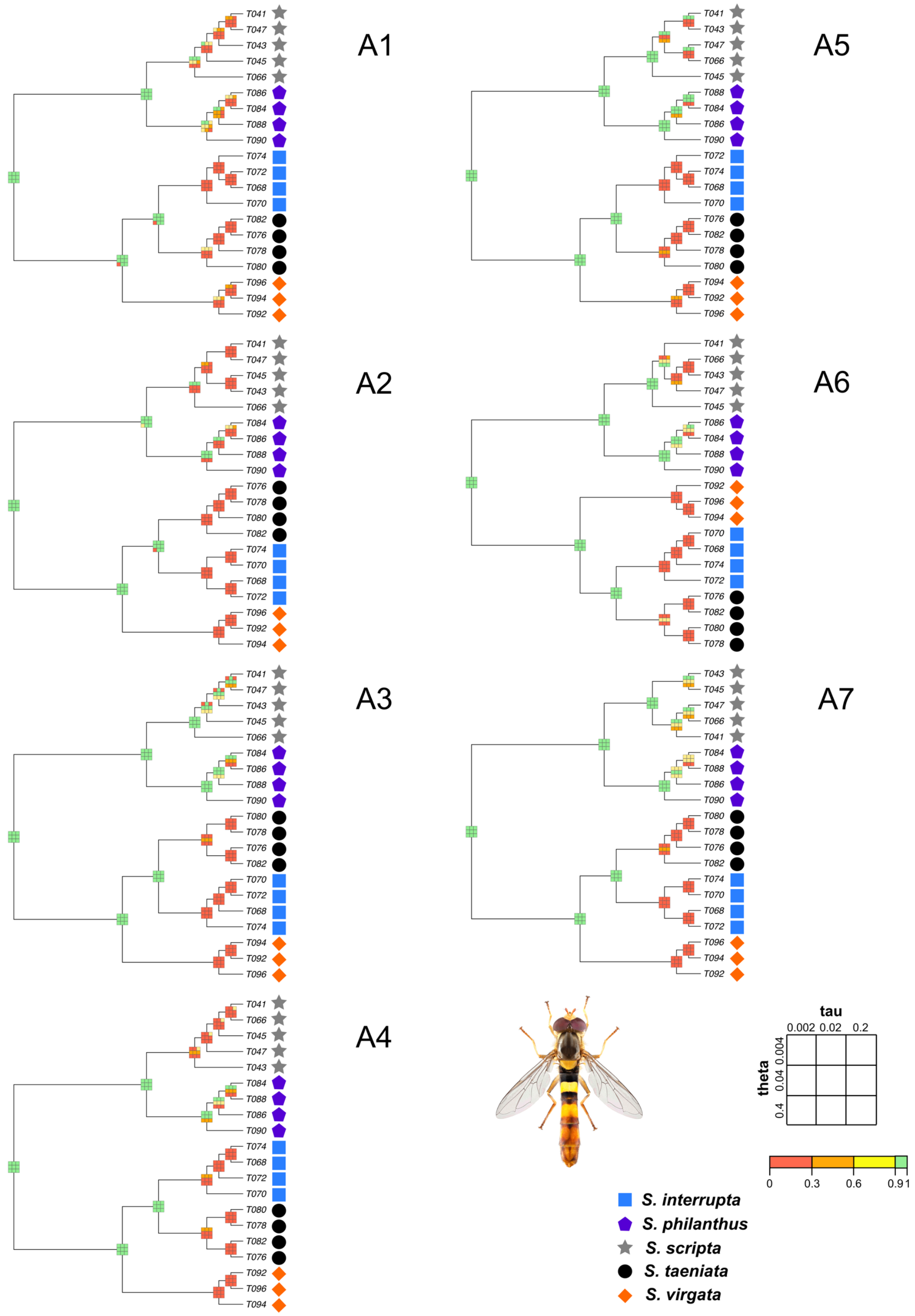


**Fig. S24.**
Results of *BPP* species delimitation with gaps (and ambiguous data, A1 and A2) removed using the all data *ASTRAL* tree as guide tree and different prior combinations for tau and theta for *Sphaerophoria*, mapped onto respective *ASTRAL* tree from A1–7, compared to morphospecies assignments (shown by symbols associated to each individual terminal).

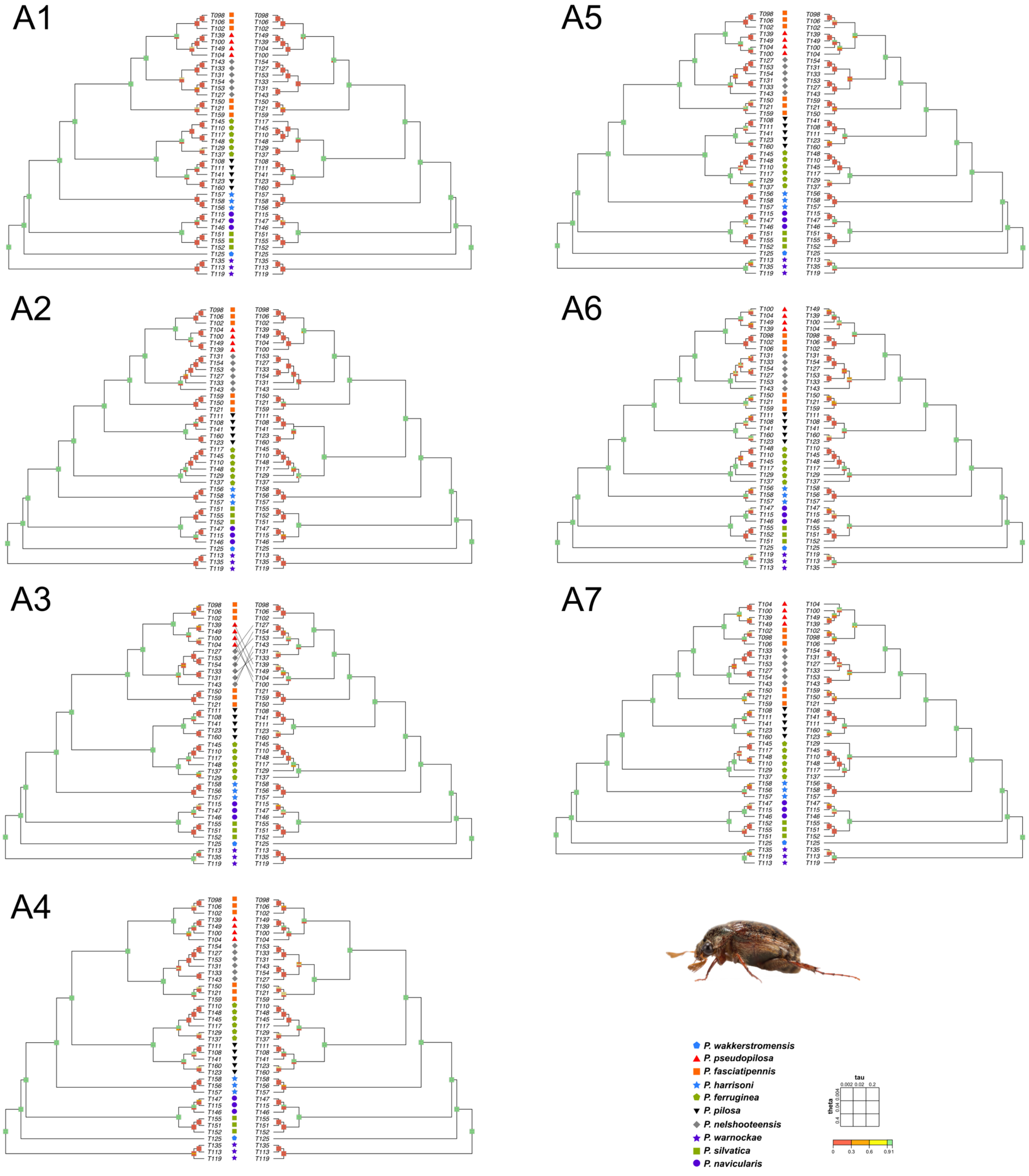


**Fig. S25.**
Results of *BPP* species delimitation with all USCO data (left tree) and reduced USCO data* (gaps (and ambiguous data, A1 and A2) removed) (right tree) (*with an *ASTRAL* guide tree inferred from these data) using different prior combinations for tau and theta for *Pleophylla*, mapped onto respective *ASTRAL* tree from A1–7, compared to morphospecies assignments (shown by symbols associated to each individual terminal).

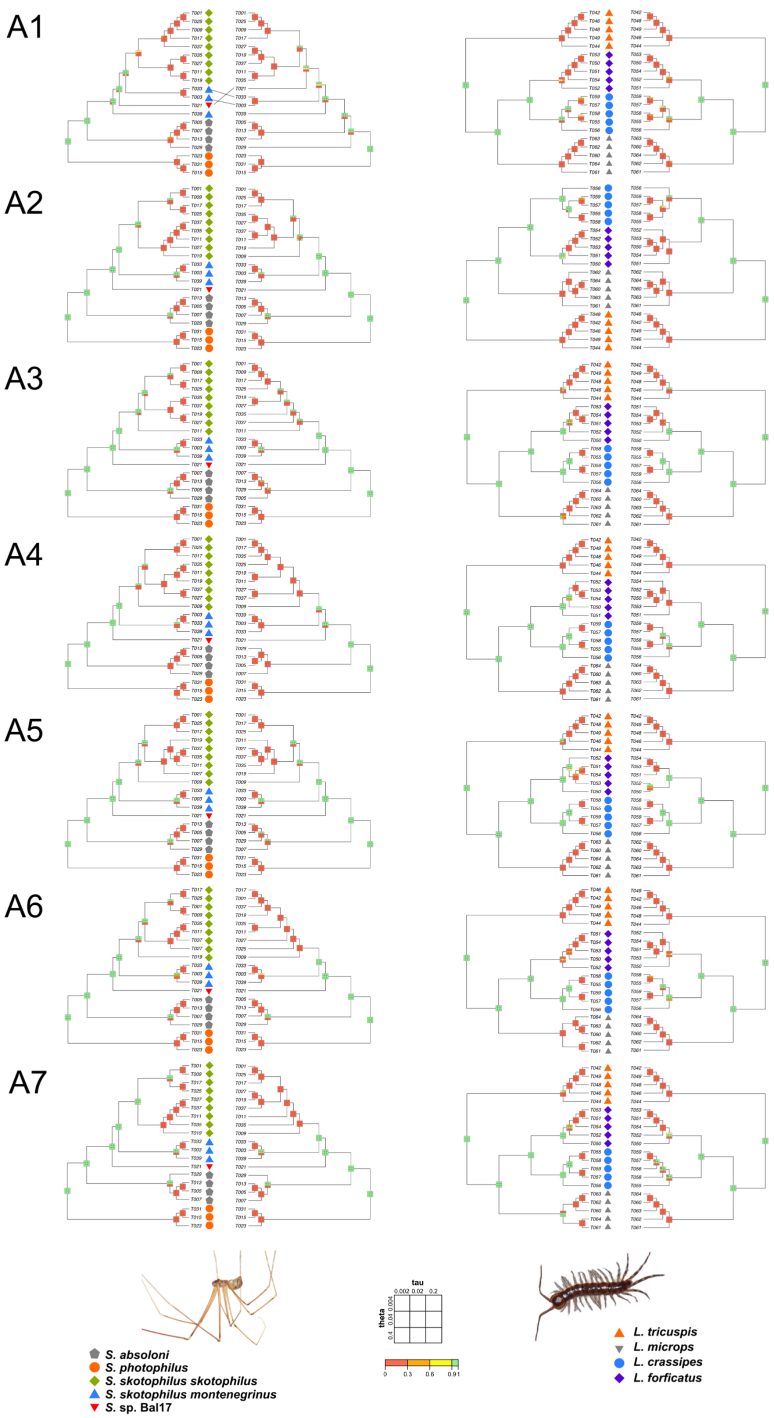


**Fig. S26.**
Results of *BPP* species delimitation with all USCO data (left tree) and reduced USCO data* (right tree; gaps (and ambiguous data, A1 and A2) removed) (*with an *ASTRAL* guide tree inferred from these data) using different prior combinations for tau and theta for *Lithobius* and *Stygopholcus*, mapped onto respective *ASTRAL* tree from A1–7, compared to morphospecies assignments (shown by symbols associated to each individual terminal).

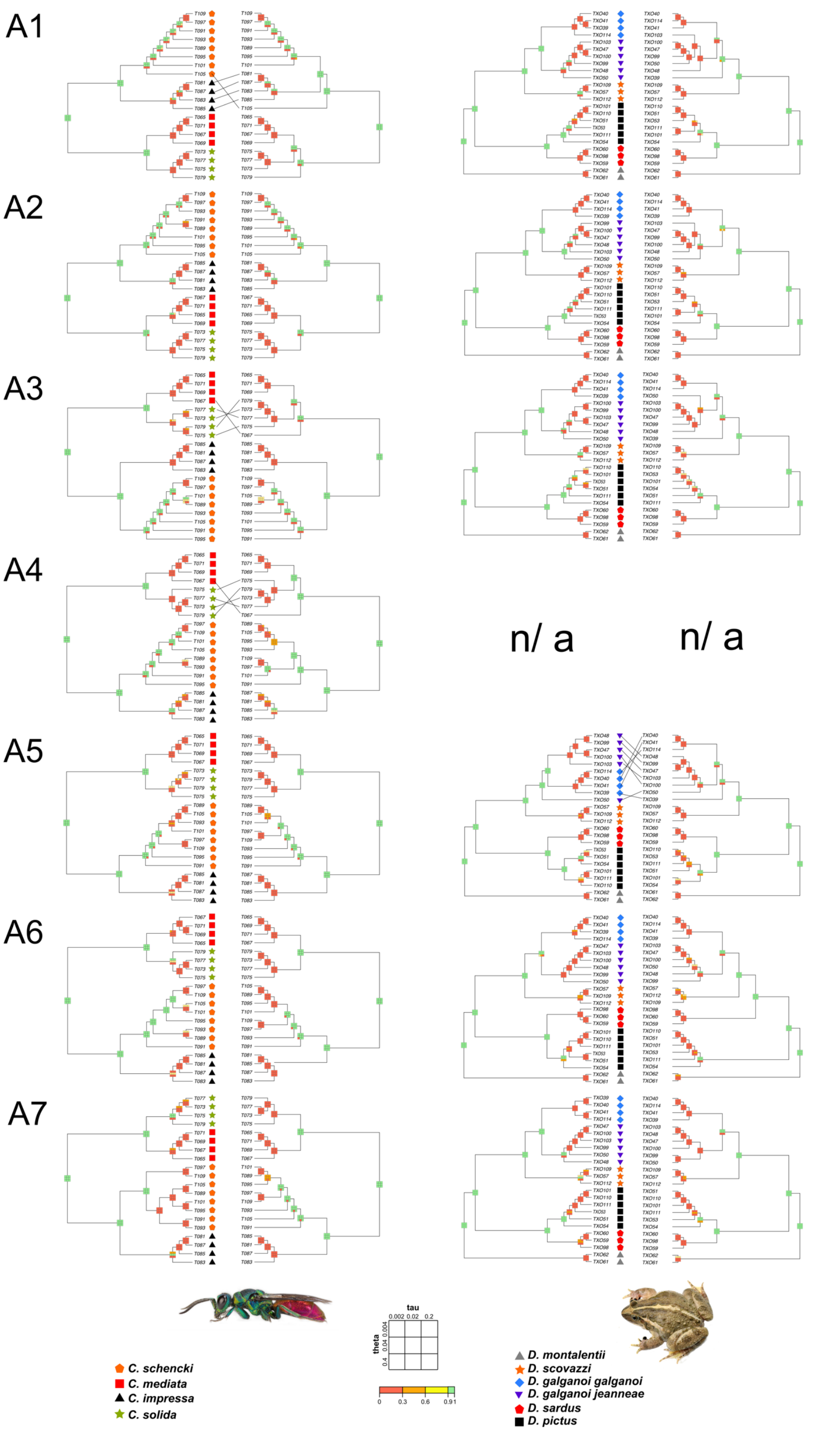


**Fig. S27.**
Results of *BPP* species delimitation with all USCO data (left tree) and reduced USCO data* (right tree; gaps (and ambiguous data, A1 and A2) removed) (*with an *ASTRAL* guide tree inferred from these data) using different prior combinations for tau and theta for *Discoglossus* and *Chrysis*, mapped onto respective *ASTRAL* tree from A1–7, compared to morphospecies assignments (shown by symbols associated to each individual terminal).

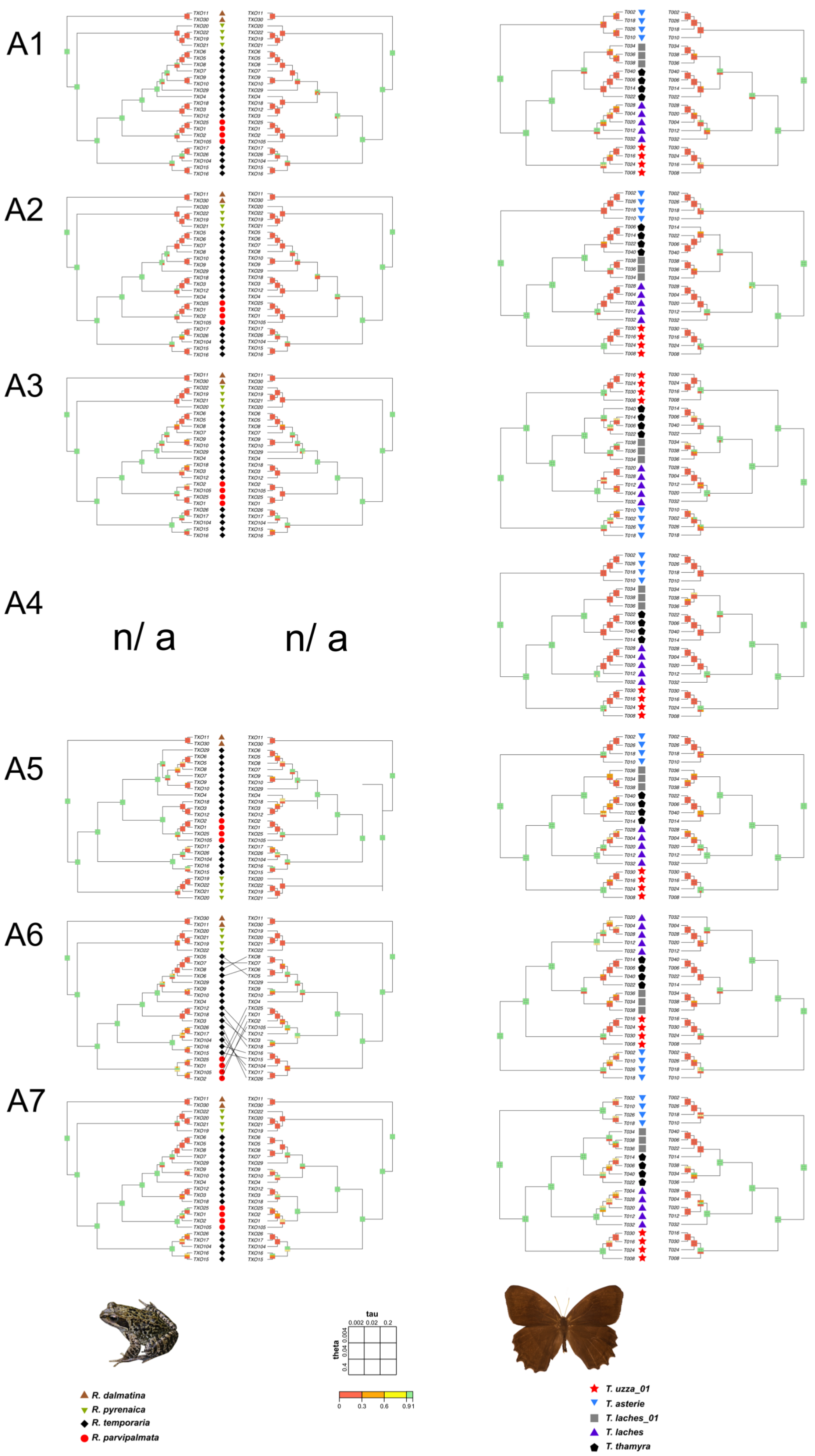


**Fig. S28.**
Results of *BPP* species delimitation with all USCO data (left tree) and reduced USCO data* (right tree; gaps (and ambiguous data, A1 and A2) removed) (*with an *ASTRAL* guide tree inferred from these data) using different prior combinations for tau and theta for *Taygetis*, and *Rana*, mapped onto respective *ASTRAL* tree from A1–7, compared to morphospecies assignments (shown by symbols associated to each individual terminal).

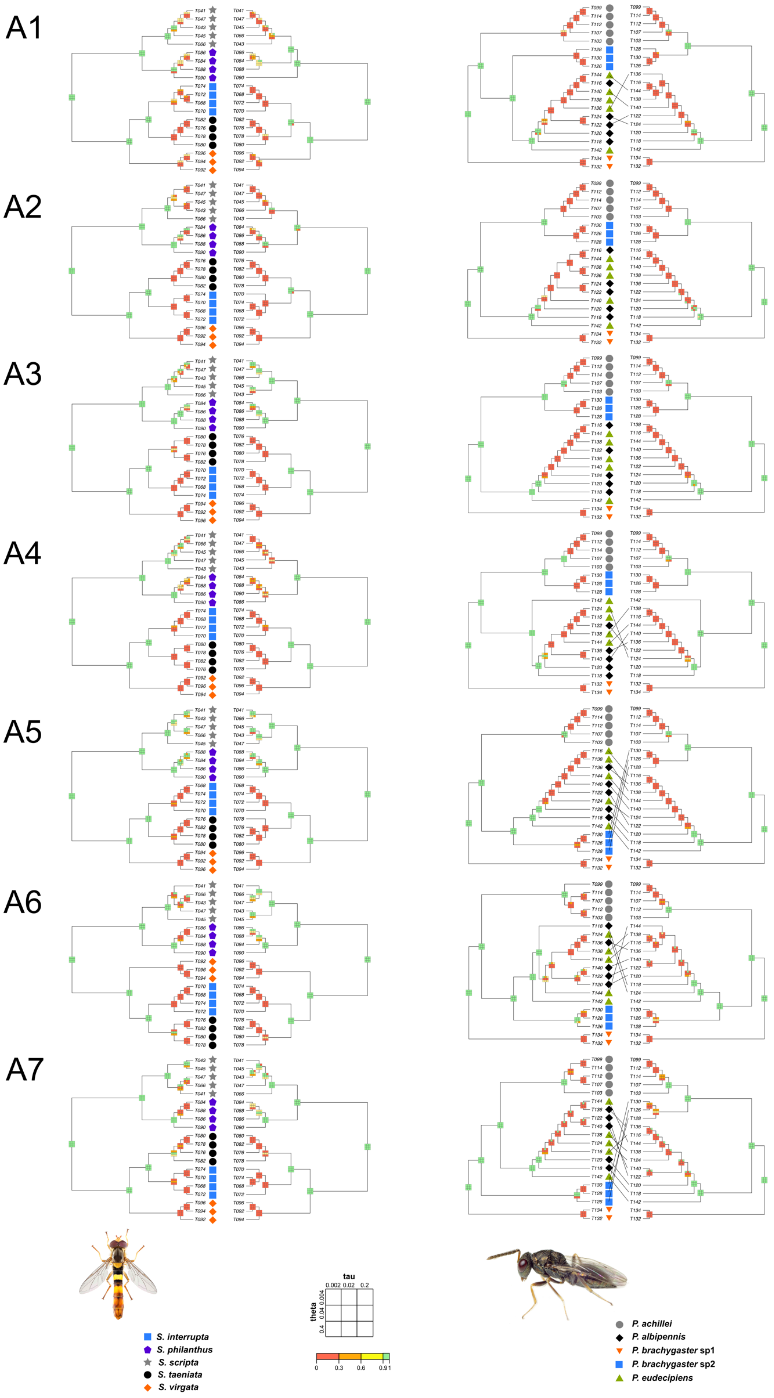


**Fig. S29.**
Results of *BPP* species delimitation with all USCO data (left tree) and reduced USCO data* (right tree; gaps (and ambiguous data, A1 and A2) removed) (*with an *ASTRAL* guide tree inferred from these data) using different prior combinations for tau and theta for *Pteromalus* and *Sphaerophoria*, mapped onto respective *ASTRAL* tree from A1–7, compared to morphospecies assignments (shown by symbols associated to each individual terminal).

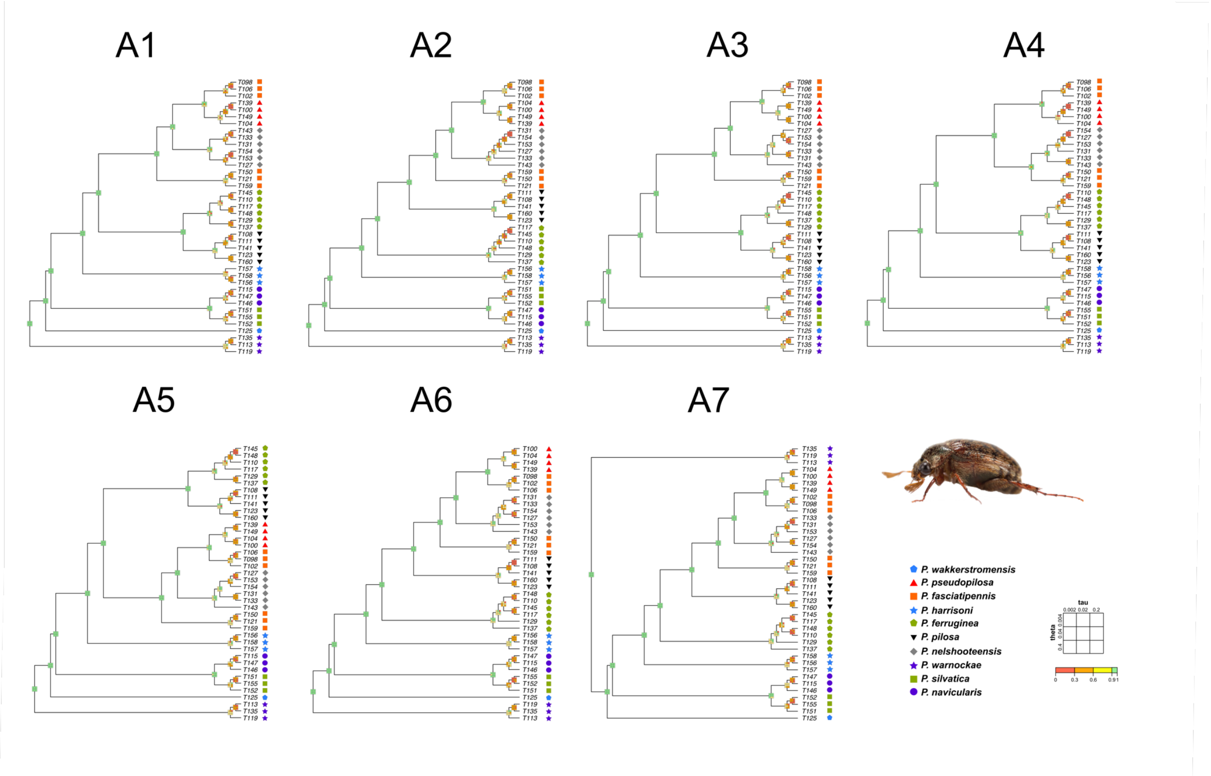


**Fig. S30.**
Results of *BPP* species delimitation of analysis without any data using different prior combinations for tau and theta for *Pleophylla*, mapped onto respective *ASTRAL* tree from A1–7, compared to morphospecies assignments (shown by symbols associated to each individual terminal).

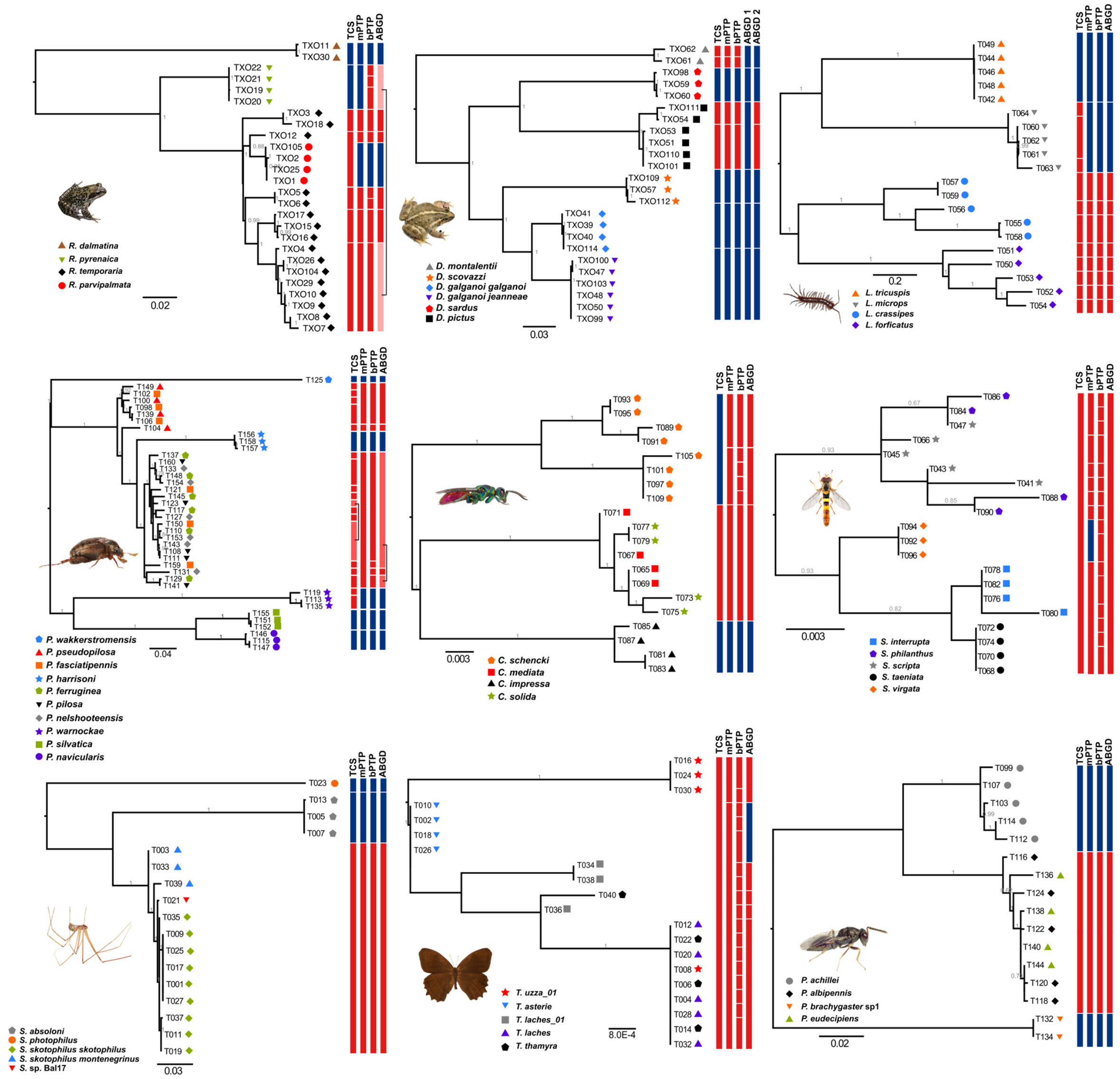

**Fig. S31.** Results of *COI* benchmarking and *COI*-based species delimitation with *mPTP*, *bPTP*, *TCS*, and *ABGD* for each study case (symbols associated to each terminal refer to morphospecies assignment), mapped onto the ML tree. Columns showing species entities colored as in Fig. S3.

**Supplementary Tables:**

**Table S1.**
**List of sampled specimens of this study**, including their collection data, sample and voucher number, NCBI Accession numbers for USCOs and COI data, as well as depository.

***(see separate excel sheet)***

**Table S2.**

**Evaluation of performance of the different assembly approaches (A1-A7),** in terms of number of retrieved USCOs, exons, and base pairs, alignment completeness (%), the number of USCOs and exons found in all specimens, as well as the amount of paralogs identified in at least one or at least 50% of the specimens.

| **Criteria of performance evaluation** | "BWA mapping" | Combined Trinity+ Orthograph+BWA | Orthograph+ hmmalign | IBA | Hybpiper | Phyluce | Orthograph+ Phyluce |
|---|---|---|---|---|---|---|---|
| | **A1** | **A2** | **A3** | **A4** | **A5** | **A6** | **A7** |
| **Dataset 1: *Pleophylla* (Coleoptera)** | | | | | | | |
| No. USCOs | 398 | **932** | 870 | 390 | 734 | 447 | 562 |
| No. Exons | 687 | na | na | 651 | na | na | na |
| No. Base pairs | 215827 | 446501 | 410658 | 171276 | 431106 | 372467 | 428693 |
| Alignment completeness (%) | 72.93 | 73.01 | 61.98 | 56.09 | 72.65 | 53.92 | 52.31 |
| No. USCOs in all specimens | 218 | **599** | 400 | 115 | 288 | 47 | 62 |
| No. Exons in all specimens | 398 | na | na | 166 | na | na | na |
| Paralogs in at least one specimen | | | | 14 | 119 | 21 | 44 |
| Paralogs in at least 50 % of the specimens | | | | 0 | 10 | 0 | 0 |
| **Dataset 2: *Lithobius* (Myriapoda)** | | | | | | | |
| No. USCOs | 142 | **716** | 713 | 256 | 691 | 88 | 367 |
| No. Exons | 205 | na | na | 366 | na | na | na |
| No. Base pairs | 38093 | 194579 | 198474 | 77700 | 246546 | 46191 | 150411 |
| Alignment completeness (%) | 44.1 | 30.87 | 55.38 | 43.5 | 51.89 | 37.59 | 37.65 |
| No. USCOs in all specimens | 18 | 28 | **262** | 25 | 126 | 8 | 21 |
| No. Exons in all specimens | 27 | na | na | 37 | na | na | na |
| Paralogs in at least one specimen | | | | 1 | 38 | 2 | 11 |
| Paralogs in at least 50 % of the specimens | | | | 0 | 0 | 0 | 0 |
| **Dataset 3: *Chrysis* (Hymenoptera)** | | | | | | | |
| No. USCOs | 636 | **964** | 960 | 595 | 910 | 617 | 551 |
| No. Exons | 1468 | na | na | 1331 | na | na | na |
| No. Base pairs | 427451 | 511914 | 494955 | 334109 | 622605 | 802214 | 576901 |
| Alignment completeness (%) | 95.52 | 88.37 | 78.72 | 90.63 | 91.47 | 82.09 | 81.92 |
| No. USCOs in all specimens | 632 | **878** | 820 | 540 | 793 | 336 | 317 |
| No. Exons in all specimens | 1484 | na | na | 1166 | na | na | na |
| Paralogs in at least one specimen | | | | 1 | 28 | 8 | 6 |
| Paralogs in at least 50 % of the specimens | | | | 0 | 1 | 0 | 0 |
| **Dataset 4: *Sphaerophoria* (Diptera)** | | | | | | | |
| No. USCOs | 209 | **940** | 888 | 285 | 752 | 445 | 623 |
| No. Exons | 241 | na | na | 363 | na | na | na |
| No. Base pairs | 73563 | 459128 | 405081 | 109438 | 418398 | 327396 | 446245 |
| Alignment completeness (%) | 68.37 | 71.96 | 67.35 | 44.92 | 74.73 | 65.3 | 60.78 |
| No. USCOs in all specimens | 94 | **638** | 533 | 73 | 421 | 91 | 137 |
| No. Exons in all specimens | 111 | na | na | 81 | na | na | na |
| Paralogs in at least one specimen | | | | 1 | 136 | 16 | 36 |
| Paralogs in at least 50 % of the specimens | | | | 0 | 8 | 0 | 2 |
| **Dataset 5: *Taygetis* (Lepidoptera)** | | | | | | | |
| No. USCOs | 172 | 832 | **848** | 446 | 766 | 139 | 417 |
| No. Exons | 210 | na | na | 714 | na | na | na |
| No. Base pairs | 43058 | 273940 | 257820 | 148820 | 309558 | 78607 | 221919 |
| Alignment completeness (%) | 65.47 | 80.86 | 71.7 | 71.84 | 78.79 | 57.95 | 67.23 |
| No. USCOs in all specimens | 75 | **660** | 564 | 244 | 514 | 36 | 140 |
| No. Exons in all specimens | 86 | na | na | 307 | na | na | na |
| Paralogs in at least one specimen | | | | 3 | 15 | 1 | 15 |
| Paralogs in at least 50 % of the specimens | | | | 1 | 0 | 0 | 3 |

**Table S2.** Continued.

| Criteria of performance evaluation | "BWA mapping" | Combined Trinity+ Orthograph+BWA | Orthograph+ hmmalign | IBA | Hybpiper | Phyluce | Orthograph+ Phyluce |
|---|---|---|---|---|---|---|---|
| | **A1** | **A2** | **A3** | **A4** | **A5** | **A6** | **A7** |
| **Dataset 6: *Stygopholcus* (Arachnida)** | | | | | | | |
| No. USCOs | 74 | **771** | 666 | 102 | 557 | 103 | 379 |
| No. Exons | 78 | na | na | 114 | na | na | na |
| No. Base pairs | 18177 | 210781 | 174678 | 27217 | 148029 | 56937 | 177861 |
| Alignment completeness (%) | 55.64 | 65.19 | 65.43 | 55.6 | 74.57 | 65.98 | 63.75 |
| No. USCOs in all specimens | 15 | **373** | 281 | 13 | 263 | 22 | 82 |
| No. Exons in all specimens | 18 | na | na | 14 | na | na | na |
| Paralogs in at least one specimen | | | | 0 | 4 | 1 | 6 |
| Paralogs in at least 50 % of the specimens | | | | 0 | 1 | 0 | 0 |
| **Dataset 7: *Pteromalus* (Hymenoptera)** | | | | | | | |
| No. USCOs | 758 | **955** | 945 | 701 | 919 | 645 | 568 |
| No. Exons | 1670 | na | na | 1525 | na | na | na |
| No. Base pairs | 533049 | 508332 | 476835 | 394880 | 639690 | 834067 | 586031 |
| Alignment completeness (%) | 90.24 | 87.22 | 81.07 | 84.61 | 91.8 | 78.93 | 75.62 |
| No. USCOs in all specimens | 722 | **875** | 800 | 602 | 750 | 315 | 261 |
| No. Exons in all specimens | 1545 | na | na | 1184 | na | na | na |
| Paralogs in at least one specimen | | | | 26 | 32 | 33 | 24 |
| Paralogs in at least 50 % of the specimens | | | | 3 | 8 | 2 | 2 |
| **Dataset 8: *Rana* (Anura)** | | | | | | | |
| No. USCOs | **767** | 746 | 739 | n/a | 658 | 165 | 271 |
| No. Exons | na | na | na | | na | na | na |
| No. Base pairs | 459224 | 309125 | 283215 | | 368706 | 182566 | 268545 |
| Alignment completeness (%) | 86.93 | 90.02 | 62.99 | | 69.58 | 37.37 | 46.22 |
| No. USCOs in all specimens | 650 | **668** | 539 | | 318 | 14 | 26 |
| No. Exons in all specimens | na | na | na | | na | na | na |
| Paralogs in at least one specimen | | | | | 46 | 22 | 29 |
| Paralogs in at least 50 % of the specimens | | | | | 23 | 1 | 1 |
| **Dataset 9: *Discoglossus* (Anura)** | | | | | | | |
| No. USCOs | **836** | 691 | 688 | n/a | 574 | 92 | 256 |
| No. Exons | na | na | na | | na | na | na |
| No. Base pairs | 422411 | 233569 | 226320 | | 259554 | 60923 | 157723 |
| Alignment completeness (%) | 69.65 | 79.71 | 59.32 | | 56.13 | 46.53 | 61.03 |
| No. USCOs in all specimens | **429** | 409 | 362 | | 82 | 9 | 32 |
| No. Exons in all specimens | na | na | na | | na | na | na |
| Paralogs in at least one specimen | | | | | 5 | 4 | 18 |
| Paralogs in at least 50 % of the specimens | | | | | 0 | 0 | 1 |

**Table S3.**
**References sequences (species) for bait design and USCO assembly** (i.e., multiple nucleotide sequence alignment cutting at exon boundaries). Transcriptomic (T) and genomic (G) origin is indicated in parentheses. EC: species used for exon cutting but not as sequence reference for bait design. For *Pteromalus*, the *Nasonia vitripennis* genome was used for both bait design and exon cutting.

| Genus | Sequence reference species and version where applicable | source |
|---|---|---|
| *Chrysis* | *Chrysis terminata* (T) | NCBI BioProject PRJNA252335 |
| *Chrysis* | *Chrysis indigotea* (T) | NCBI BioProject PRJNA252126 |
| *Chrysis* | EC: *Apis mellifera* (G) v. 4.5 | ftp://ftp.ensemblgenomes.org/pub/metazoa/release-39 |
| *Lithobius* | *Lithobius forficatus* (T) v. 1.1 | NCBI BioProject PRJNA254277 |
| *Lithobius* | *Eupolybothrus cavernicolus* (T) | NCBI BioProject PRJEB4548 |
| *Lithobius* | EC: *Strigamia maritima* (G) | ftp://ftp.ensemblgenomes.org/pub/metazoa/release-41 |
| *Pleophylla* | *Pleophylla* sp. (T) | NCBI BioProject PRJNA714151 |
| *Pleophylla* | *Trochalus* sp. (T) | NCBI BioProject PRJNA286594 |
| *Pleophylla* | EC: *Tribolium castaneum* (G) v. 5.2 | ftp://ftp.ensemblgenomes.org/pub/metazoa/release-37/fasta/tribolium_castaneum/cds/ |
| *Pteromalus* | *Nasonia vitripennis* (G) v. 2.1 | ftp://ftp.ensemblgenomes.org/pub/metazoa/release-39/ |
| *Pteromalus* | *Lariophagus distinguendus* (T) | NCBI BioProject PRJNA252174 |
| *Stygopholcus* | *Hoplopholcus cecconii* (T) | NCBI BioProject PRJNA275709 |
| *Stygopholcus* | *Smeringopus pallidus* (T) | NCBI BioProject PRJNA275722 |
| *Stygopholcus* | *Loxosceles reclusa* (G) v. 0.5.3 | https://i5k.nal.usda.gov/data/Arthropoda/loxrec-(Loxosceles_reclusa)/BCM-After-Atlas/ |
| *Sphaerophoria* | *Leucozona lucorum* (T) | NCBI BioProject PRJNA267954 |
| *Sphaerophoria* | *Episyrphus balteatus* (T) | NCBI BioProject PRJNA267940 |
| *Sphaerophoria* | EC: *Drosophila melanogaster* (G) v. 40 | ftp://ftp.ensemblgenomes.org/pub/metazoa/release-40/fasta/drosophila_melanogaster/dna/ |
| *Taygetis* | *Bicyclus anynana* (T) | NCBI BioProject PRJNA267867 |
| *Taygetis* | *Pararge aegeria* (T) | NCBI BioProject PRJNA267891 |
| *Taygetis* | EC: *Danaus plexippus* (G) v. 3.0 | http://download.lepbase.org/v4/sequence/ |
| *Discoglossus* | *Discoglossus galganoi* (T) | NCBI BioProject PRJNA542138 |
| *Rana* | *Rana parvipalmata* (T) | NCBI BioProject PRJNA542138 |

**Table S4.**
**Summary of characteristics of complete and reduced USCO data sets of the different assembly approaches (A1-A7),** in terms of the number of parsimony informative sites, the number of USCOs without gaps, the number of exons without gaps, the number of base pairs without gaps, the number of base pairs without ambiguities, the number of parsimony informative sites without gaps, and the number of parsimony informative sites without ambiguities.

| Criteria of performance evaluation | "BWA mapping" | Combined Trinity+ Orthograph+BWA | Orthograph+ hmmalign | IBA | Hybpiper | Phyluce | Orthograph+ Phyluce |
|---|---|---|---|---|---|---|---|
| | **A1** | **A2** | **A3** | **A4** | **A5** | **A6** | **A7** |
| **Dataset 1:** ***Pleophylla*** **(Coleoptera)** | | | | | | | |
| No. USCOs | 398 | **932** | 870 | 390 | 734 | 447 | 562 |
| No. Exons | 687 | na | na | 651 | na | na | na |
| No. Base pairs | 215827 | 446501 | 410658 | 171276 | 431106 | 372467 | 428693 |
| No. Pars. Inf. Sites | 7237 | 14781 | 21099 | 5405 | 24744 | 18615 | 21287 |
| No. USCOs without gaps | 215 | **584** | 266 | 115 | 282 | 47 | 62 |
| No. Exons without gaps | 396 | na | na | 166 | na | na | na |
| No. Base pairs without gaps | 89375 | 180674 | 73059 | 32336 | 99942 | 26964 | 32945 |
| No. Base pairs without ambiguities | 83061 | 165474 | na | na | na | na | na |
| No. Pars. Inf. Sites without gaps | 3175 | 6495 | 3694 | 1479 | 4473 | 1459 | 1845 |
| No. Pars. Inf. Sites without ambiguities | 1486 | 2763 | na | na | na | na | na |
| **Dataset 2:** ***Lithobius*** **(Myriapoda)** | | | | | | | |
| No. USCOs | 142 | **716** | 713 | 256 | 691 | 88 | 367 |
| No. Exons | 205 | na | na | 366 | na | na | na |
| No. Base pairs | 38093 | 194579 | 198474 | 77700 | 246546 | 46191 | 150411 |
| No. Pars. Inf. Sites | 1634 | 4947 | 35428 | 9478 | 45501 | 5200 | 16170 |
| No. USCOs without gaps | 17 | 24 | **148** | 25 | 115 | 8 | 21 |
| No. Exons without gaps | 26 | na | na | 37 | na | na | na |
| No. Base pairs without gaps | 4153 | 3719 | 27288 | 6959 | 22314 | 2635 | 6133 |
| No. Base pairs without ambiguities | 4090 | 3626 | na | na | na | na | na |
| No. Pars. Inf. Sites without gaps | 346 | 417 | 6936 | 1324 | 5228 | 707 | 1572 |
| No. Pars. Inf. Sites without ambiguities | 310 | 363 | na | na | na | na | na |
| **Dataset 3:** ***Chrysis*** **(Hymenoptera)** | | | | | | | |
| No. USCOs | 636 | **964** | 960 | 595 | 910 | 617 | 551 |
| No. Exons | 1468 | na | na | 1331 | na | na | na |
| No. Base pairs | 427451 | 511914 | 494955 | 334109 | 622605 | 802214 | 576901 |
| No. Pars. Inf. Sites | 5112 | 6191 | 8198 | 4877 | 12747 | 18801 | 12792 |
| No. USCOs without gaps | 632 | **865** | 740 | 540 | 793 | 336 | 317 |
| No. Exons without gaps | 1454 | na | na | 1166 | na | na | na |
| No. Base pairs without gaps | 371020 | 365242 | 247701 | 260848 | 428133 | 374366 | 274512 |
| No. Base pairs without ambiguities | 363488 | 357472 | na | na | na | na | na |
| No. Pars. Inf. Sites without gaps | 4476 | 4498 | 4121 | 4052 | 7221 | 9998 | 7059 |
| No. Pars. Inf. Sites without ambiguities | 1573 | 1565 | na | na | na | na | na |

**Table S4.** continued.

| Criteria of performance evaluation | "BWA mapping" | Combined Trinity+ Orthograph+BWA | Orthograph+ hmmalign | IBA | Hybpiper | Phyluce | Orthograph+ Phyluce |
|---|---|---|---|---|---|---|---|
| | **A1** | **A2** | **A3** | **A4** | **A5** | **A6** | **A7** |
| **Dataset 4: *Sphaerophoria* (Diptera)** | | | | | | | |
| No. USCOs | 209 | **940** | 888 | 285 | 752 | 445 | 623 |
| No. Exons | 241 | na | na | 363 | na | na | na |
| No. Base pairs | 73563 | 459128 | 405081 | 109438 | 418398 | 327396 | 446245 |
| No. Pars. Inf. Sites | 1661 | 8914 | 13023 | 1613 | 16721 | 12255 | 15103 |
| No. USCOs without gaps | 92 | **627** | 410 | 73 | 418 | 91 | 137 |
| No. Exons without gaps | 107 | na | na | 81 | na | na | na |
| No. Base pairs without gaps | 21024 | 181822 | 107529 | 15979 | 134268 | 52579 | 70956 |
| No. Base pairs without ambiguities | 20047 | 165149 | na | na | na | na | na |
| No. Pars. Inf. Sites without gaps | 375 | 3409 | 3954 | 459 | 4545 | 2359 | 3113 |
| No. Pars. Inf. Sites without ambiguities | 223 | 623 | na | na | na | na | na |
| **Dataset 5: *Taygetis* (Lepidoptera)** | | | | | | | |
| No. USCOs | 172 | 832 | **848** | 446 | 766 | 139 | 417 |
| No. Exons | 210 | na | na | 714 | na | na | na |
| No. Base pairs | 43058 | 273940 | 257820 | 148820 | 309558 | 78607 | 221919 |
| No. Pars. Inf. Sites | 773 | 3490 | 5383 | 2310 | 11346 | 1247 | 5544 |
| No. USCOs without gaps | 75 | **653** | 404 | 244 | 492 | 36 | 140 |
| No. Exons without gaps | 86 | na | na | 307 | na | na | na |
| No. Base pairs without gaps | 15327 | 169613 | 87705 | 58763 | 126321 | 16694 | 55974 |
| No. Base pairs without ambiguities | 15026 | 163746 | na | na | na | na | na |
| No. Pars. Inf. Sites without gaps | 223 | 2341 | 1774 | 1149 | 3075 | 359 | 1679 |
| No. Pars. Inf. Sites without ambiguities | 156 | 978 | na | na | na | na | na |
| **Dataset 6: *Stygopholcus* (Arachnida)** | | | | | | | |
| No. USCOs | 74 | **771** | 666 | 102 | 557 | 103 | 379 |
| No. Exons | 78 | na | na | 114 | na | na | na |
| No. Base pairs | 18177 | 210781 | 174678 | 27217 | 148029 | 56937 | 177861 |
| No. Pars. Inf. Sites | 492 | 6742 | 7443 | 786 | 7154 | 3045 | 11426 |
| No. USCOs without gaps | 13 | **360** | 213 | 13 | 262 | 22 | 82 |
| No. Exons without gaps | 16 | na | na | 14 | na | na | na |
| No. Base pairs without gaps | 1937 | 67768 | 39612 | 2449 | 49155 | 7140 | 23982 |
| No. Base pairs without ambiguities | 1903 | 66016 | na | na | na | na | na |
| No. Pars. Inf. Sites without gaps | 67 | 3165 | 2067 | 108 | 2499 | 457 | 1864 |
| No. Pars. Inf. Sites without ambiguities | 56 | 2610 | na | na | na | na | na |

**Table S4.** continued.

| Criteria of performance evaluation | "BWA mapping" | Combined Trinity+ Orthograph+BWA | Orthograph+ hmmalign | IBA | Hybpiper | Phyluce | Orthograph+ Phyluce |
|---|---|---|---|---|---|---|---|
| | **A1** | **A2** | **A3** | **A4** | **A5** | **A6** | **A7** |
| **Dataset 7: *Pteromalus* (Hymenoptera)** | | | | | | | |
| No. USCOs | 758 | **955** | 945 | 701 | 919 | 645 | 568 |
| No. Exons | 1670 | na | na | 1525 | na | na | na |
| No. Base pairs | 533049 | 508332 | 476835 | 394880 | 639690 | 834067 | 586031 |
| No. Pars. Inf. Sites | 7985 | 6901 | 10325 | 6570 | 15307 | 19967 | 13875 |
| No. USCOs without gaps | 720 | **869** | 736 | 602 | 749 | 314 | 261 |
| No. Exons without gaps | 1542 | na | na | 1184 | na | na | na |
| No. Base pairs without gaps | 407909 | 373652 | 243819 | 259453 | 412683 | 333626 | 211707 |
| No. Base pairs without ambiguities | 397881 | 362376 | na | na | na | na | na |
| No. Pars. Inf. Sites without gaps | 5737 | 5230 | 5050 | 4751 | 8025 | 5227 | 5824 |
| No. Pars. Inf. Sites without ambiguities | 4031 | 3203 | na | na | na | na | na |
| **Dataset 8: *Rana* (Anura)** | | | | | | | |
| No. USCOs | **767** | 746 | 739 | na | 658 | 165 | 271 |
| No. Exons | na | na | na | | na | na | na |
| No. Base pairs | 459224 | 309125 | 283215 | | 368706 | 182566 | 268545 |
| No. Pars. Inf. Sites | 7390 | 5361 | 6988 | | 22457 | 6138 | 10536 |
| No. USCOs without gaps | 640 | **663** | 332 | | 303 | 14 | 26 |
| No. Exons without gaps | na | na | na | | na | na | na |
| No. Base pairs without gaps | 284517 | 228101 | 73674 | | 97899 | 13868 | 25317 |
| No. Base pairs without ambiguities | 274019 | 218840 | na | na | na | na | na |
| No. Pars. Inf. Sites without gaps | 4701 | 3875 | 1967 | | 2985 | 598 | 1051 |
| No. Pars. Inf. Sites without ambiguities | 2337 | 1852 | na | na | na | na | na |
| **Dataset 9: *Discoglossus* (Anura)** | | | | | | | |
| No. USCOs | **836** | 691 | 688 | na | 574 | 92 | 256 |
| No. Exons | na | na | na | | na | na | na |
| No. Base pairs | 422411 | 233569 | 226320 | | 259554 | 60923 | 157723 |
| No. Pars. Inf. Sites | 7491 | 4912 | 5284 | | 16982 | 1670 | 8367 |
| No. USCOs without gaps | **400** | 391 | 180 | | 65 | 9 | 32 |
| No. Exons without gaps | na | na | na | | na | na | na |
| No. Base pairs without gaps | 97522 | 91653 | 36729 | | 13920 | 5488 | 13112 |
| No. Base pairs without ambiguities | 94160 | 88620 | na | na | na | na | na |
| No. Pars. Inf. Sites without gaps | 2175 | 2115 | 1160 | | 433 | 194 | 770 |
| No. Pars. Inf. Sites without ambiguities | 1341 | 1354 | na | na | na | na | na |

**Table S5.**
**Syntopical occurrence of the investigated species for each study case,** as far as known, given either as presence/ absence data (1/0), or as number of sites with syntopical occurrence (n).

| *Stygopholcus* | *St.absoloni* | *St. photophilus* | *St. skotophilus skotophilus* | *St. skotophilus montenegrinus* | *St.* sp. Bal17 |
|---|---|---|---|---|---|
| *St. absoloni* | x | | | | |
| *St. photophilus* | 0 | x | | | |
| *St. skotophilus skotophilus* | 0 | 0 | x | | |
| *St. skotophilus montenegrinus* | 0 | 0 | 0 | x | |
| *St.* sp. Bal17 | 0 | 0 | 0 | 0 | x |

| *Sphaerophoria* | *Sph. interrupta* | *Sph. philantha* | *Sph. scripta* | *Sph. taeniata* | *Sph. virgata* |
|---|---|---|---|---|---|
| *Sph. interrupta* | x | | | | |
| *Sph. philantha* | 22 | x | | | |
| *Sph. scripta* | >120 | 46 | x | | |
| *Sph. taeniata* | 67 | 26 | >120 | x | |
| *Sph. virgata* | 33 | 25 | 56 | 28 | x |

| *Pleophylla* | *P. fasciatipennis* | *P. ferruginea* | *P. harrisoni* | *P. navicularis* | *P. nelshoogteensis* | *P. pilosa* | *P. pseudopilosa* | *P. silvatica* | *P. wakkerstromensis* | *P. warnockae* |
|---|---|---|---|---|---|---|---|---|---|---|
| *P. fasciatipennis* | x | | | | | | | | | |
| *P. ferruginea* | 16 | x | | | | | | | | |
| *P. harrisoni* | - | | x | | | | | | | |
| *P. navicularis* | 7 | 13 | - | x | | | | | | |
| *P. nelshoogteensis* | - | - | - | - | x | | | | | |
| *P. pilosa* | 5 | 11 | - | 6 | - | x | | | | |
| *P. pseudopilosa* | 1 | - | - | - | - | - | x | | | |
| *P. silvatica* | 2 | 4 | - | 5 | 6 | 4 | - | x | | |
| *P. wakkerstromensis* | - | 1 | - | - | - | - | - | - | x | |
| *P. warnockae* | 1 | - | - | - | - | - | - | - | - | x |

| *Chrysis* | *C. impressa* | *C. mediata* | *C. schencki* | *C. solida* |
|---|---|---|---|---|
| *C. impressa* | x | | | |
| *C. mediata* | ? | x | | |
| *C. schencki* | 1 | ? | x | |
| *C. solida* | 1 | ? | ? | x |

| *Taygetis* | *T. asterie* | *T. laches* | *T. laches_01* | *T. thamyra* | *T. uzza_01* |
|---|---|---|---|---|---|
| *T. asterie* | x | | | | |
| *T. laches* | 0 | x | | | |
| *T. laches_01* | 0 | 1 | x | | |
| *T. thamyra* | 0 | 1 | 1 | x | |
| *T. uzza_01* | 0 | 0 | 0 | 0 | x |

**Table S5.** Continued.

| *Pteromalus* | *P. achillei* | *P. albipennis* | *P. brachygaster* sp1 | *P. brachygaster* sp2 | *P. eudecipiens* |
|---|---|---|---|---|---|
| ***P. achillei*** | x | | | | |
| ***P. albipennis*** | 1 | x | | | |
| ***P. brachygaster* sp1** | ? | ? | x | | |
| ***P. brachygaster* sp2** | 1 | 1 | 1 | x | |
| ***P. eudecipiens*** | ? | ? | ? | ? | x |

| *Lithobius* | *L. tricuspis* | *L. forficatus* | *L. crassipes* | *L. microps* |
|---|---|---|---|---|
| ***L. tricuspis*** | x | | | |
| ***L. forficatus*** | 0 | x | | |
| ***L. crassipes*** | 0 | 1 | x | |
| ***L. microps*** | 0 | 1 | 1 | x |

| *Discoglossus* | *D. galganoi galganoi* | *D. galganoi jeanneae* | *D. montalentii* | *D. pictus* | *D. sardus* | *D. scovazzi* |
|---|---|---|---|---|---|---|
| ***D. galganoi galganoi*** | x | | | | | |
| ***D. galganoi jeanneae*** | 0 | x | | | | |
| ***D. montalentii*** | 0 | 0 | x | | | |
| ***D. pictus*** | 0 | 0 | 0 | x | | |
| ***D. sardus*** | 0 | 0 | 1 | 0 | x | |
| ***D. scovazzi*** | 0 | 0 | 0 | 0 | 0 | x |

| *Rana* | *R. dalmatina* | *R. parvipalmata* | *R. pyrenaica* | *R. temporaria* |
|---|---|---|---|---|
| ***R. dalmatina*** | x | | | |
| ***R. parvipalmata*** | 0 | x | | |
| ***R. pyrenaica*** | 0 | 0 | x | |
| ***R. temporaria*** | 1 | 0 | 1 | x |

**Table S6.**
**Accession numbers of *COI* sequences used to infer complete amino acid consensus *COI* sequences of *Discoglossus* and Ranidae.** These sequences have been aligned with mafft-linsi v7.475 and consensus sequences have been determined with an in-house script and a 50% consensus threshold.

| **Accession numbers (*Discoglossus*)** |
| --- |
| YP_192929.1, AAS83383.1 |
| ***Consensus sequence(Discoglossus)*** |
| MAITRWLFSTNHKDIGTLYLIFGAWAGMVGTALSLLIRAELSQPGTLLGDDQIYNVIVTAHAFVMIFFMVMPIMIGGFGNWLIPLMIGAPDMAFPRMNNMSFW LLPPSFLLLLASSGVEAGAGTGWTVYPPLAGNLAHAGASVDLTIFSLHLAGVSSISGAINFITTTINMKPPSMSQYQTPLFVWSVLITAVLLLLSLPVLAAGITMLLTDR NLNTTFFDPAGGGDPVLYQHLFWFFGHPEVYILILPGFGMISHIVTYYSGKKEPFGYMGMVWAMMSIGLLGFIVWAHHMFTVDLNVDTRAYFTSATMIIAIPTGV KVFSWLATMHGGTIKWDAAMLWALGFIFLFTVGGLTGIVLANSSLDIVLHDTYYVVAHFHYVLSMGAVFAIMGGFVHWFPLFTGYTLHETWTKIHFGVMFAGV NLTFFPQHFLGLAGMPRRYSDYPDAYTLWNTVSSIGSLVSLVAVIMMMFIIWEAFSAKREVILTELTMTNVEWLHGCPPPYHTFEEPAFVQMPYRA |

| **Accession numbers (Ranidae)** |
| --- |
| AAK56868.1, AAK56871.1, AAK56874.1, AAK56877.1, AAK56880.1, AAK56883.1, AAK56886.1, AAK56889.1, AAK56892.1, AAK56895.1, AAK56898.1, AAK56901.1, AAK56904.1, AAK56907.1, AAK56910.1, AAT09568.1, AAT09571.1, AAT09574.1, AAT09577.1, AAT09580.1, AAT09583.1, AAT09586.1, AAT09589.1, AAT09592.1, AAT09595.1, AAT09598.1, AAT09601.1, AAT09604.1, ABF06460.1, ACB30320.1, AEC12164.1, AFL65900.1, AGN71258.1, AGS43912.1, AHG32624.1, AHG32637.1, AHG32650.1, AHG53814.1, AHH80770.1, AHH80783.1, AHL84091.1, AHZ87099.1, AIJ20081.1, AIP86855.1, AIP86856.1, AIP86857.1, AIP86858.1, AIP86860.1, AIP86862.1, AIP86863.1, AIP86865.1, AIP86867.1, AIP86868.1, AIP86869.1, AIP86871.1, AIP86873.1, AIP86875.1, AIP86877.1, AIP86878.1, AIP86879.1, AIP92349.1, AIQ78407.1, AIU38923.1, AIU44429.1, AIZ97050.1, AIZ97121.1, AJO99983.1, AJW75353.1, AJW75535.1, AKA55338.1, AKA55351.1, AKA55364.1, AKA55377.1, AKE36766.1, AKN10614.1, AKQ19770.1, AKQ19784.1, AKQ19796.1, AKQ19809.1, AKQ19822.1, AKQ19835.1, AKQ19848.1, ALJ78651.1, ALM54856.1, ALN94575.1, ALN94576.1, ALN94577.1, ALN94578.1, ALN94579.1, ALN94580.1, ALN94581.1, ALN94582.1, ALN94583.1, ALN94584.1, ALN94585.1, ALN94586.1, ALN94587.1, ALN94588.1, ALN94589.1, ALN94590.1, ALN94591.1, ALN94592.1, ALN94593.1, ALN94594.1, ALN94595.1, ALN94596.1, ALN94597.1, ALN94598.1, ALN94599.1, ALN94600.1, ALN94601.1, ALN94602.1, ALN94603.1, ALN94604.1, ALN94605.1, ALN94606.1, ALN94607.1, ALT66294.1, AMD09888.1, AMD09889.1, AMD09890.1, AMD09891.1, AMD09898.1, AMD09899.1, AMD09900.1, AMD09901.1, AMD09902.1, AMR74881.1, AMY15643.1, ANC62869.1, ANW37039.1, ANW37052.1, ANW37065.1, AOW69080.1, AOW70447.1, APB02945.1, APT37072.1, APT37073.1, APT37074.1, APT37075.1, APT37076.1, APT37077.1, APT37078.1, APT37079.1, APT37080.1, APT37081.1, APT37082.1, APT37083.1, APT37084.1, APT37085.1, APT37086.1, APT37087.1, APT37088.1, APT37089.1, APT37090.1, APT37091.1, APT37092.1, APT37093.1, APT37094.1, APT37095.1, APT37096.1, APT37097.1, APT37098.1, APT37099.1, APT37100.1, APT37101.1, APT37102.1, APT37103.1, APT37104.1, APT37105.1, APT37106.1, APT37107.1, APT37108.1, APT37109.1, APT37110.1, APT37111.1, APT37112.1, APT37113.1, APT37114.1, APT37115.1, APT37116.1, APT37117.1, APT37118.1, APT37119.1, APT37120.1, APT37121.1, APT37122.1, APT37123.1, APT37124.1, APT37125.1, APT37126.1, APT37127.1, APT37128.1, APT37129.1, APT37130.1, APT37131.1, APT37132.1, APT37133.1, APT37134.1, APT37135.1, APT37136.1, APT37137.1, APT37138.1, APT37139.1, APT37140.1, APT37141.1, APT37142.1, APT37143.1, APT37144.1, APT37145.1, APT37146.1, APT37147.1, APT37148.1, APT37149.1, APT37150.1, APT37151.1, APT37152.1, APT37153.1, APT37154.1, APT37155.1, APT37156.1, APT37157.1, APT37158.1, APT37159.1, APT37160.1, APT37161.1, APT37162.1, APT37163.1, APT37164.1, APT37165.1, APT37166.1, APT37167.1, APT37168.1, APT37169.1, APT37170.1, APT37171.1, APT37172.1, APT37173.1, APT37174.1, APY20713.1, ARO35535.1, ARO35548.1, ARO35560.1, ASO66798.1, ATE88449.1, ATE88450.1, ATE88451.1, ATY37523.1, ATY37524.1, ATY37525.1, ATY37526.1, ATY37527.1, ATY37528.1, ATY37529.1, ATY37530.1, ATY37531.1, ATY37532.1, AWE05761.1, AWE05762.1, AWE05763.1, AWE05764.1, AWE05765.1, AWE05766.1, AWE05767.1, AWE05768.1, AWE05769.1, AWE05770.1, AWE05771.1, AWE05772.1, AWE05773.1, AWE05774.1, AWE05775.1, AWE05776.1, AWE05777.1, AWE05778.1, AWE05779.1, AWE05780.1, AWE05781.1, AWE05782.1, AWE05783.1, AWE05784.1, AWE05785.1, AWE05786.1, AWE05787.1, AWE05788.1, AWE05789.1, AWE05790.1, AWE05791.1, AWE05804.1, AWE05805.1, AWE05806.1, AWE05807.1, AWE05808.1, AWE05809.1, AWE05810.1, AWE05811.1, AWE05812.1, AWE05813.1, AWE05814.1, AWE05815.1, AWE05816.1, AWE05817.1, AWE05818.1, AWE05819.1, AWE05820.1, AWE05821.1, AWE05822.1, AWE05886.1, AWE05887.1, AWE05888.1, AWE05889.1, AWE05890.1, AWE05891.1, AWE05892.1, AWE05893.1, AWE05894.1, AWE05895.1, AWE05896.1, AWE05897.1, AWE05898.1, AWW14132.1, AWW14133.1, AWW14134.1, AWW14135.1, AWW14136.1, AWW14137.1, AWW14138.1, AWW14139.1, AWW14140.1, AWW14141.1, AWW14142.1, AWW14143.1, AWW14144.1, AWW14145.1, AWW14146.1, AWW14147.1, AWW14148.1, AWW14149.1, AWW14150.1, AWW14151.1, AWW14152.1, AWW14153.1, AWW14154.1, AWW14155.1, AWW14156.1, AWW14157.1, AWW14158.1, AWW14159.1, AWW14160.1, AXE73273.1, AXE73274.1, AXE73275.1, AXE73276.1, AXE73277.1, AXE73279.1, AXE73281.1, AXE73282.1, AXE73287.1, AXE73288.1, AXE73289.1, AXE73290.1, AXE73291.1, AXE73292.1, AXE73293.1, AXE73294.1, AXE73295.1, AXE73296.1, AXE73297.1, AXE73298.1, AXE73299.1, AXE73300.1, AXE73302.1, NP_116769.1, QBA17834.1, QBK18851.1, QBK18852.1, QBK18853.1, QBK18854.1, QBK18855.1, QBK18856.1, QBK18857.1, QBK18858.1, QBK18859.1, QBK18860.1, QBK18861.1, QBK18862.1, QBK18863.1, QBK18864.1, QBK18865.1, QBK18866.1, QBK18867.1, QBK18868.1, QBK18869.1, QBK18870.1, QBK18871.1, QBK18872.1, QBK18873.1, QBK18874.1, QBK18875.1, QBK18876.1, QBK18877.1, QBK18878.1, QBK18879.1, QBK18880.1, QBK18881.1, QBK18882.1, QBK18883.1, QBK18884.1, QBK18885.1, QBK18886.1, QBK18887.1, QBK18888.1, QBK18889.1, QBK18890.1, QBK18891.1, QBK18894.1, QBK18895.1, QBK18896.1, QBK18897.1, QBK18898.1, QBK18899.1, QBK18900.1, QBK18901.1, QBK18902.1, QBK18903.1, QBK18904.1, QBK18905.1, QBK18906.1, QBK18907.1, QBK18908.1, QBK18909.1, QBK18910.1, QBK18911.1, QBK18912.1, QBK18913.1, QBK18914.1, QBK18915.1, QBK18916.1, QBK18917.1, QBK18918.1, QBK18919.1, QBK18920.1, QBK18921.1, QBK18922.1, QBK18923.1, QBK18924.1, QBK18925.1, QBK18926.1, QBK18927.1, QBK18928.1, QBK18929.1, QBK18930.1, QBK18931.1, QBK18932.1, QBK18933.1, QBK18934.1, QBK18935.1, |

QBK18936.1, QBK18937.1, QBK18938.1, QBK18939.1, QBK18940.1, QBK18941.1, QBK18942.1, QBK18943.1, QBK18944.1, QBK18945.1, QBK18946.1, QBK18947.1, QBK18948.1, QBK18949.1, QBK18950.1, QBK18951.1, QBK18952.1, QBK18953.1, QBK18954.1, QBK18955.1, QBK18956.1, QBK18957.1, QBK18958.1, QBK18959.1, QBK18960.1, QBK18961.1, QBK18962.1, QBK18963.1, QBK18964.1, QBK18965.1, QBK18966.1, QBK18967.1, QBK18968.1, QBK18969.1, QBK18970.1, QBK18971.1, QBK18972.1, QBK18973.1, QBK18974.1, QBK18975.1, QBK18976.1, QBK18977.1, QBK18978.1, QCA41672.1, QDH12198.1, QDJ94070.1, QDJ94071.1, QDJ94072.1, QEQ76300.1, YP_001122915.1, YP_001165458.1, YP_004327604.1, YP_004891264.1, YP_006883435.1, YP_008757900.1, YP_008816325.1, YP_008816338.1, YP_008816350.1, YP_009003745.1, YP_009003758.1, YP_009024571.1, YP_009034285.1, YP_009048386.1, YP_009049185.1, YP_009054495.1, YP_009072517.1, YP_009107310.1, YP_009107518.1, YP_009132519.1, YP_009132532.1, YP_009132545.1, YP_009132558.1, YP_009144128.1, YP_009164151.1, YP_009178062.1, YP_009178231.1, YP_009183154.1, YP_009228341.1, YP_009228354.1, YP_009228367.1, YP_009228380.1, YP_009228393.1, YP_009228406.1, YP_009229624.1, YP_009250821.1, YP_009254183.1, YP_009382742.1, YP_009382755.1, YP_009423971.1, YP_009423984.1, YP_009423997.1, YP_009631376.1, YP_009667252.1, YP_009701402.1,

***Consensus sequence (Ranidae)***

MMFTRWFFSTNHKDIGTLYLIFGAWAGMVGTALSLLIRAELSQPGTLLGDDQIYNVIVTAHAFVMIFFMVMPILIGGFGNWLVPLMIGAPDMAFPRMNNMSF
WLLPPSFFLLLASSTVEAGAGTGWTVYPPLAGNLAHAGPSVDLAIFSLHLAGVSSILGAINFITTIINMKPXSTTQYQTPLFVWSVLITAVLLLLSLPVLAAGITMLLTDR
NLNTTFFDPAGGGDPVLYQHLFWFFGHPEVYILILPGFGIISHVVAYYSNKKEPFGYMGMVWAMLSIGLLGFIVWAHHMFTTDLNVDTRAYFTSATMIIAIPTGVK
VFSWLATMHGGIIKWEAPMLWALGFIFLFTVGGLTGIVLANSSIDIVLHDTYYVVAHFHYVLSMGAVFAIMAGFVHWFPLFTGFTLHELWTKIHFVVMFTGVNLT
FFPQHFLGLAGMPRRYSDYPDAYTLWNTVSSVGSLISLVAVVMMMFIIWEAFAAKRLFXGXELTSTNIEWLLGFPPHYHTFEESTFSIKLTRE

**Table S7.**
**Primer names with their references for *COI* sequencing in the study cases of Arthropoda.**

| Taxa/ Individuals | Primer | Reference |
|---|---|---|
| *Chrysis* | LCO1490/ Cl-N-2191 (Nancy) | (*112, 113*) |
| *Lithobius* (T044, T046, T048, T052, T050, T051, T52, T062) | LCO1490/ Cl-N-2191 (Nancy) | (*112, 113*)) |
| *Lithobius* (T049, T061) | HCO2198-JJ2/ LCO1490-JJ2 | (*114*) |
| *Lithobius* (T058, T056, T060, T054, T059, T064, T058, T042, T055, T063) | HCO2198-JJ/ LCO1490-JJ | (*114*) |
| *Pleophylla* | C1-J-2183 (Jerry)/ TL2-N-3014 (Pat) | (*113*) |
| *Pteromalus* | HCO2198-JJ/ LCO1490-JJ | (*115*) |
| *Sphaerophoria* (T041, T043, T047, T078, T080, T088, T090) | COI-780-R (COI-Dipt-2183R)/ LCO1490-F | (*112*) |
| *Sphaerophoria* (T066, T068, T082, T084, T086, T092, T094) | HCO2198-JJ/ LCO1490-JJ | (*115*) |
| *Sphaerophoria* (T070, T076) | HCO2198/ LCO1490 | (*112*) |
| *Sphaerophoria* (T096, T072, T074, T045) | LCO1490/ C1-N2191 (Nancy) | (*112, 113*)) |
| *Stygopholcus* | HCO2198-JJ/ LCO1490-JJ | (*115*) |
| *Taygetis* | LCO_nym/ HCO_nym | (*116*) |
| *Taygetis* (T016, T022, T024, T030) | HCO2198-JJ/ LCO1490-JJ | (*115*) |
